# A fast-slow trait continuum at the level of entire communities

**DOI:** 10.1101/2023.07.12.548516

**Authors:** Margot Neyret, Gaëtane Le Provost, Andrea Larissa Boesing, Florian D. Schneider, Dennis Baulechner, Joana Bergmann, Franciska de Vries, Anna Maria Fiore-Donno, Stefan Geisen, Kezia Goldmann, Anna Merges, Ruslan A. Saifutdinov, Nadja K. Simons, Joseph A. Tobias, Andrey S. Zaitsev, Martin M. Gossner, Kirsten Jung, Ellen Kandeler, Jochen Krauss, Caterina Penone, Michael Schloter, Stefanie Schulz, Michael Staab, Volkmar Wolters, Antonios Apostolakis, Klaus Birkhofer, Steffen Boch, Runa S. Boeddinghaus, Ralph Bolliger, Michael Bonkowski, Francois Buscot, Kenneth Dumack, Markus Fischer, Huei Ying Gan, Johannes Heinze, Norbert Hölzel, Katharina John, Valentin H. Klaus, Till Kleinebecker, Sven Marhan, Jörg Müller, Swen C. Renner, Matthias Rillig, Noëlle V. Schenk, Ingo Schöning, Marion Schrumpf, Sebastian Seibold, Stephanie Socher, Emily F. Solly, Miriam Teuscher, Mark van Kleunen, Tesfaye Wubet, Pete Manning

**Affiliations:** Senckenberg Biodiversity and Climate Research Centre, Frankfurt, Germany; Laboratoire d’Écologie Alpine, CNRS, UGA, Grenoble, France; National Research Institute for Agriculture, Food and Environment, Bordeaux, France; ISOE - Institute for social-ecological research, Frankfurt am Main, Germany; Justus Liebig University, Department of Animal Ecology, Giessen, Germany; Leibniz Center for Agricultural Landscape Research (ZALF), Müncheberg, Germany; Institute for Biodiversity and Ecosystem Dynamics, University of Amsterdam, The Netherlands; Terrestrial Ecology, Institute of Zoology, University of Cologne, Köln, Germany; Laboratory of Nematology, Wageningen University and Research, Wageningen, The Netherlands; Helmholtz Centre for Environmental Research (UFZ), Soil Ecology Department, Leipzig, Germany; A.N. Severtsov Institute of Ecology and Evolution, Russian Academy of Sciences, Moscow, Russia; Ecological Networks, Technical University Darmstadt, Darmstadt, Germany; Department of Life Sciences, Imperial College London, Ascot, UK; Forest Entomology, Swiss Federal Research Institute WSL, Birmensdorf, Switzerland; Department of Environmental Systems Science, Institute of Terrestrial Ecosystems, ETH Zürich, Zürich, Switzerland; Universität Ulm, Ulm, Germany; Department of Soil Biology, Institute of Soil Science and Land Evaluation, University of Hohenheim, Stuttgart, Germany; Department of Animal Ecology and Tropical Biology, Biocenter, University of Würzburg, Würzburg, Germany; Institute of Plant Sciences, University of Bern, Bern, Switzerland; Helmholtz Zentrum Muenchen, Research Unit for Comparative Microbiome Analysis, Oberschleissheim, Germany; Chair for Soil Sciences, Technical University of Munich, Freising, Germany; Department of Biogeochemical Processes, Max-Planck-Institute for Biogeochemistry, Jena, Germany; Brandenburg University of Technology, Cottbus, Germany; Swiss Federal Research Institute WSL, Birmensdorf, Switzerland; Department Plant Production and Production Related Environmental Protection, Center for Agricultural Technology Augustenberg (LTZ), Karlsruhe, Germany; German Centre for Integrative Biodiversity Research (iDiv) Halle - Jena- Leipzig, Germany; Senckenberg Centre for Human Evolution and Palaeoenvironments Tübingen (SHEP), Germany; Department of Biodiversity, Heinz Sielmann Foundation, Wustermark, Germany; Institute of Landscape Ecology, University of Münster, Münster, Germany; Institute of Agricultural Sciences, ETH Zürich, Zürich, Switzerland; Forage Production and Grassland Systems, Agroscope, Zürich, Switzerland; Institute for Landscape Ecology and Resources Management (ILR), Research Centre for BioSystems, Land Use and Nutrition (iFZ), Justus Liebig University Giessen, Giessen, Germany; Centre for International Development and Environmental Research (ZEU), Justus Liebig University Giessen, Giessen, Germany; Department of Nature Conservation, Heinz Sielmann Foundation, Wustermark, Germany; Ornithology, Natural History Museum Vienna, Vienna, Autria; Freie Universität Berlin, Institute of Biology, Berlin, Germany; Technical University of Munich, TUM School of Life Sciences, Freising, Germany; Berchtesgaden National Park, Berchtesgaden, Germany; Paris Lodron University Salzburg, Department Environment and Biodiversity, Salzburg, Austria; Group for Sustainable Agroecosystems, Department of Environmental Systems Science, ETH Zürich, Zürich, Switzerland; University of Göttingen, Centre of Biodiversity and Sustainable Land Use, Göttingen, Germany; Zhejiang Provincial Key Laboratory of Plant Evolutionary Ecology and Conservation, Taizhou University, Taizhou, China; Ecology, Department of Biology, University of Konstanz, Konstanz, Germany; Helmholtz Centre for Environmental Research (UFZ), Community Ecology Department, Leipzig, Germany; Department of Biological Sciences, University of Bergen, Bergen, Norway

## Abstract

Across the tree of life, organismal functional strategies form a continuum from slow-to fast-growing organisms, in response to common drivers such as resource availability and disturbance. However, the synchronization of these strategies at the entire community level is untested. We combine trait data for >2800 above-and belowground taxa from 14 trophic guilds spanning a disturbance and resource availability gradient in German grasslands. Most guilds consistently respond to these drivers through both direct and trophically-mediated effects, resulting in a ‘slow-fast’ axis at the level of the entire community. Fast trait communities were also associated with faster rates of whole ecosystem functioning. These findings demonstrate that ‘slow’ and ‘fast’ strategies can be manifested at the level of whole ecosystems, opening new avenues of ecosystem-level functional classification.

## Introduction

Understanding how functional strategies respond to environmental drivers is one of the longest standing and most fundamental questions in ecology (e.g. r/K strategist theory (MacArthur and Wilson, 1967), CSR (competitive/stress-resistant/ruderals) strategies (Grime, 1979)). Due to evolutionary trade-offs, species allocate resources differently to their capacity to grow, reproduce and survive, and for several taxa, it is well established that this leads to sets of co-varying traits that represent ecological strategies (Adler et al., 2014; Bakewell et al., 2020; Stearns, 1976). At the community level, a range of positive and negative biotic interactions (Mayfield and Levine, 2010) and abiotic factors constrain which individuals, bearing specific traits, persist in a community (Diaz et al., 1998). In any given environment this is likely to lead to the dominance of the trait set best adapted to local conditions, leading to trait similarity between co-occurring species and resulting in community-level trait co-variation.

Functional strategies, and their trait proxies, have been particularly well studied and characterised in vascular plants, both at the species (Bergmann et al., 2020; Díaz et al., 2016; Grime, 1979; Weigelt et al., 2021) and community levels (Bruelheide et al., 2018), but have also been described for groups as diverse as fishes, arthropods (Grime and Pierce, 2012), and more recently, microorganisms (Westoby et al., 2021a). While the hypotheses underlying such responses are often underdeveloped compared to those for plants, similar drivers of strategy orientation have been identified, namely resource availability and physical disturbance; and these are consistently seen to act concurrently to shape both individual species and community-level strategies traits in terms of growth, reproduction, and survival across the tree of life (Pedley and Dolman, 2014; Simons et al., 2016). Body size in particular is a fundamental trait of functional strategies that shows consistent responses to these drivers, with undisturbed environments filtering for larger (‘slow’) organisms and disturbed environments for smaller organisms (‘fast’) across groups (Pedley and Dolman, 2014; Suraci et al., 2021), a finding that is consistent with general theory (Lytle, 2001).

At the level of guilds and communities, winning strategies can be seen as manifesting as an emergent property, and represented in community-level trait measures, typically the community abundance weighted trait mean (Bruelheide et al., 2018; Garnier et al., 2004) (CWM), which captures the average functional strategy of the community. As traits are selected upon by the biotic and abiotic environment and suboptimal trait values are selected against, CWM trait values can be considered to reflect the locally ‘optimal’ trait value given the regional species pool and environmental conditions of a site (Muscarella and Uriarte, 2016). This is because while there can be significant variation within a community, species with traits closer to the CWM value at any given site should be those with highest fitness and thus highest abundances (Muscarella and Uriarte, 2016). Accordingly, changes in CWM across space and time reflect both turnover in species with different trait values, and variation in their relative abundance, in response to changes to the species pool and environmental conditions. The combined response of multiple trait CWMs thus represents a change in the overall functional strategy at a community level, and this joint response is usually stronger than that of univariate traits (Muscarella and Uriarte, 2016). This means that slow-fast strategy responses – encompassing a range of traits – may emerge at the community level from a concurrent change in individual CWM traits related to ‘slow’ and ‘fast’ strategies. In particular, communities of resource-acquisitive, fast-growing organisms with numerous offspring, a fast pace of life and good dispersal abilities tend to be found in resource-rich and disturbed habitats, while resource-conservative, slow-growing organisms with longer life span and fewer offspring tend to be favoured in undisturbed or resource-poor habitats (Börschig et al., 2013; Daou et al., 2021; de Vries et al., 2006; Pedley and Dolman, 2014; Simons et al., 2016).

Consistent with these common responses at both species and community levels, associations between traits have been reported between interacting guilds, including plants and soil microorganisms (Boeddinghaus et al., 2019; Buzzard et al., 2019; de Vries et al., 2012), plants and arthropods (Le Provost et al., 2017), and plants and frugivores (McFadden et al., 2022). These shared responses between guilds to similar environmental drivers, along with the existence of widespread strategies within guilds, suggest the possibility of synchronised responses across trophic groups and therefore the potential existence of trait syndromes at the level of entire communities. The possibility of such community-level coupling (Ochoa-Hueso et al., 2021) has been discussed previously, in particular in terms of linkages between plants, herbivores and the soil food web (Wardle et al., 2004). However, while there is theoretical and empirical support for a community-level slow-fast trait response, conflicting observations challenge this hypothesis. First, while the slow-fast spectrum is well defined in plants, additional axes of variation in functional and life-history strategies (e.g. reproductive strategies, and their defining traits such as the timing of reproductive onset and reproductive allocation) have been identified in both plants and other organisms (Rodríguez-Caro et al., 2023; Salguero-Gómez et al., 2016), and these may respond to different drivers and dominate the distribution of certain organismal groups, decoupling them from the response of others (Bakewell et al., 2020; Capdevila et al., 2020). Furthermore, guilds might vary in their strength of response to the drivers of the slow-fast functional axis (Allan et al., 2014) and could respond over different spatio-temporal scales (Le Provost et al., 2021) leading to weak overall coupling.

If community-level slow-fast responses to disturbance and resources are present, then they may be driven by shared direct responses to the environment such as rapid reproduction and effective dispersal of both plants and their consumers in highly disturbed environments. In contrast, synchronous responses could also be mediated by bottom-up (Hairston et al., 1960) trophic interactions: at higher resource availability, we expect higher productivity and resource concentrations, and faster rates of resource transfer between trophic levels, encouraging organisms with faster growth and reproduction both above and below ground (Audusseau et al., 2015; Wardle et al., 2004). Thus, the slow-fast response of the entire community to resource and disturbance drivers might emerge from both direct effects (mainly through shared responses to disturbance) and indirect effects (i.e. a trophic cascade driven by the resource availability component).

Finally, many studies have shown that community-level trait measures of individual guilds explain variation in individual ecosystem functions (Buzzard et al., 2019; Cadotte, 2017; Evans et al., 2017). Meanwhile, several recent studies have demonstrated that overall ecosystem functioning can be described in terms of just a few fundamental functional axes (Eskelinen et al., 2020; Manning et al., 2018; Migliavacca et al., 2021). Given this, we predict that the entire community ‘slow-fast’ trait axis corresponds to an ecosystem functioning ‘slow-fast’ axis, with ‘fast’ functioning defined as fast process rates (e.g. high productivity, rapid nutrient turnover).

While theory often considers the effects of resource availability and disturbance as orthogonal (e.g. CSR theory, (Grime, 1979)), these drivers are often confounded in real ecosystems. Agricultural systems in particular, tend to be found on a continuum between low intensity (low disturbance, no nutrient inputs) and high intensity (high disturbance, such as mowing, and high nutrient input). We can thus expect to find a continuum between ‘slow and steady’ to ‘grow fast, die young’ strategies along land-use intensity gradients – and indeed land-use intensification tends to select for faster strategies of plants (Allan et al., 2015), arthropods (Kőrösi et al., 2022; Le Provost et al., 2017), and microorganisms (Leff et al., 2015).

Despite the evidence base presented above, until now, a lack of coordinated multitrophic abundance, functional trait, and ecosystem function data has hindered the investigation of synchronised responses at the level of entire communities, and their link to ecosystem functioning. However, a recent explosion in trait data availability (Gallagher et al., 2020; Kattge et al., 2011; Madin et al., 2020; Tobias et al., 2022), combined with large-scale research platforms that survey multiple organismal groups and ecosystem functions simultaneously (Bruelheide et al., 2014; Fischer et al., 2010; Li et al., 2022), now allows this long-standing question to be addressed. Here, we use data from the large-scale and long-term Biodiversity Exploratories (Fischer et al., 2010) to test the hypotheses that there is a common, whole community-level slow-fast response to disturbance and resource availability in the form of land-use intensity, and that this community-level strategy drives a slow-fast ecosystem functioning response. The Biodiversity Exploratories is, to our knowledge, the most comprehensive data source for multiple guilds, with abundance data available from the same sites and at all trophic levels, both above-and belowground, including bacteria, fungi, protists, as well as plants, invertebrates and vertebrates (Le Provost et al., 2021). These are collected in 150 grassland plots in three regions of Germany. We focus on the land-use intensity gradient of the Exploratories, which is a combined gradient of both resource availability (fertilisation, with high fertilisation being usually applied to the most inherently productive sites) and disturbance (mowing, which is strongly correlated with fertilisation, and grazing) (Blüthgen et al., 2012). Because existing theory on slow-fast strategies was lacking for some groups, we first conducted expert workshops to identify traits expected to represent the slow-fast continuum for each group. Based on pre-defined hypotheses from existing theories, observational studies, and expert knowledge, we selected several traits for each guild to represent the slow-fast continuum, and generated expectations of how these respond to resource availability and disturbance (Figure 1, see Table S1 for full detail). Focusing on these pre-selected traits, we then tested three hypotheses: (H1) There is a synchronous functional response across trophic levels, shifting from ‘slow and steady’ to ‘grow fast, die young’ strategies with increasing land-use intensity. (H2) Land-use intensity drives the slow-fast response of the entire community through both direct and indirect (through a trophic cascade) pathways. (H3) The entire community ‘slow-fast’ trait axis corresponds to an ecosystem functioning ‘slow-fast’ axis, with ‘fast’ functioning defined as fast process rates (e.g. high productivity, rapid nutrient turnover). Together, the entire community and ecosystem functioning axes form a whole ecosystem slow-fast axis.

**Figure 1.**
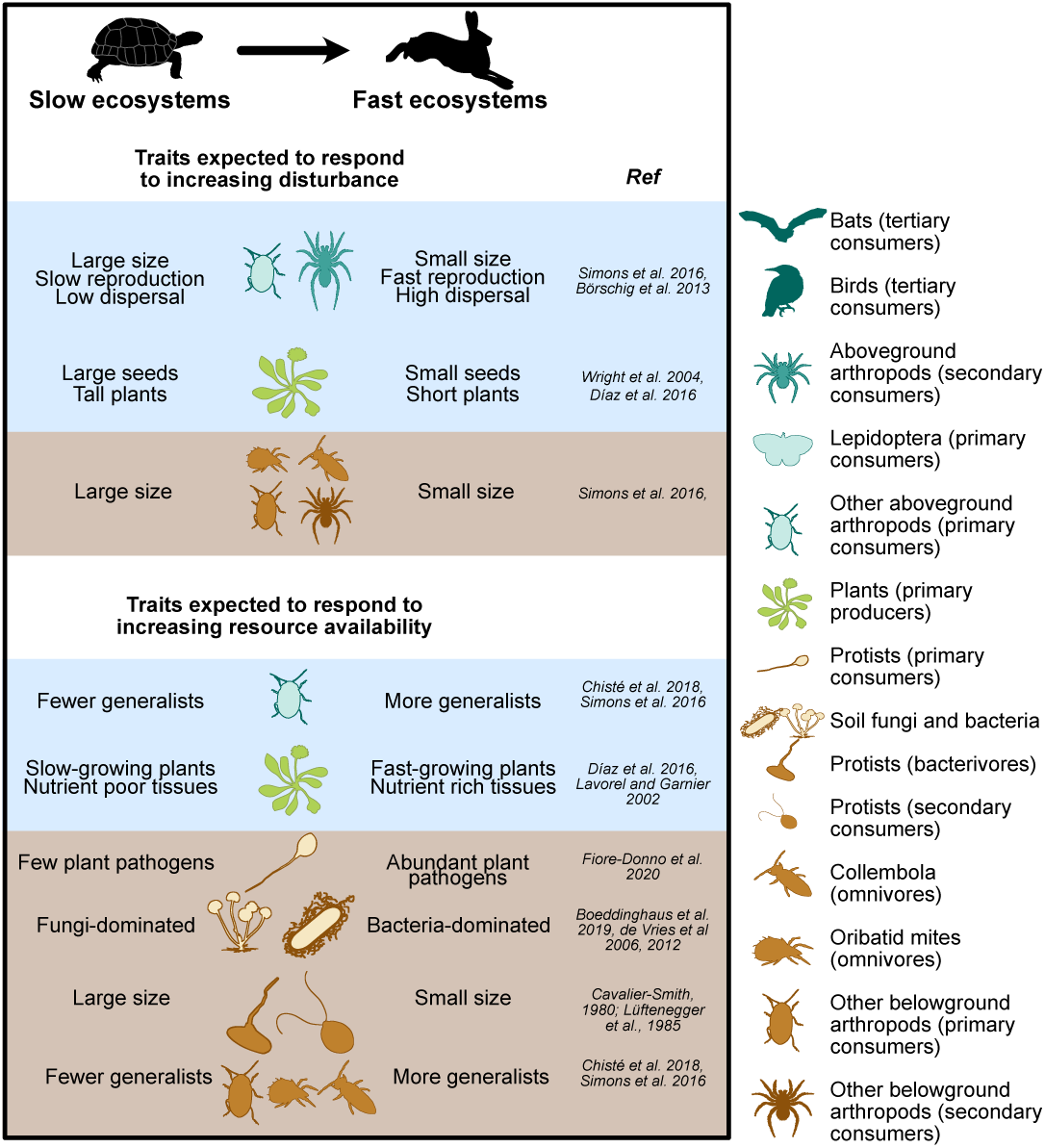
Expected trait variations in ‘slow’ (low resource, low disturbance) and ‘fast’ (high resource, high disturbance) communities. The tortoise and the hare icons represent Aesop’s fable “The Tortoise and the Hare”, which is the origin of the phrase “slow and steady wins the race”, here a potentially winning ecological strategy under conditions of low resource availability and low disturbance. The resource and disturbance gradients are, in this study, represented by a fertilisation and a grazing/mowing gradient. Only a subset of hypotheses is shown, see Table S1 for full details and references. Belowground guilds are shown in brown, aboveground guilds in blue.

## Results

We first identified ‘slow-fast’ traits at the community level within individual trophic guilds. To do this, we classified individual species into trophic guilds of broadly comparable trophic status (e.g. above-ground primary consumers or belowground omnivores). Different taxonomic groups with the same trophic status were aggregated into a single trophic guild if all trait data was consistent, otherwise they were kept separate. For instance, we had comparable trait data for many above-ground arthropods (size, generalism, dispersal ability compiled from Gossner et al., 2015), but data for Lepidoptera was available for different traits (hibernation stage, voltinism, etc) and compiled from a range of other sources (Börshig and Krauss, 2011; Cook et al., 2021; Mangels et al., 2017; Middleton-Welling et al., 2020). Therefore, this group was treated separately from other primary consumer arthropods. This led to the definition of 14 homogeneous trophic guilds (hereafter ‘guilds’), listed in Figure 1 and Methods Table 1. For each guild, abundance-weighted, community-level mean (CWM) trait values were calculated and corrected for environmental covariates (see Methods). The expected trait response of different guilds to resource availability and disturbance is shown in Figure 1 (see Table S1 for full detail).

### Identification of guild-level slow-fast axes

We assumed that for each guild, the main axis of covariation between the selected slow-fast traits (Table S2) should represent the guild’s overall slow-fast response (‘axis’). If the selected traits were not strongly associated (weak covariation) in a principal coordinates analysis (PCA) ordination, we concluded that there was no observable slow-fast axis in the considered guild. Conversely, if most selected traits within a guild covaried, then their main axis of covariation was retained as the guild’s slow-fast axis. This covariation axis was defined through a principal component analysis with all traits within each group, and retained if it explained more than twice of the shared variance as expected if traits were independent; and if the axis was correlated (r > 0.4) with at least 80% of the traits (see specific criteria in Methods).

The existence of the hypothesised guild-level slow-fast axis was supported by the data for most guilds, and it explained on average 52% (+/-sd 13) of the total variation in the 4.2 (+/-1.8) CWM traits per guild that were included in the analyses. For three protist guilds (primary consumers, bacterivores, secondary consumers), only one trait was available, cell size, and we therefore assumed the existence of a ‘slow-fast’ axis defined by large to small cell size in subsequent analyses (Cavalier-Smith, 1980; MacArthur and Wilson, 1967). Of the remaining 11 trophic guilds, we found strong support for a community-level ‘slow-fast’ axis in nine guilds: plants (primary producers), aboveground arthropods (primary consumers), aboveground arthropods (secondary consumers), birds (tertiary consumers), belowground bacteria and fungi (decomposers), belowground arthropods (primary consumers), Collembola (omnivores), Oribatid mites (omnivores), and other belowground arthropods (secondary consumers). For instance, aboveground arthropods (primary consumers) displayed a clear trade-off between ‘slow’ communities dominated by large body size and slow reproduction, and ‘fast’ communities dominated by generalist-feeding species with smaller body size, higher dispersal abilities and multiple generations per year (PC1: 53.8% variance, Table S2). Similarly, Collembola (omnivores) displayed a trade-off between ‘slow’ communities, dominated by soil-surface dwelling species with a large body size and sexual reproduction, and ‘fast’ communities, characterised by more organisms living deeper in the soil and capable of parthenogenesis, which leads to faster generation times (PC1: 65% variance). There was partial support for the existence of a ‘slow-fast’ axis in the remaining two guilds: Lepidoptera (primary consumers) and bats (tertiary consumers) (Table S2).

### Slow-fast trait continuum across trophic levels

Next, we explored the degree of correspondence between the community-level ‘slow-fast’ trait axes of individual guilds. The data supported our overarching hypothesis H1: most of the guild-level ‘slow-fast’ axes were closely correlated with each other and with the land use intensity gradient. A Principal Components Analysis (PCA) conducted on the slow-fast axes of all guilds with sufficient data availability (all except belowground arthropods (primary consumers), see Methods) revealed that 25% of the combined variation in all the guilds’ slow-fast axes was explained by the first PC axis of variation. This axis was strongly correlated with land-use intensity (Pearson correlation r = 0.74, p < 0.001, Figure 2). We also observed a significant and positive correlation between land-use intensity and the slow-fast axis score of nine (out of 14) guilds (p < 0.05 and |r| > 0.2), with this correlation being particularly strong at low trophic levels. Conversely, the highest trophic levels tended to have ‘slower’ traits at high land-use intensity (birds, r = - 0.2). This functional response to land use was also prominent, albeit somewhat weaker, at the level of individual slow-fast related CWM traits, with 53% of these across all guilds (24 out of 45) responding in the hypothesised direction (Figure 3).

**Figure 2.**
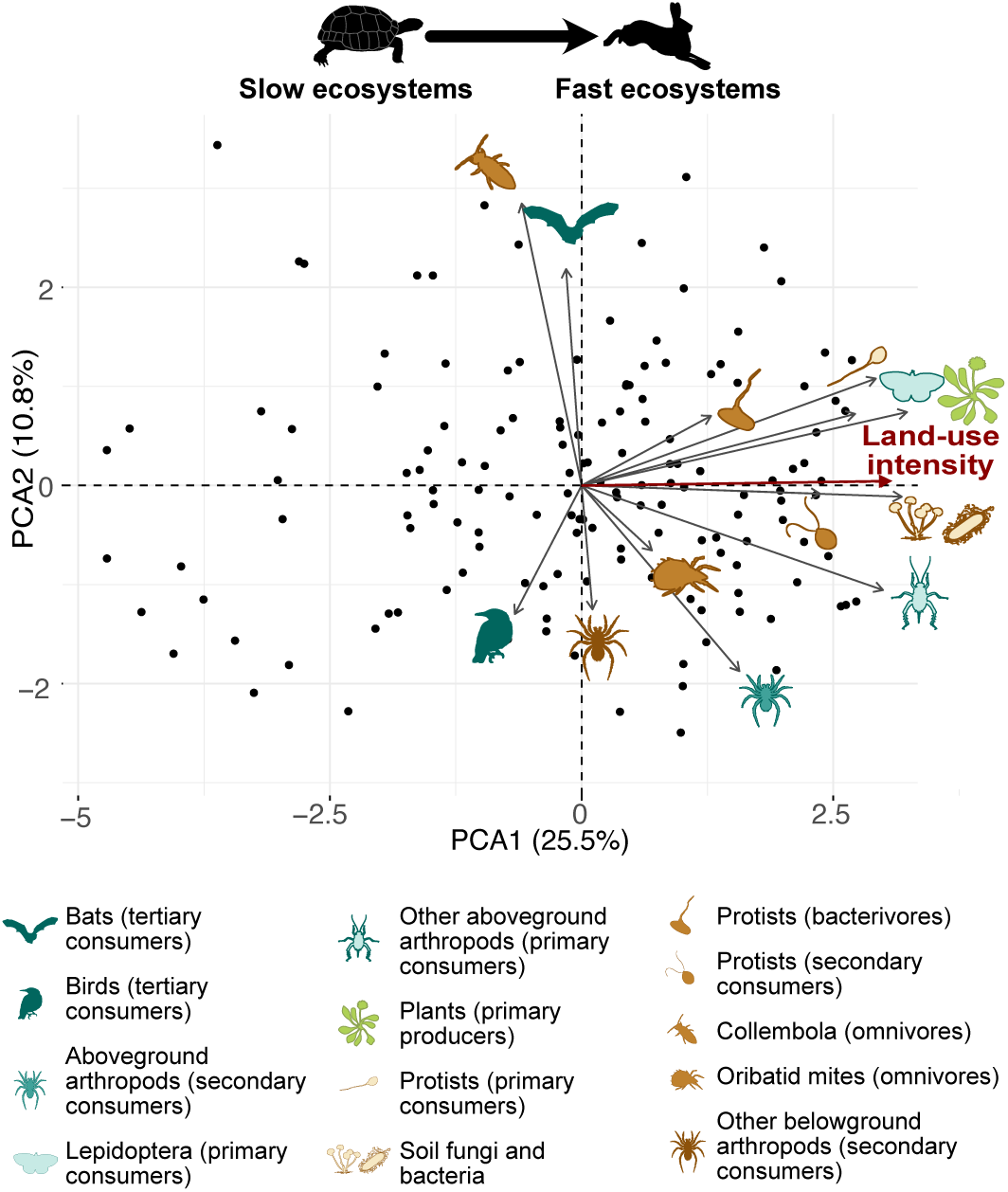
The slow-fast trait axis of individual guilds is strongly related to land-use intensity. The variables included in the PCA are the slow-fast axes of each guild, i.e. the PC axis (PC1 in 90% of cases) of the guild community-level PCA that best represents a slow-fast axis. Land-use intensity (which is projected on the PC axes, shown in red) is strongly associated with Axis 1. Belowground guilds are shown in brown, aboveground guilds in blue, and plants in green. Sample size: 150 (sample sizes for each individual group are shown in Fig. 3, missing values were imputed to run the PCA, see methods).

**Figure 3.**
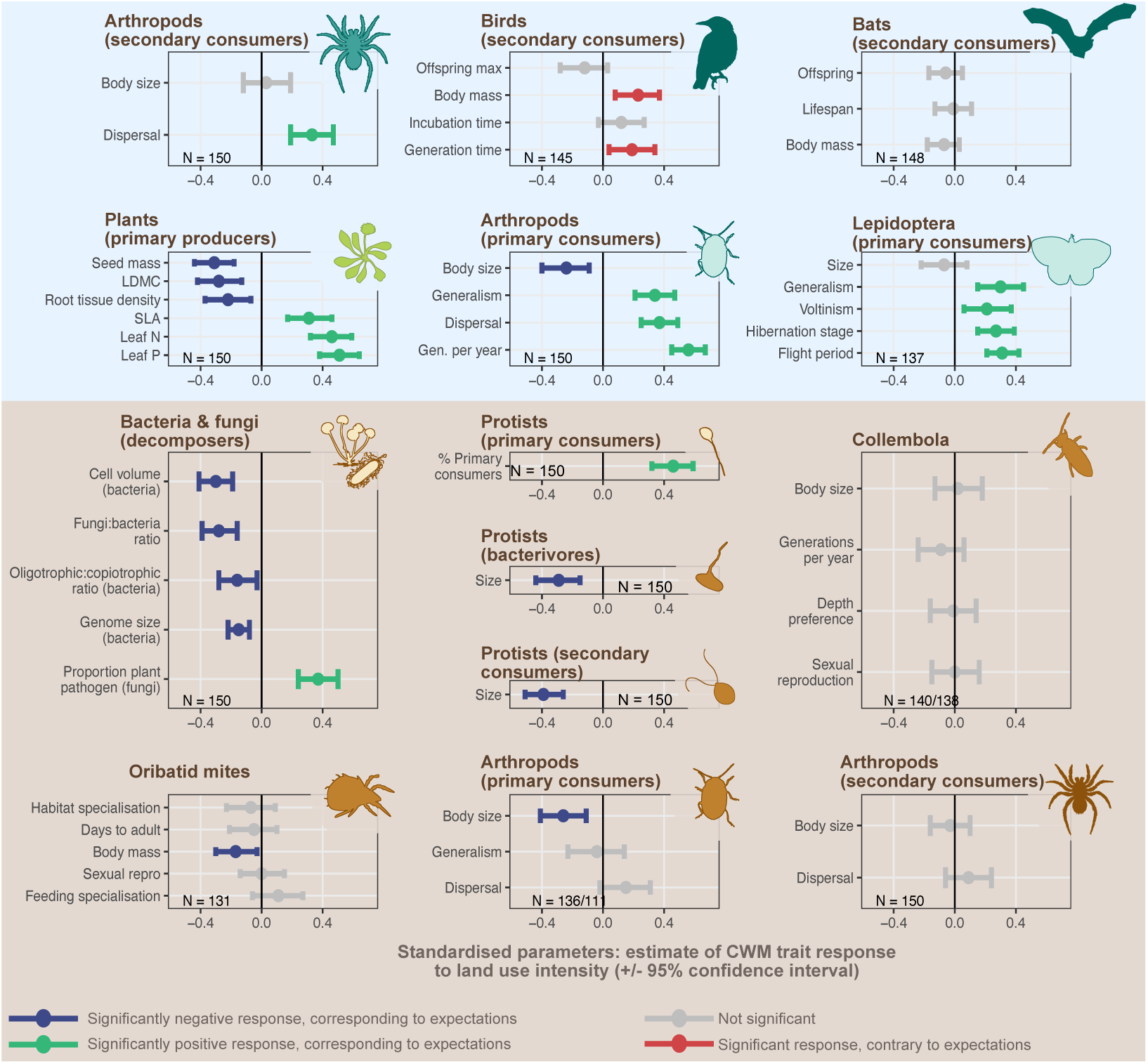
Observed responses of individual traits at the guild level (CWM) to land-use intensity, shown as estimated parameter +/-95% confidence intervals. Belowground guilds are shown above a brown background, aboveground guilds over blue. A standardised response below 0 (blue lines) indicate a negative response to land-use intensity, in accordance with hypotheses (expected higher trait values in ‘slow’ communities). Responses above 0 (green line) indicate a positive response to land-use intensity, in accordance to hypotheses (expected higher trait values in ‘fast’ communities). Red lines indicate traits with opposite responses to that hypothesised. 95% intervals crossing the 0 line indicate no significant response (grey lines). The number of replicates (sites with available data) are shown for each group. Collembola: 138 for the number of generations per year, 140 for the other traits. Arthropods below-ground (primary consumers): 111 for generalism, 136 for other traits.

The observed covariation in slow-fast traits across guilds demonstrates that land-use intensity drives differentiation between entire communities, with ‘slow’ and ‘fast’ communities characterised by distinct trait syndromes (Figure 2, Figure 3, Table S1). ‘Slow’ communities are found at low resource availability and infrequent disturbance, and are characterised by slow-growing resource conservative plants (correlation of leaf dry matter content with land-use intensity: r =-0.28, p <0.001; root tissue density: r = -0.22, p = 0.007). Their soil microbial communities are dominated by fungi rather than bacteria (fungal:bacteria ratio: r = -0.28, p < 0.001), consistent with previous observations that low nutrient availability selects for fungal-dominated communities and slow-growing plants (Boeddinghaus et al., 2019; de Vries et al., 2006). Bacterial communities in ‘slow’ communities also contain a higher proportion of small-celled organisms with large genomes (cell volume: r = -0.30, p < 0.001; genome size: r = -0.15, p < 0.001), an adaptation to growth in nutrient-poor (oligotrophic) conditions (Piton et al., 2020) (oligotrophic:copiotrophic ratio: r = -0.16, p < 0.001). In contrast to the overall trend, birds had slower strategies in high-intensity than low-intensity grasslands (slower generation time, r = 0.19, p = 0.02 and larger body size, r = 0.23, p = 0.005).

At the ‘fast’ end of the spectrum, high resource availability is associated with fast-growing plants with high nutrient content (leaf N content: r = 0.46, p < 0.001, leaf P content: r = 0.51, p < 0.001), which is often related to lower levels of physical and chemical defence (Endara and Coley, 2011). This strategy was associated with a higher dominance of pathogenic fungi (Liu et al., 2021) (proportion of pathogenic soil fungi, r = 0.37, p < 0.001) and plant pathogenic soil protists (Fiore-Donno et al., 2020) (r = 0.46, p < 0.001). In addition, high land-use intensity selected for smaller body size in four guilds (bacterivore protists: r = -0.29, p < 0.001; secondary consumer protists: r = -0.39, p < 0.001, aboveground arthropods (primary consumer) size: r = -0.24, p < 0.001; belowground arthropods (primary consumers): r = -0.26, p = 0.001) and faster reproduction (Börschig et al., 2013) in two guilds (generations per year for Lepidoptera (primary consumers): r = 0.19, p = 0.01; and for other aboveground arthropods (primary consumers): r = 0.56, p < 0.001)). This response is likely driven by disturbance, which causes higher mortality in large bodied arthropods (Birkhofer et al., 2017; Hanson et al., 2016), and favours early reproductive maturity and fast reproduction as organisms can reproduce before disturbance, and population numbers can recover quickly afterwards. In addition, dispersal ability was greater in the fast communities of high land-use intensity (dispersal ability of aboveground arthropod (secondary consumers): r = 0.33, p < 0.001, aboveground arthropods (primary consumers): r = 0.37, p < 0.001). This higher dispersal capacity enables faster recolonization after disturbance (Birkhofer et al., 2017; Simons et al., 2016).

The slow-fast trait responses of some consumer guilds maybe driven not directly by resources and disturbance, but indirectly via losses in plant diversity associated with increased fertilisation (Chisté et al., 2016; Gámez-Virués et al., 2015). This commonly leads to the loss of associated specialist plant species that are typically found in slow ecosystems (Boch et al., 2020; Le Provost et al., 2021; Socher et al., 2012). As such specialist plant species often have higher degrees of physical and chemical defences, they tend to be associated to specialist herbivores (Ali and Agrawal, 2012). In our study, this effect was manifested by an increased dominance of generalist species with land-use intensity, both among Lepidoptera and other aboveground arthropods (primary consumers) (dietary generalism of Lepidoptera: r = 0.30, p = 0.001; and of aboveground arthropods (primary consumers): r = 0.34, p < 0.001).

Changes in community-level traits along environmental gradients can be due to changes in species identity (taxonomic turnover) or variation in their relative abundance. In our case, we found that both factors were responsible for the observed slow-fast responses to land-use intensity within the different guilds (average turnover across guilds: 0.95 +/-0.03; average nestedness: 0.03 +/-0.03; results by using non-abundance-weighted trait metrics are comparable to main results - Table S7 to S12 and Figure S5 to S8).

### Land-use intensity effect on community-level slow-fast response

The observed entire community-level slow-fast continuum could be driven by a common, but independent, response of individual guilds to land-use intensity, or mediated by cascading bottom-up trophic interactions between guilds, and we hypothesised that both pathways were important (Hypothesis 2). To test this, we used Structural Equation Modelling (SEM) to assess the effect of these pathways on the ‘slow-fast’ axes of the 13 guilds. We found that both pathways were important, supporting hypothesis H2; slow-fast strategies axes are shaped both directly via shared responses to land-use intensity and indirectly via trophic interactions. Land-use intensity showed significant direct effects on the ‘slow-fast’ axis of eight guild out of 13 (average estimate: 0.23 +/- 0.06), and indirect, i.e. trophically-mediated, effects on six guilds (0.07 +/- 0.02). The pathways combined to make significant total effects of land-use intensity (0.29 +/- 0.07) on nine out of the 13 groups (Figure 4a).

**Figure 4.**
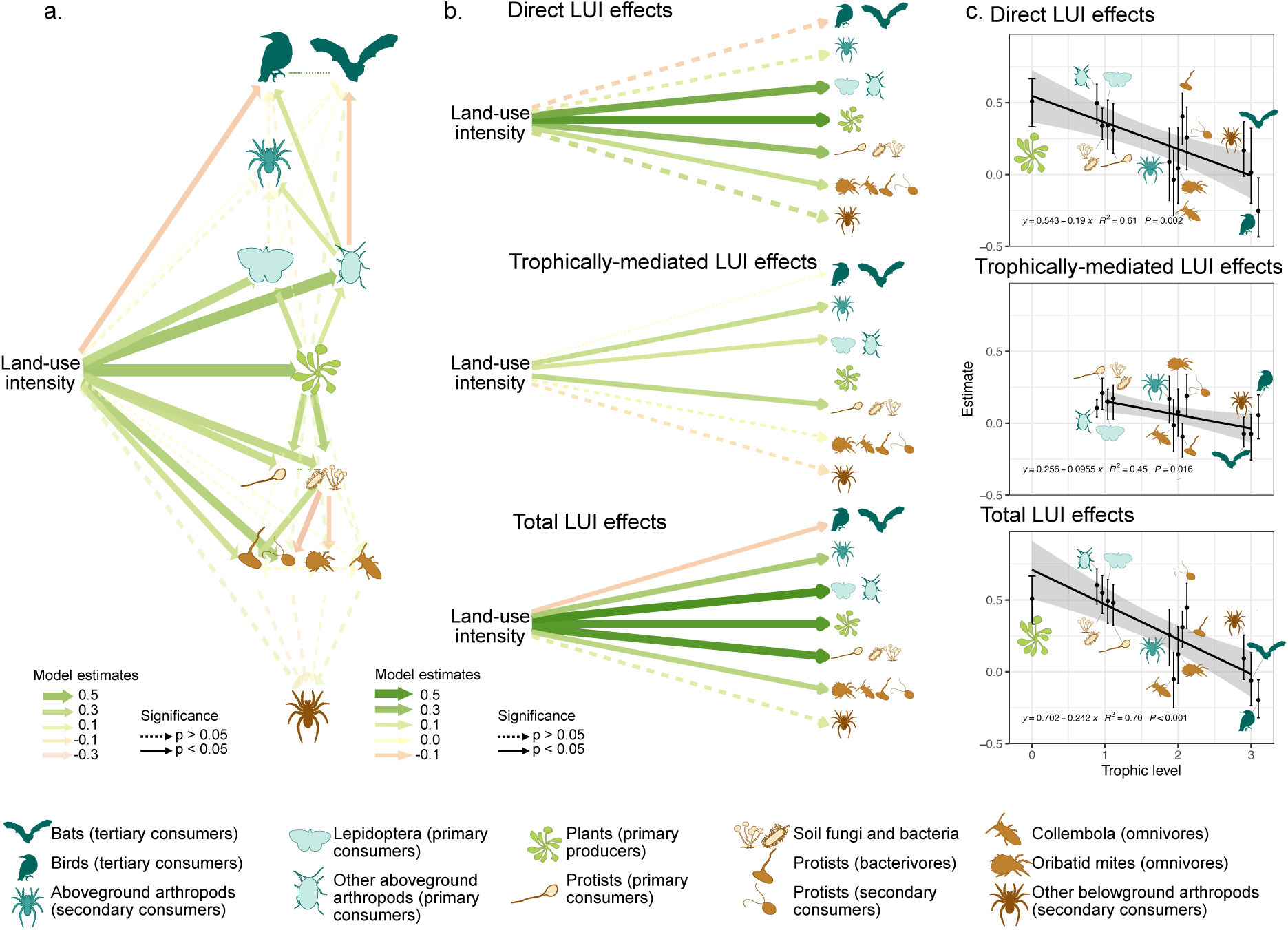
Direct and trophically-mediated effects of land-use intensity on the slow-fast axis of individual trophic guilds. a. Full SEMs including all guilds. Two independent models were fitted for below-and aboveground guilds; plants were included in both. b. Average direct, trophically-mediated (indirect) and total land-use intensity (LUI) effects on each trophic level (averaged across guilds within each trophic level from the SEM estimates shown in a). c. Decreasing direct, indirect and total land-use intensity effects with trophic level. Each dot represents the estimated effect (mean +/- standard error) of land-use intensity on the slow-fast axis of each individual guild in the full SEM. Sampling size per group: ranging from 131 to 150, as shown in Fig. 3. Below-ground, primary consumer arthropods were excluded (see Methods).

Next, we obtained a general assessment of how land-use intensity affected the slow-fast axis of each trophic level, by averaging the direct, indirect and total effects for all guilds at each trophic level (producers vs. primary vs. secondary vs. tertiary consumers). This showed that the ‘slow-fast’ axes of lower trophic levels responded more strongly than higher trophic levels to land-use intensity, via both the direct and indirect paths. While there was a significant total effect of land-use intensity on all trophic levels, except belowground secondary consumers (Figure 4b), the direct (p = 0.002), indirect (p = 0.01) and total (p < 0.001) effects all significantly decreased in intensity with trophic levels, indicating more synchronous trait responses at lower trophic levels.

### Slow-fast ecosystem functioning response

To test our third hypothesis, that the community-level slow-fast continuum was related to whole ecosystem function we first conducted a PCA based on 15 ecosystem functions related to carbon fluxes, nitrogen fluxes, biomass production and decomposition (see Methods Table 5). This showed that ecosystem functions covary, from ‘slow’ sites characterised by slow decomposition, biomass production and low enzyme activities to ‘fast’ sites with faster nutrient cycling (first axis of the PCA: 29% variance, Figure 5a). This ecosystem functioning slow-fast axis was better explained by the entire community ‘slow-fast’ traits axis (r = 0.4, R^2^ = 0.33, p < 10^-6^) than by other hypothesised drivers of ecosystem functioning (single guild community traits measures (plants and microbes), land-use intensity or taxonomic diversity, Table S5). A mediation analysis found that the effect of land-use intensity on the ecosystem functioning ‘slow-fast’ axis was both direct and mediated by the functional traits ‘slow-fast’ axis, as both direct and indirect paths were significant and of similar strength (Figure 5b). Together these results support hypothesis 3 and provide evidence that the whole community fast-slow trait continuum is causally linked to a slow-fast axis of whole ecosystem functioning.

**Figure 5.**
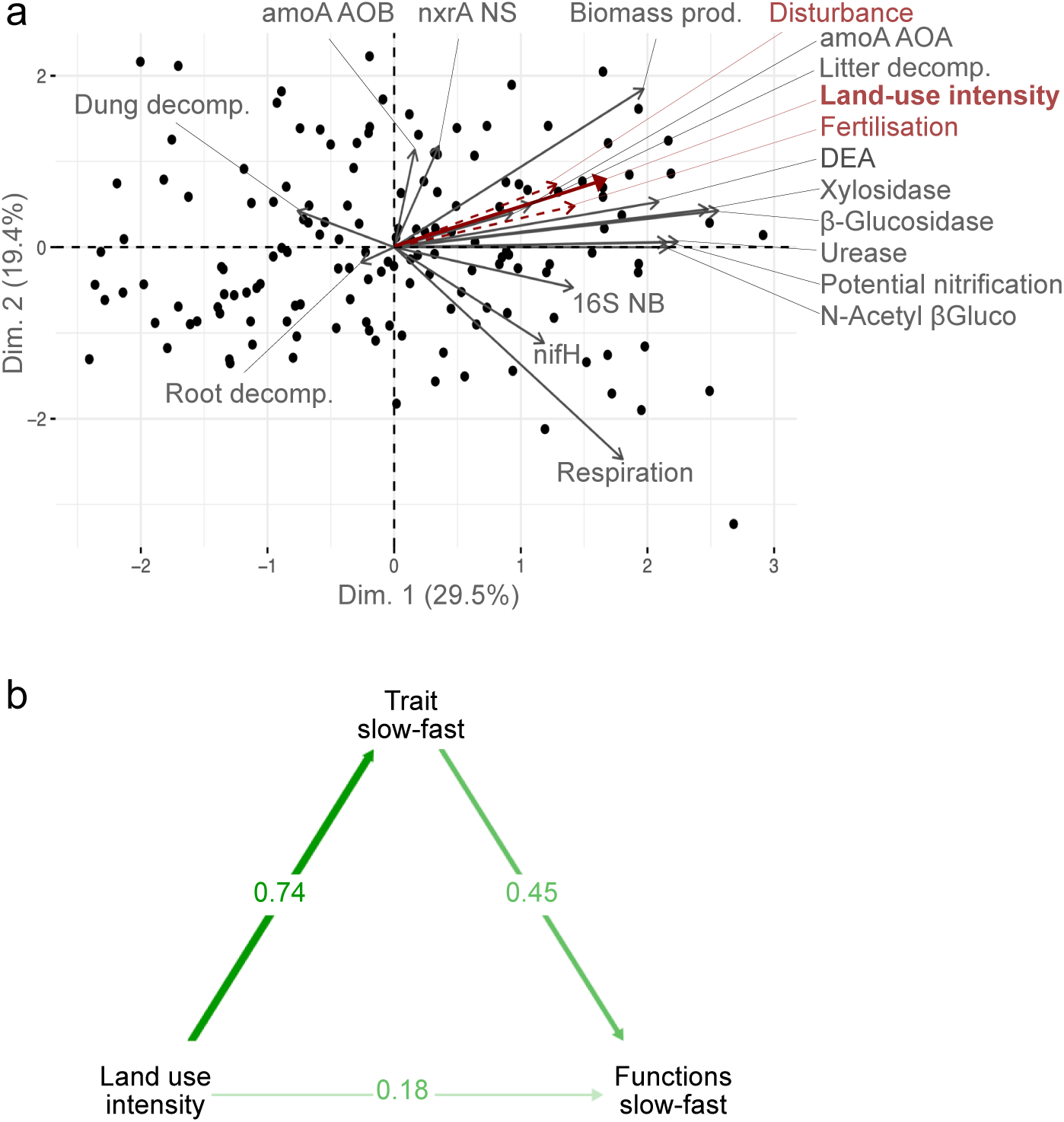
The ecosystem functions slow-fast axis is linked more strongly to the trait slow-fast axis than to land-use intensity. a. PCA of the 15 selected ecosystem functions. Functions within each bundle (carbon fluxes, nitrogen fluxes, biomass production and decomposition) are weighted inversely to the number of functions in the bundle. The first axis shows a slow-fast axis from slow decomposition, biomass production and low enzyme activities to faster nutrient cycling. Land-use intensity and its two components (disturbance and fertilisation) were projected onto the PCA, and are strongly associated with axis 1. b. Direct and indirect links between land-use intensity, the entire community traits slow-fast axis (identified in Figure 1: PC1 of the PCA) and the ecosystem function slow-fast axis. R^2^ lavaan estimate for functions fast-slow axis: 19.6%. R^2^ lavaan estimate for community traits fast-slow axis: 22%. DEA: denitrification enzyme activity; urease, beta-glucosidase, N-Acetyl beta glucominidase, xylosidase: activity of the respective enzyme, amoA_AOA: abundance of ammonia oxidation gene of archaea, amoA_AOB: abundance of ammonia oxidation gene of bacteria, nifH: abundance of nitrogen fixation gene in soil bacteria; nxrA NS: abundance of nitrite oxidation gene of Nitrobacterbacteria; 16S NB: abundance of nitrite oxidizing bacteria; dung decomp: dung decomposition; biomass prod: plant aboveground biomass production; litter decomp: litter decomposition; root decomp: root decomposition. Sample size: 150 sites.

### Sensitivity analyses

In our analyses we were interested in assessing correspondence between guild level strategies at the whole community level. This required the successive aggregation of data into guild-level slow-fast strategies. To assess if this aggregation was affecting our results, we also tested for the existence of the whole-community slow-fast axis by running a PCA on all considered traits from all guilds simultaneously (rather than guild-level slow-fast axes, themselves extracted from PCAs). This analysis also supported the existence of a whole community slow-fast axis, with ‘slow’ and ‘fast’ traits clearly separated across sites (Figure S3). This all-traits slow-fast axis showed even stronger mediation of land-use intensity on the ecosystem slow-fast axis (Fig S4).

While there were strong hypothetical reasons to expect body size to be a key trait driving fast-slow variation at the community level, it can respond to a range of drivers and drive life history variation in the absence of other trait responses; as a result, the effect of body mass is often removed in analyses that seek to identify life history trade-offs (Bielby et al., 2007; Stearns, 1983). To assess the robustness of our results to this trait we conducted additional analyses in which we excluded all body size data (resulting in the exclusion of bacterivores and predator protists as other trait data were not available for these groups). Even in the absence of body size were still able to identify a strong guild-level slow-fast axis for most groups, except collembola. This resulted in somewhat weaker, but still consistent results regarding the synchrony of slow-fast axes across guilds and the effect of the whole community slow-fast axis on ecosystem functioning (Table S18, Figure S14, S15).

Other sensitivity analyses included checking the impact of correcting for environmental covariates and using abundance-weighted trait values. These are discussed in the supplementary analyses (Fig. S4-S13).

## Discussion

Our results provide strong evidence for the existence of synchronous, whole ecosystem-level responses to environmental drivers, and more specifically the existence of a slow-fast axis of variation at the level of entire multitrophic communities. We show that there are similar functional responses to resource availability and disturbance across taxa and trophic levels, from large organisms with slow reproduction and slow dispersal at low land use-intensity to small, fast-paced organisms in more intensively managed sites. The finding of this axis extends earlier studies which investigated the coordinated responses of a few trophic guilds (de Vries et al., 2012; Garibaldi et al., 2015; Le Provost et al., 2017; McFadden et al., 2022) and demonstrates a previously unrecognised emergent property of multitrophic community assembly. We also demonstrate that entire community-level synchrony is driven by both the shared responses of individual guilds to land management and trophically-mediated cascades. Synchrony was stronger at the lower trophic levels which make up the vast majority (>99%) of community biomass (Bar-On et al., 2018), with weaker effects for the less abundant organisms of higher trophic levels (e.g., birds). This is likely due to higher trophic level organisms, e.g. birds and bats, being less dependent on local management conditions and instead responding to larger-scale drivers, such as landscape composition (Le Provost et al., 2021). Finally, we provide evidence that this whole-community strategy variation mediates the effect of land use intensity on overall ecosystem functioning, extending previous findings that demonstrate the linkages between traits of individual guilds and ecosystem functions (Buzzard et al., 2019; Cadotte, 2017; Evans et al., 2017).

The theory describing of slow and fast strategies, and the identity of their defining traits, is very well developed in some organisms such as plants (Díaz et al., 2016; Reich, 2014; Salguero-Gómez et al., 2016). In most other taxa, however, such theories are only emerging (e.g. micro-organisms (Westoby et al., 2021), arthropods (Gibb et al., 2023)). Here, we built upon both in-depth expert discussions and observational studies to define and test hypotheses on the responses of individual taxa to resource availability and disturbance. While this allowed us to generate insights into possible slow-fast strategies for multiple functional groups our need to do this also highlights the need for further theory development on the functional strategies of many taxa. Such theoretical advances would provide key building blocks towards the improved understanding of the response of slow-fast functional strategies to environmental drivers at the level of individual organisms, guilds and entire communities. Such theory could also support the integration of other axes of functional variation, which have been described for individual taxa at species (Bakewell et al., 2020; Capdevila et al., 2020), and community level (DeMalach et al., 2019).

Our results highlight that the average functional strategy of guilds, as represented by the CWM of multiple traits, can be meaningfully related to land use intensity, linkages with other guilds, and measures of ecosystem functioning. By focusing on CWM trait means, we did not consider the role of strategy variation within a community, which can be considerable (Lamanna et al., 2014). This functional diversity can represent either multiple winning strategies within a community and/or niche differentiation from the optimal strategy, and thus the avoidance of competitive exclusion. Functional and taxonomic diversity both within and across taxa has been shown to play an important role in driving multitrophic interactions (Scherber et al., 2010) as well as ecosystem functioning (Walde et al., 2021) and plant diversity is commonly related to higher levels of ecosystem functioning (Allan et al., 2011; Cardinale et al., 2012; Craven et al., 2018). Previous studies have explored the relative response of dominant strategies, versus their variability, in response to both environmental factors (Ricotta and Moretti, 2011) and as drivers of ecosystem functioning (Zhu et al., 2016). Expanding this approach from single groups to whole communities could provide new insights into the behaviour of functional diversity as a multitrophic property that may drive ecosystem structure and functioning at a ‘whole systems’ level.

It is worth noting that the whole ecosystem ‘slow-fast’ gradient observed here was manifested across a relatively short environmental gradient - all sites were temperate agricultural grasslands. It remains to be seen whether the results hold across time (as suggested by dynamic linkages between plant and microbial traits in time (Boeddinghaus et al., 2019)) and for other systems, in particular a diversity of longer, natural and more orthogonal gradients of disturbance and resource availability. We hypothesise that across stronger environmental gradients of climate and soils, e.g. between biomes, and when incorporating a more comprehensive array of slow-fast traits, that ecosystem level trait synchrony will be even more marked. Correspondence between global gradients in climate and soils and plant community-level trait measures (Joswig et al., 2021) support this idea, especially given that the traits of other trophic levels are often strongly associated with plant communities (Boeddinghaus et al., 2019; Buzzard et al., 2019; de Vries et al., 2012). Such synchrony may also correspond to recently described global trends in the covariance of multiple ecosystem functions (Gounand et al., 2020; Migliavacca et al., 2021). When combined with these results, our findings highlight the potential for a new generation of whole ecosystem-level studies that go beyond the description of single trophic guild-level strategies to characterise communities, or single functions to characterise ecosystem responses. Instead, these would work at the level of universal ecosystem-level functional types and axes (Manning et al., 2018; Migliavacca et al., 2021). This may form the basis of a new typology for classifying ecosystems, and provide a generalizable, predictive and simple means of describing their responses to land-use and environmental change.

## Material and methods

### Study area

The study was conducted as part of the long-term Biodiversity Exploratories project (www.biodiversity-exploratories.de). Data was collected in 150 grassland plots in three regions of Germany: the Schwäbische Alb plateau and UNESCO Biosphere Reserve in south-western Germany; the Hainich National Park and surrounding areas in central Germany (both are hilly regions with calcareous bedrock) and the UNESCO Biosphere Reserve Schorfheide-Chorin in the post-glacial lowlands of north-eastern Germany. The three regions differ in climate, geology and topography, but each is characterised by a gradient of grassland land-use intensity that is typical for large parts of temperate Europe (Fischer et al., 2010). In each region, 50 plots (50 m × 50 m) were chosen in secondary wet, mesic and dry grasslands by stratified random sampling from a total of 500 candidate plots on which initial vegetation, soil and land-use surveys were conducted. This ensured that plots covered the whole range of land-use intensities and management types, while minimising confounding factors such as spatial position or soil type.

### Land-use intensity

Land use intensity was assessed annually via questionnaires sent to land managers in which they reported the level of fertilisation (kg N ha^−1^ yr^−1^), the number of mowing events per year (from one to three cuts, starting in May and continuing up to September/October in the most intensive plots), and the number and type of livestock and their duration of grazing (number of livestock units ha^−1^ yr^−1^)(Blüthgen et al., 2012; Fischer et al., 2010). Mowing and grazing intensities determine the frequency, and intensity, at which aboveground biomass is removed; and thus represent the intensity of disturbance in the plot. Fertilisation provides additional nutrients, and is typically mostly applied in naturally productive plots: it thus represents a resource availability gradient. In our study system, mowing and fertilisation intensities are positively correlated (r = 0.70), while grazing and mowing intensities are negatively correlated (r = −0.61). Thus, independent effects of each land-use component, and the respective effect of resource availability and disturbance, cannot be reliably estimated. We therefore used a compound index of land-use intensity, characterising a combined resource availability and disturbance gradient. The land-use intensity index (LUI) was calculated as the square-root-transformed sum of standardised measures of global mowing, fertilisation and grazing intensities across the three regions for each year (Ostrowski et al., 2020). We calculated the mean LUI for each plot over the years 2008–2018 because this reflects the average LUI around the years when most of the data was collected. At low LUI (0.5–0.7), grasslands are typically neither fertilised or mown, but grazed by one cow (>2 years old) per hectare for 30 days (or one sheep per hectare for the whole year). At an intermediate LUI (around 1.5), grasslands are usually fertilised with less than 30 kg N ha−1y−1, and are either mown twice a year or grazed by one cow per hectare for most of the year (300 days). At a high LUI (3), grasslands are typically intensively fertilised (60–120 kg N ha−1y−1), are mown 2–3 times a year or grazed by three cows per hectare for most of the year (300 days), or are managed by a combination of grazing and mowing. The study area did not cover very high intensity grasslands (e.g. cut more than three times per year and ploughed annually).

In some figures, LUI was added as a supplementary variable to e.g. PCAs, along with a ‘fertilisation’ variable and a ‘disturbance’ index. The fertilization variable is the standardized fertilization value. The disturbance index is calculated as the square-root of the sum of mowing and grazing intensities, both standardized by dividing by their overall average.

### Plot species abundance

In each plot, we measured the relative abundance of multiple guilds using standard methodology. We sampled vascular plants in an area of 4 m × 4 m on each plot, and estimated the percentage cover of each occurring species every year from 2008 to 2019. Adult Aranaea, Coleoptera, Hemiptera and Orthoptera were sampled by conducting 60 double sweeps along three 50-m plot-border transects; those spending much of their lifecycle below-ground (e.g. as larvae) were classified as below-ground arthropods. Lepidoptera were recorded along three 300-m transects, each during 30 min. Birds were recorded using audio-visual point counts, at the centre of each respective grassland plot (50 m × 50 m). Bats were sampled along two 200-m plot-border transects: acoustic recordings were taken in real time with a Pettersson-D1000x bat detector (Pettersson Electronic AG, Uppsala, Sweden). To sample belowground bacterial, fungal and protist communities, fourteen soil cores (diameter 4.8 cm) were taken from a 20 m × 20 m subarea of each grassland plot, and soil from the upper 10 cm of soil was homogenised after removal of root material. The bulk sample was split into representative subsamples to analyse each group. Oribatid mites and Collembola were sampled using a Kempson extraction from four soil cores of 4.5 cm × 10 cm per plot. Details of each survey can be found in Methods Table 2.

These taxa were compiled into functional guilds in which trophic status was broadly comparable (e.g. above-ground primary consumers or belowground omnivores). Within each functional guild, organisms of different taxa were aggregated if trait data was comparable, and if not, the functional guild was classified at the taxonomic level (Methods Table 1). Aboveground guilds included vascular plants (primary producers); Lepidoptera (primary consumers – incl. butterflies and day-flying moths); other above-ground primary consumers arthropods (incl. primary consumers of Orthoptera, Coleoptera, and Hemiptera); above-ground secondary consumers arthropods (including omnivorous Hemiptera, carnivorous Coleoptera, Araneae, Orthoptera and Hemiptera); bats (tertiary consumers); and birds (tertiary consumers, i.e. excluding herbivorous and granivorous birds). Belowground guilds included: bacterial and fungal communities (symbionts, decomposers and parasites); plant parasite protists (primary consumers); bacterivorous protists; secondary consumers protists (incl. predatory and omnivores); Collembola (omnivores); Oribatid mites (omnivores); other herbivores, detritivores and fungivores belowground arthropods (primary consumers, incl. Coleoptera and Hemiptera); and predatory and omnivorous, soil-dwelling arthropods (incl. Araneae, Coleoptera, and Hemiptera).

### Functional trait data

Although it is general in overall concept, and has been discussed in a wide range of contexts, the spectrum of slow-fast strategies and their associated trade-offs has a well-developed, trait-specific theory for only a few specific taxa, such as plants. Such theories are also emerging in the literature for other taxa (Westoby et al., 2021a, 2021b) but in many cases are not fully developed yet. Selecting appropriate traits thus required us to combine hypotheses from existing group-specific theories (Díaz et al., 2016), more general ecological theories (e.g. r/K, (MacArthur and Wilson, 1967; Stearns, 1976), empirical observations, and expert knowledge. We conducted multiple expert consultation discussions within subteams of the authors to identify traits potentially related to a resource availability and disturbance, and thus potentially falling on a slow-fast axis, i.e. size, dispersal abilities, feeding niche, etc. Detailed hypotheses guiding the choice of traits for each guild can be found in Table S1.

Trait data was obtained from multiple sources. Abundance data and trait databases were matched using the GBIF taxonomy (package traitdataform (Schneider, 2020)) when necessary. Guild-specific details on trait data acquisition and treatment can be found in Methods Table 3. Briefly, aboveground plant functional traits were extracted from the TRY database (Kattge et al., 2011), aggregated from Central European measurements. Belowground plant trait data was determined in pot experiments for most plant grassland species, and this represented at least 90% cover of all plots (Lachaise et al., 2021). Bird trait data was assembled from the literature (Bird et al., 2020; Pigot et al., 2020; Renner and Hoesel, 2017; Tobias et al., 2022; Wilman et al., 2014). Bat trait data was extracted from Conenna et al., 2021 and Wilkinson and South, 2002. We used data from Gossner et al., 2015, completed using additional sources, for Aranaea, Coleoptera, Hemiptera and Orthoptera; the traits were recoded to allow ranking of some traits (e.g. dispersal ability) across these four orders. Day-flying moths and butterfly traits were combined from previous data syntheses (Cook et al., 2021; Mangels et al., 2017; Middleton-

Welling et al., 2020). Oribatid mites and Collembola traits were collated from the literature and completed with expert assessments for voltinism. Protist traits were extracted from literature reviews (Dumack et al., 2020), completed with expert assessments for size data. Microbial (fungi and bacteria) traits covered both community-level properties that serve as trait proxies (C:N and fungal:bacteria ratio, based on PFLA (phospholipid fatty acids) ratio in a mixed sample from each plot; proportion of parasitic fungi, identified from Illumina MiSeq ASV abundances, functionally assigned through the FUNGuild (version 1.0 (Nguyen et al., 2016)) and individual bacterial traits from Madin et al., 2020, aggregated at the genus or order level when possible.

## Data analysis

All the data analyses were conducted using the R software v. 4.0.3 (R Core Team, 2022).

Initially, in a confirmatory analysis, species-level fast-slow axes were identified from PCA that included all species within most guilds (Table S2).

As we hypothesised changes in community level properties across environmental gradients (rather than changes in individual species), we focused on changes in average trait values. These can represent both changes in species identity (taxonomic turnover) and variation in their relative abundance along the gradient. Thus, all further analyses were conducted at the level of community abundance-weighted trait mean (Garnier et al., 2004) (CWM). This was done using relative abundance data depending on the considered guild (e.g. cover for plants, number of individuals for arthropods, number of ASVs (bacteria, fungi) or OTUs (protists), use of the habitat and activity for birds and bats). As a result of the Biodiversity Exploratories design, data from different guilds was sometimes collected in an uncoordinated way (e.g. every year, every three years, or on fewer consecutive years). To allow comparison across groups, when data was available for more than one year, we first calculated CWM values independently for each year, then aggregated the CWM across years. Data collected in different years, but from the same plot, were considered comparable because both land use intensity and multivariate CWM for all groups that were sampled more than once differed far more across plots than across time. This was shown by conducting variance partitioning analyses (using the varpart function, R package vegan) with either the land use intensity or the trait CWM value for each Plot x Year combination as the response matrix, and the plot and year (as factors) as explanatory matrices. The variance explained by the plot term was on average >15 higher by that explained by the year term (Table S6). Biologically this is because high intensity fields tend to be managed at high intensity year-after-year, and vice-versa.

Each guild was represented by 2 to 6 traits (4.2 on average). Taxa with missing trait data were excluded from the corresponding CWM calculation (except in a few specific cases where data was imputed, see Methods Table 3); the resulting coverage (% of abundance, in terms of cover or individuals with available trait data) was always above 80% except for bacterial and Oribatid mites traits (Table S1). The CWM approach was favoured over joint species distribution modelling of trait-environment relationships (Pollock et al., 2014), as it was the trait values of entire guilds and communities, not species, that was the appropriate unit of replication in our study. It was therefore essential to summarise variables at the whole guild, community, and ecosystem levels prior to analysis.

To gain a more reliable estimate of how CWM trait data was related to the hypothesised drivers, it was necessary to correct for environmental covariates before analysis. As this has been shown to produce biased parameter estimates in case of correlation between the environmental covariates (Freckleton, 2002), we identified highly correlated variables from our list of potential covariates (mean annual temperatures and precipitation, topographic wetness index, soil clay and sand content, pH, depth and region). We excluded soil depth and sand content which were both highly anticorrelated to soil clay content (|r| > 0.72); and precipitations which were highly anticorrelated with temperature (r = -0.77). After that, we fitted linear models with each trait CWM as a response, and the remaining covariates (mean annual temperatures, topographic wetness index, soil clay content and pH as well as the region) as explanatory variables. The residuals or the linear models were used for all further analyses. Details on the measurements of environmental variables and the results of sensitivity analyses with uncorrected values can be found in Methods Table 4, Table S12 to S16 and Figure S9 to S13.

In cases where abundance data was not available for all 150 plots, guild-level analyses were run on all available plots (never < 110 plots). To test for the response of each individual CWM trait to land-use intensity, we calculated correlations (excluding NAs) between each trait and land-use intensity (p-values corrected for false detection rates using the p.adjust function, n = 47). For each guild, we then sought to identify a slow-fast axis based on pre-established hypotheses based on either environmental filtering through resource availability and disturbance, or indirect, trophically-mediated mechanisms (Table S2). Indeed, because traits represent consistent functional strategies, ‘slow’ and ‘fast’ traits are expected to covary. We ran a Principal Component Analysis (PCA) on all CWM traits of each considered guild separately. For these PCA, we used the R package FactoMineR, retained three principal components axes (or two if only two traits were available)) and identified the PCA axis that best represented a slow-fast continuum. This was axis 1 for all guilds except the aboveground predatory arthropods (Table S2). Notably, the fast-slow axis was also always that which most strongly correlated with LUI, except for belowground primary consumer arthropods and birds where axes 1 and 3 (respectively) had a similarly strong correlation (Figure S1). Guild-level PCA axes were considered to represent a slow-fast continuum (and called hereafter ‘guild slow-fast axes’) if they fulfilled all following conditions: i) the considered axis explained at least twice the variance expected at random (i.e. 2*1/(number of traits), with 1/(number of traits) the average variance per axis expected if traits are independent); and ii) the considered axis was significantly correlated with all traits (p-value < 0.05, corrected for false detection rates, n = 34), with at least 60% correlations equal to or above 0.4. The ‘slow-fast’ axis was considered only partially represented if it i) explained at least twice the variance expected at random; ii) was correlated (r >0.25) with at least 60% of the traits, but iii) had correlations in the unexpected direction with one trait.

For analyses across guilds, we excluded one guild (belowground primary consumer arthropods) as more than 20% of plot data were missing due to absence of this functional guild in the samples associated to some plots.

To test whether the slow-fast responses were synchronous across guilds, we ran a PCA on the previously identified slow-fast axes of all guilds, and projected the land-use intensity index (LUI) as a supplementary variable on this PCA. For this PCA only, missing values for the slow-fast axis at the plot level (shown at the individual trait level in Table S1) were given the average of the considered guild across all plots to allow comparison across guilds (imputed values: 1.8% of all values, 0-12% range within groups). This was done to allow comparison across guilds, and because PCA cannot be used with incomplete data. Additionally, we also tested for the existence of the whole-community slow-fast axis by running a PCA on all considered traits (rather than guild-level slow-fast axes, themselves extracted from PCAs, Fig. S3).

In follow-up analyses, we used Structural Equation Models to assess whether the ecosystem level slow-fast continuum was driven by a common, but independent, response of individual guilds to land-use intensity, or mediated by trophic interactions between them leading to a cascade of trait matching. Because plants can affect the traits of other organisms not only through direct consumption but also by e.g. structuring the habitat (Kéfi et al., 2012), we also included paths between all guilds and plants. We also allowed for some correlation paths between guilds when it made biological sense and significantly improved model fit (e.g. between birds and bats, which might be jointly affected by landscape-scale variables). Models were fitted using the lavaan R package (Rosseel, 2012), separately for below and aboveground guilds; plants were included in both. The hypothesised model structures are shown in Figure S2 and model statistics in Table S3 and S4.

The SEM were fitted using the maximum likelihood estimator using all available data (i.e. excluding NAs but using all existing data for each estimate). Estimates were bootstrapped 1000 times. To assess how path strength varied across trophic levels, we extracted the coefficients and corresponding confidence intervals for all direct, indirect and total effects of land-use intensity on each trophic level. We then fitted linear models with estimated direct, indirect and total effects of LUI as the response variables and trophic level as the explanatory variable. Effect estimates were weighted by the inverse of the standard error to account for variable uncertainties across guilds. To evaluate the overall ecosystem response to LUI, we averaged the direct, indirect and total effects at each trophic level, independently above-and belowground. This was done by defining custom parameters within the SEM as the average of all corresponding parameters (direct/indirect/total for all guilds within each trophic level). This allowed the same bootstrap procedure to also estimate average effects and their confidence intervals.

Finally, we investigated the possible linkage between the entire community slow-fast axis and a potential ecosystem functioning slow-fast axis. We selected 15 ecosystem function indicators from 5 bundles of related functions: decomposition (dung, litter and root decomposition), nitrogen cycling in soils (potential nitrification, activity of urease and denitrification enzyme, gene abundances of ammonia oxidising bacteria and archaea (amoA AOA and amoA AOB), abundances of nitrogen fixation (nifH) and nitrite reductase genes (nxrA)), functions related to the carbon cycle in soils (activity of beta-glucosidase, N-acetyl-beta-glucosaminidase, xylosidase), aboveground biomass production and total soil respiration (Methods Table 5). All functions were corrected for environmental variables (as described above for traits) before analysis. All selected functions were hypothesised to represent ‘fast’ ecosystem functioning (e.g. fast nutrient cycling) at high values.

We conducted a PCA on all functions, weighted so that each functional bundle would have the same final weight (e.g. functions related to N fluxes were weighted ⅛, decomposition functions were weighted ⅓, etc.). This was done to prevent bundles with more functions (e.g. N fluxes) from having a disproportionate impact upon the PCA. The first axis (29% of total variance) represented a continuum from slow to fast nutrient cycling with high potential nitrification, and high activities of most enzymes. The plot-level PC scores of this ecosystem functions axis were then regressed against the ‘slow-fast’ axes of individual guilds (microbes and plants, expected to be the main driver of soil functioning), the overall ecosystem functional trait slow-fast axis, land-use intensity and multidiversity, a measure of the overall species diversity of all considered groups (Table S5). This was calculated as the average richness of all taxonomic groups, scaled by the total species richness of each group (Allan et al., 2014), and also corrected for environmental variables, as described above. Finally, to test the direct and indirect effect of land-use intensity on the slow-fast ecosystem functioning axis, we fitted a SEM (using the lavaan package) testing the indirect effect via the entire community slow-fast axis (Figure 5b).

Additionally, we tested for the direct and indirect effects of land-use intensity on the ecosystem slow-fast axis, but on the community slow-fast axis that was derived from the ‘all traits’ PCA (rather than guild-level slow-fast axes extracted from individual PCAs, Fig. S4).

## Acknowledgements

Konstans Wells, Swen Renner, Kirsten Reichel-Jung, Sonja Gockel, Kerstin Wiesner, Katrin Lorenzen, Andreas Hemp, Martin Gorke maintained the plot and project infrastructure; Simone Pfeiffer, Maren Gleisberg, Christiane Fischer, Jule Mangels and Victoria Grießmeier provdided administrative support, Jens Nieschulze, Michael Owonibi and Andreas Ostrowski provided database management. Eduard Linsenmair, Dominik Hessenmöller, Daniel Prati, Ingo Schöning, François Buscot, Ernst-Detlef Schulze, Wolfgang W. Weisser and Elisabeth Kalko helped establish the Biodiversity Exploratories project. The administration of the Hainich National Park, the UNESCO Biosphere Reserves Swabian Alb and Schorfheide-Chorin and all land owners provided logistical support. Tom Lachaise, Jörg Overmann and Johannes Sikorski contributed data.

## Funding

The work was partly funded by the DFG Priority Program 1374 ‘Infrastructure-Biodiversity-Exploratories’, and by Senckenberg biodiversity and Climate Research Center. Field work permits were issued by the state environmental offices of Baden-Württemberg, Thüringen, and Brandenburg.

## Authors contribution

M.N., F.D.S and P.M. designed the study. M.N. and P.M. conducted the analyses with inputs from G.L.P. and A.L.B. M.N., P.M., G.L.P., D.B., F.d.V., J.B., A.M.F.D., S.G., K.G., A.K., R.S., N.S., J.T., A.Z. participated to workshops and expert discussions to define guild-specific trait lists and hypotheses. M.N. and P.M. wrote the manuscript with significant inputs from G.L.P., A.L.B., D.B., J.B., S.G., T.J., M.G., E.K., K.B., K.J., J. K., C.P., M. Schlöter, S. Schulz, M. Staab, M.v.K. and V.W.

M.N., G.L.P., D.B., J.B., A.M.F.D., K.G., R.S., N.S., J.T., A.Z., M.G., K.J., E.K., J.K., C.P., M. Schloter, S. Schulz, M. Staab, V.W., A.A., S.B., R.B., M.B., F.B., K.D., H.Y.G., N.H., J.H., K.J., V.H.K, T.K., S.M., J.M., S.C.R., N.V.S., I.S., M.S., S. Seibold, S. Socher, E.S., M.T., M.v.K., T.W., M.F. and P.M. contributed data. All authors commented on the manuscript.

Author order was determined as follow: main authors, workshop and discussion participants (alphabetical), other authors with significant input (alphabetical), other data contributors (alphabetical), senior author.

## Competing interests

Authors declare that they have no competing interests.

## Data and code availability

This work is based on data elaborated by several projects of the Biodiversity Exploratories program (DFG Priority Program 1374). Most datasets are publicly available in the Biodiversity Exploratories Information System (http://doi.org/10.17616/R32P9Q). However, to give data owners and collectors time to perform their analysis the Biodiversity Exploratories’ data and publication policy includes by default an embargo period of three years from the end of data collection/data assembly which applies to the remaining datasets. These datasets will be made publicly available via the same data repository. All datasets are listed in Tables 1-4 and corresponding references.

Full code to replicate the analyses is stored on GitHub and will be made openly available before publication.

## Notes

### Competing Interest Statement

The authors have declared no competing interest.

