## Supplementary material for "A fast-slow trait continuum at the level of entire communities"

**List of Methods Tables**

**List of supplementary Tables**

**Table S 1.** Trait-specific hypothesis testing: expected response of each trait to resource availability (fertilisation) and disturbance (mowing/grazing); overall coverage (number of individual with available trait data / total number of individual for each plot, median (min-max)); number of taxa with data available / total number of taxa; Expected correlation with land-use intensity; test of the hypothesised response. P-values were corrected for multiple testing (false detection rate, n = 47). ........ 15

**Table S 7.** Abundance-based nestedness (balanced variation) and turnover (abundance gradient) components of Bray-Curtis dissimilarity. The components of dissimilarity were calculated either across all plots, or by first aggregating the 10 highest- and lowest-LUI plots and calculating the dissimilarity between the two groups. Dissimilarities were then calculated using the beta.multi.abund function (package betapart). Colours indicate trophic level (pale to dark colours) and position above (blue) or belowground (brown). ........ 44

**List of supplementary Figures**

**Figure S 1.** Correlation between guild-level PCA axes or single traits and land use intensity. Three axes were retained when more than two traits were available; otherwise only two were retained. For protists, only one trait was available, and the correlation between the CWM of this trait and LUI is shown. The fast-slow axis is always the first PC axis, except for arthropod (secondary consumers) above-ground, for which it was PC axis 2. The axes were transformed were needed (inversed sign) so that higher axis values indicate “faster” strategies. P-values were corrected for false detection rates (***: P < 0.001, **: P < 0.01, *: P< 0.05, n.s.: P > 0.05) ........ 39

**Figure S 5.** Correlation between guild-level PCA axes or single traits and land use intensity. Three axes were retained when more than two traits were available; otherwise only two were retained. For protists, only one trait was available, and the correlation between the CWM of this trait and LUI is shown. The fast-slow axis is always the first PC axis, except for arthropod (primary and secondary consumers) above-ground, for which it was PC axis 2. The axes were transformed were needed (inversed sign) so that higher axis values indicate “faster” strategies. P-values were corrected for false detection rates (***: P < 0.001, **: P < 0.01, *: P< 0.05, n.s.: P > 0.05). **Community-level trait data (CWM) was not weighted by taxa abundance.** ........ 53

**Figure S 6.** Synchronised slow-fast trait response of individual guilds is strongly related to land-use intensity. **Community-level trait data (CWM) was not weighted by taxa abundance**. The variables included in the PCA are the slow-fast axes of each guild. Land-use intensity, added as a supplementary variable, was strongly associated with axis 1. Belowground guilds are shown in brown, aboveground guilds in blue. ........ 58

**Figure S 7.** Direct and trophically mediated effects of land-use intensity on the slow-fast axis of different trophic levels. **Community-level trait data (CWM) was not weighted by taxa abundance**. a. Full SEMs including all guilds. Two independent models were fitted for below- and aboveground guilds; plants being included in both. b. Average direct, indirect and total LUI effects on each trophic level (averaged from the full SEM). c. Decreasing direct, indirect and total LUI effects with trophic level. Each dot represents the estimated effect (+/- standard error) of an individual guild in the full SEM. ........ 59

**Figure S 8.** Direct and indirect links between land- use intensity, functional traits slow-fast axis and ecosystem function slow-fast axis. **In contrast to Fig. S4, community-level trait data (CWM) was not weighted by taxa abundance** ........ 66

**Figure S 9.** Correlation between guild-level PCA axes or single traits and land use intensity. Three axes were retained when more than two traits were available; otherwise only two were retained. For protists, only one trait was available, and the correlation between the CWM of this trait and LUI is shown. The fast-slow axis is always the first PC axis, except for arthropod (secondary consumers) above-ground, for which it was PC axis 2. The axes were transformed were needed (inversed sign) so that higher axis values indicate “faster” strategies. P-values were corrected for false detection rates (***: P < 0.001, **: P < 0.01, *: P< 0.05, n.s.: P > 0.05). **Community-level traits (CWM) were not corrected for environmental covariates**. ........ 78

**Figure S 10.** Synchronised slow-fast trait response of individual guilds is strongly related to land-use intensity **Community-level traits (CWM) were not corrected for environmental covariates**. The variables included in the PCA are the slow-fast axes of each guild. Land-use intensity, added as a supplementary variable, was strongly associated with axis 1. Belowground guilds are shown in brown, aboveground guilds in blue. ........ 79

**Figure S 11.** Direct and trophically mediated effects of land-use intensity on the slow-fast axis of different trophic levels**. Community-level traits (CWM) were not corrected for environmental covariates.** a. Full SEMs including all guilds. Two independent models were fitted for below- and aboveground guilds; plants being included in both. b. Average direct, indirect and total LUI effects on each trophic level (averaged from the full SEM). c. Decreasing direct, indirect and total LUI effects with trophic level. Each dot represents the estimated effect (+/- standard error) of an individual guild in the full SEM. ........ 80

**Figure S 15.** Direct and trophically mediated effects of land-use intensity on the slow-fast axis of different trophic levels**. Analyses excludes size- and body mass- related traits.** a. Full SEMs including all guilds. Two independent models were fitted for below- and aboveground guilds; plants being included in both. b. Average direct, indirect and total LUI effects on each trophic level (averaged from the full SEM). c. Decreasing direct, indirect and total LUI effects with trophic level. Each dot represents the estimated effect (+/- standard error) of an individual guild in the full SEM. ........ 94

**Methods Table 1.** Trophic guilds taxonomic composition and assigned trophic position. Colours indicate trophic level (pale to dark colours) and position above (blue) or belowground (brown)

| Guild | Taxa included | Trophic position |
|---|---|---|
| Birds (tertiary consumers) | Insectivorous and predatory birds | 3 |
| Bats (tertiary consumers) | Bats | 3 |
| Herb- and litter-dwelling arthropods (secondary consumers) | Omnivores Hemiptera, carnivorous Coleoptera, Araneae, Orthoptera and Hemiptera | 2 |
| Lepidoptera (primary consumers) | Butterflies and day-flying moths | 1 |
| Other herb- and litter-dwelling arthropods (primary consumers) | Herbivorous Orthoptera, pollinator, herbivorous and decomposer Coleoptera, and herbivorous Hemiptera | 1 |
| Vascular plants (primary producers) | All vascular plants | 0 |
| Microbial communities (decomposers) | Bacteria and fungi, including saprotrophs and parasites | 1 |
| Protists (primary consumers) | Plant parasites | 1 |
| Protists (bacterivores) | Bacterivorous protists | 2 |
| Protists (secondary consumers) | Predatory and omnivore protists | 2 |

| | | |
|---|---|---|
| Collembola (omnivores) | Collembola | 2 |
| Oribatid mites (omnivores) | Oribatid mites | 2 |
| Other belowground arthropods (primary consumers) | Herbivores, detritivores and fungivores coleoptera and Hemiptera | 2 |
| Predatory and omnivorous, soil-dwelling arthropods (secondary consumers) | Predatory or omnivorous Aranea, Coleoptera, and Hemiptera | 3 |

**Methods Table 2.** Sampling protocols for all trophic guilds. Most data was extracted from an aggregated diversity datasets from the Biodiversity Exploratories(Penone et al., 2021a, 2021b); original datasets and their current status (publicly available or not yet) are listed below.

| Taxonomic group | Methods | Data owners and references |
|---|---|---|
| Vascular plants | Visual estimates of % cover of each vascular plant species growing in a 4 m × 4 m subplot, done in 2008-2018 | Boch, Bolliger, Fischer, Heinze, Hölzel, Klaus, Kleinebecker, Müller, Prati, Socher<br><br>Open datasets from Biodiversity Exploratories: Bolliger et al., 2021 |
| Arthropods: Lepidoptera | Sweep netting along transects of 300 m during 30 min, done three times per plot in 2008 | Börschig, Krauss<br><br>Open datasets from Biodiversity Exploratories: Börschig et al., 2013 |
| Arthropods: Araneae, Coleoptera, Hemiptera, Orthoptera | All arthropods of the herb layer were sampled twice per year between 2009 and 2017 in June and August to represent different phenological windows within the peak season of adult arthropod activity. Based on monthly samplings at the beginning of the study, we identified these two months to represent the best trade-off between reducing sampling effort and covering most species. Arthropods were sampled by sweep netting along a 150-m long transect comprising three of the virtual borders of a site by conducting 60 double sweeps per site. Sweep netting was only conducted on days without rain, low wind speed and after morning dew had dried. All samples were sorted to order level in the laboratory. For taxonomic groups that occurred in larger numbers and for which expert taxonomists were available, adult specimens were identified at species level: Araneae, Coleoptera, Hemiptera (Heteroptera and Auchenorrhyncha), Orthoptera. | Gallenberger, Gossner, Lange, Pasalic, Seibold, Simons, Staab, Türke, Weisser,<br><br>Open datasets from Biodiversity Exploratories: Weisser et al., 2019 |
| Birds | Bird surveying data during breeding times (March-June), estimated by audio-visual point-counts at the centre of each plot, done in 2008-2012 and 2018. | Jung, Renner, Teuscher, Tschapka<br><br>Open datasets from Biodiversity Exploratories: Goldmann et al., 2018; Tschapka et al., 2017a, 2017b, 2017c, 2017d |

| | | |
|---|---|---|
| | | Other dataset from the Biodiversity Exploratories: Teuscher and Fischer, 2021 |
| Bats | Acoustic recordings were conducted along the edges of each Experimental Plot, using a combination of point-stop and transect monitoring in 2009 and 2010. Point-stops were located at each corner of the EP. Transects were walked slowly and in a direct line between the point-stops. Survey time at point-stops and during transect walks in forests was 6 min each and in grasslands 3 minutes min each. This resulted in a total survey time of 48 min along a 400 m transects (per EP) in the forest and a 24 min survey time along a 200 m transect (per EP) in the grassland. Recordings were made using a Pettersson D 1000x ultrasound detector (Pettersson Electronic AG, Uppsala, Sweden). Echolocation calls were identified to species level or to Sonotype using the software Avisoft SAS Lab Pro, Version 5.0.24 and onward (Raimund Specht, Avisoft Bioacoustics, Berlin Germany) | Jung, Tschapka<br><br>Open datasets from Biodiversity Exploratories: Jung and Tschapka, 2016a, 2016b |
| Arthropods: Collembola and Oribatid mites | Kempson extraction from four soil cores of 4.5 cm × 10 cm per plot, done in 2019 | Baulechner, John, Wolters, Zaytsev<br><br>Dataset from the Biodiversity Exploratories: Baulechner et al., 2020 |
| Protists | Soil DNA was extracted in 2011 and in 2017, from 400 mg of soil (not sieved), 3- to 6-times, using the DNeasy PowerSoil Kit (Qiagen GmbH, Hilden, Germany) following the manufacturer's protocol. Specific primers for Cercozoa and Endomyxa were used to amplify, by two semi-nested PCRs, the V4 region of the 18S rRNA gene. Libraries were prepared using TruSeqDNA PCR-Free (Illumina, San Diego, CA, US). Sequencing was performed with a MiSeq v3 Reagent kit of 300 cycles (on a MiSeq Desktop Sequencer, Illumina). The bioinformatics pipeline was conducted with Mothur v. 39.5. After assembling and quality filtering, sequences were clustered at 97% similarity, and rare clusters (<0.01% of the reads) were removed. OTUs were identified in the PR2 database using BLASTn with an e-value of $1^{e50}$ and keeping only the best hit. Chimeric OTUs were identified using UCHIME and removed (Fiore-Donno et al., 2020a). | Bonkowski, Fiore-Donno<br><br>Open datasets from Biodiversity Exploratories: Fiore-Donno and Bonkowski, 2019a, p., 2019b |
| Fungi and bacteria | Composite samples were taken in 2011, 2014 and 2017 in all plots, by mixing 14 mineral topsoil samples (0–10 cm, using a split tube manual soil corer with 5 cm diameter).<br>For bacteria the analysis only included data from 2011 and 2014. 10 g of the homogenized soil was put immediately on liquid nitrogen and stored until RNA extraction. RNA was extracted using a custom protocol (Lueders protocol). Total RNA was isolated from soils and reverse transcribed into cDNA. Amplicons of the V3 region of the 16S rRNA gene were sequenced on an Illumina Hiseq platform using universal bacterial primers.<br><br>For fungi, total microbial DNA was isolated from the bulk soil sample using a MoBioPowerSoil DNA Isolation Kit. A PCR approach was used to amplify fungal ITS-rDNA by using the primer pair ITS4/fITS7, containing the Illumina adapter sequences. PCR products were then purified, cleaned and sequenced using Illumina MiSeq | Buscot,<br>Goldmann,<br>Klemmer, Marzini, Overmann, Sikorski,<br>Wubet<br><br>Open datasets from Biodiversity Exploratories: Buscot et al., 2021, 2020; Obermann and Sikorski, 2019<br><br>Other datasets from the Exploratories: Kandeler et al., 2017b, 2017c; Sikorski et al., 2019 |

**Methods Table 3.** Trait data acquisition and treatment. Colours indicate trophic level (pale to dark colours) and position above (blue) or belowground (brown).

| Taxonomic group | Methods | Data owners and references |
| --- | --- | --- |
| Plants | Aboveground plant traits were compiled from the TRY database. TRY data was first cleaned to remove duplicates, non-mature, non-healthy plants and plants which were grown in non-standard exposition as well as non-standard trait measurements. The trait data was then subsetted by data from central Europe to avoid geographic bias. Finally, resulting trait values were averaged by contributing sources, with outlier sources being excluded; and were then averaged for each species.<br>Belowground trait data was obtained from pot experiments(Lachaise et al., 2021).<br><br>Where appropriate, data from synonyms was used. When individuals were identified at the genus level, average trait values for all other species from the considered genus found during the survey was used.<br><br>Community-weighted means were calculated by excluding tree saplings which are not part of the stable grassland communities and do not have the (adult) traits usually reported in databases. | Ref from TRY: (Adler et al., 2014; Adriaenssens, 2012; Akhmetzhanova et al., 2012; Atkin et al., 2015; Baruch and Goldstein, 1999; Blonder et al., 2015; Bond-Lamberty et al., 2002, p.; Boucher et al., 2013; Bragazza, 2009; Burrascano et al., 2015; Buzzard et al., 2019; Cadotte, 2017; Campetella et al., 2011; Cavender-Bares et al., 2006; Chave et al., 2009; Chen et al., 2013; Choat et al., 2012; Ciccarelli, 2015; Cornelissen, 1996; Cornelissen et al., 2001, 2003, 2004, 1996; Craine et al., 2012, 2009; Dahlin et al., 2013; Dalke et al., 2018; Diaz et al., 2004; Dwyer et al., 2014; Fagúndez and Izco, 2008; Falster et al., 2015; Fonseca et al., 2000; Freschet et al., 2010; Giarrizzo et al., 2017; Gonzalez-Akre et al., 2015; Gos et al., 2016; Green, 2002; Guy et al., 2013; Han et al., 2012; Herz et al., 2017a, 2017b; Iversen et al., 2017; Kattge et al., 2011, 2009; Kazakou et al., 2006; Kerkhoff et al., 2006; Kichenin et al., 2013; Kleyer et al., 2008; Koele et al., 2012; Kühn et al., 2004; La Pierre and Smith, 2015; Lachaise et al., 2021; Laughlin et al., 2011, 2010; Lhotsky et al., 2016; Li and Shipley, 2018; Liebergesell et al., 2016; Lin et al., 2015; Louault et al., 2005; Lukeš et al., 2013; Maire et al., 2016, 2015; Martin et al., 2018; Medlyn et al., 1999; Meir et al., 2002; Mencuccini, 2003; Moles et al., 2004; Moretti and Legg, 2009; *NRCS: The PLANTS Database*, 2009; Onoda et al., 2017; Ordoñez et al., 2010; Pahl et al., 2013; Paine et al., 2015; Paula et al., 2009; Peco et al., 2005; Prentice et al., 2011; Preston et al., 2006; Price and Enquist, 2007; Quested et al., 2003; Rolo et al., 2012; Sandel et al., 2011; Scherer-Lorenzen et al., 2007; Schmitt et al., 2017; Schweingruber and Landolt, 2005; Sheremetev, 2005; Siefert et al., 2014; Smith and Dukes, 2017; Smith et al., 2014; Spasojevic et al., 2016; Spasojevic and Suding, 2012; Takkis, 2014; Thuiller, n.d.; Tribouillois et al., 2015; VERGUTZ et al., 2012; Vile, 2005; WALKER, 2014; Wang et al., 2017; Werner et al., 2018; Wilson et al., 2000; Wirth and Lichstein, 2009; Wright et al., 2004; Yguel et al., 2011; Zanne et al., 2009)<br><br>Other dataset from the Exploratories: Neyret et al., 2021 |

| | | |
|---|---|---|
| Birds (tertiary consumers) | Bird trait data for body length and incubation time was obtained from the literature and own datasets. Data on functional groups and trophic levels was extracted from (Pigot et al., 2020) and the Avonet dataset(Tobias et al., 2022). Identification of ground-nesting species followed (McMahon et al., 2020). Maximum longevity and generation length were extracted from (Bird et al., 2020).<br>While trait data was also available for herbivorous species, the abundance data was not sufficient to include this group as 46 plots did not have any primary consumer (primarily herbivorous, frugivorous, granivorous species). Thus, only secondary consumers (including carnivores, invertivores, and omnivores whose diet includes vertebrates or insects) were included in the study. | Penone, Renner, Tobias<br><br>Ref: (Bird et al., 2020; McMahon et al., 2020; Penone et al., 2022; Pigot et al., 2020; Renner and Hoesel, 2017; Tobias et al., 2022) |
| Bats (tertiary consumers) | Bat traits data was obtained from (Conenna et al., 2021) (morphological traits) and (Wilkinson and South, 2002) (life history traits). Both datasets were then restricted to species occurring in Germany based on (Schall and Petermann, 2014).<br>Abundance data was based on acoustic recording, which did not always allow for species-level identification: some individuals were classified as genera (*Myotis sp*., Plecotus sp.) or similarly acoustic groups (Nyctaloids). In that case, traits were attributed based on the average values of species from the same genus present in Germany, or on all species in the Nyctaloid group (*Nyctalus noctula, Vespertilio murinus, Eptesicus serotinus, Nyctalus leisleri, Eptesicus nilssonii*), respectively. The interpolation was usually relatively conservative (trait mean > 2*trait sd). | Conenna et al., 2021; Schall and Petermann, 2014; Wilkinson and South, 2002 |
| Arthropods: Araneae, Coleoptera, Hemiptera and Orthoptera | Body size (in mm), feeding guild, stratum use and dispersal ability of Araneae, Coleoptera, Hemiptera and Orthoptera were assembled from various literature sources and validated in correspondence with taxonomic experts for the respective groups. These were augmented by published trait data on feeding generalism from (Gossner et al., 2015) and voltinism from multiple sources (Bakewell et al., 2020; Neff et al., 2020; Nickel and Remane, 2002; Saulich and Musolin, 2021).<br>Species were classified as secondary consumers if one of their life stages was mostly carnivorous, and as primary consumers otherwise (i.e., all life stages primarily herbivorous, fungivorous or detritivorous).<br>Species were classified as belowground if at least one of their life stages was primarily soil- or ground-dwelling, and aboveground otherwise.<br>Voltinism was coded as a numeric variable (0.5: semivoltine, 1: univoltine; 1.5: uni- or bi-voltine; 2: bivoltine; 3: multivoltine). Dispersal was coded as a numerical variable (in five gradations (from low to high: 0, 0.25, 0.5, 0.75, 1) based on flight ability and wing dimorphism for insects and ballooning for spiders).<br>Body size was log-transformed to avoid non-normal distribution.<br>When sampled individuals were only identified at genus level in the abundance dataset, they were attributed the average traits of the other species in the genus only when the trait data was homogeneous (i.e. dominant category if it is representative of 90% of the species in the genus among the sample species, and if the mean is > 2*standard deviation for quantitative variables); and was considered as a missing value otherwise. | Ref: Bakewell et al., 2020; Neff et al., 2020; Nickel and Remane, 2002; Saulich and Musolin, 2021<br><br>Other datasets from the Exploratories: Staab et al., 2022 |
| Arthropods: Lepidoptera | Traits for Lepidoptera were compiled from multiple databases. We used as primary references (Mangels et al., 2017) for moth traits and (Börschig et al., 2013) for butterfly traits. In addition, we completed this data with traits from (Cook et al., 2021), and the European and Maghreb Butterfly Trait database(Middleton-Welling et al., 2020). When datasets disagreed, we primarily kept the data from (Mangels et al., 2017) which was specifically curated for German moths. | Ref: (Börschig et al., 2013; Cook et al., 2021; “Lepiforum e.V.,” n.d.; Mangels et al., 2017; Middleton-Welling et al., 2020; Wolfgang Wagner, n.d.)<br><br>Open datasets from Biodiversity Exploratories: Blüthgen et al., 2018; Börshig and Krauss, 2011; Mangels and Blüthgen, 2017 |

| | | |
|---|---|---|
| | Voltinism was coded as a numerical variable: 0.5 for semivoltine species, 1 for strictly univoltine species, 1.25 for univoltine species with partial generation, 1.5 for uni- or multivoltine species, and 2 for multivoltine species.<br>Generalism was coded as a numerical variable: 1 for monophagous species, 2 for oligophagous species (within one plant genus), 3 for oligophagous species (within one plant family), and 4 for polyphagous species.<br>Body size was provided as different metrics for the different databases: estimated dry mass, forewing minimum and maximum length in (Cook et al., 2021)), forewing average length in (Middleton-Welling et al., 2020), and body mass and wing length in (Mangels et al., 2017). The most complete was the forewing maximum for (Cook et al., 2021); we used data imputation (function mice, using default parameters) to impute the missing values in this variable, using all other size-related variables as predictors. The distribution of the data was similar before and after imputation, indicating reliable imputation. Body size was then log-transformed.<br>Overwintering stage was coded as a numeric variable, from 1 (overwintering as egg), 2 (larvae), 3 (puppa) and 4 (adult). When datasets disagreed, we retained the latest stage of development as overwintering stage.<br>Flight period was measured as the maximum number of flight months for adults. Again, (Cook et al., 2021) was the most complete database. We used this data when available, and inputed missing values (function mice, default parameters) based on additional data from (Middleton-Welling et al., 2020). Flight period was then log-transformed. We completed the resulting data for three additional species (*Lythria purpuraria*, *Zygaena carniolica, Aphantopus hyperantus*) based on https://lepiforum.org/https://lepiforum.org/ and http://www.pyrgus.de/.http://www.pyrgus.de/. | |
| Arthropods: Collembola | Data on adult body size and depth preference was collected on specimens collected during sampling (see above). Voltinism and reproduction type (sexual or parthogenetic) were provided by expert opinion and completed from the literature. Voltinism was coded as a numeric variable: 1 for multivoltine species and 0 for bivoltine species.<br>One plot had only unidentified species without possibilities to match corresponding trait values: it was removed from the analysis. Two plots had missing values for voltinism community weighted mean value due to missing voltinism values in all the species present in the plot; they were given the average voltinism value of all plots. | Baulechner, John, Wolters, Zaytsev |
| Arthropods: Oribatid mites | Data on adult body mass and habitat specificity was collected on specimens collected during sampling (see above). Habitat specificity was coded numerically from 1 (non-specific habitat, most generalist), 2 (soil-dwelling), 3 (surface-dwelling) and 4 (litter-dwelling, which are the most specific species). Feeding specialisation was coded numerically from 1 (omnivorous, most generalists), 2 (herbifungivorous), 3 (herbivorous) and 4 (fungivorous, most specialist). V. Wolters and A Zaytsev, pers. comm.<br>Time to maturity (in days) and reproduction type (sexual or parthogenetic) were provided by experts and completed from the literature and coded as binary variables.<br>9 plots had trait data coverage under 20% due to the dominance of individuals identified only at the family level (Gamasideae family), making the matching of trait data difficult. These nine plots were given NA values for all CWM values. | Baulechner, John, Wolters, Zaytsev |
| Bacteria and fungi | We used the dataset published by (Madin et al., 2020) to characterise bacterial communities.<br>For bacteria, trait data was used at genus level when available. If no genus-level data was available, it was extrapolated to the Order level if the values of all genera within the order were consistent (mean > standard deviation for quantitative traits, and > 60% of all genera sharing one trait value for qualitative traits). If the data was not consistent within the order, it was kept as missing values. Then, cell volume was estimated as (cell radius)^2*3.14*cell length and log-transformed before further analysis. Other traits such as doubling time were | |

| | | |
|---|---|---|
| | considered but had very low coverage (e.g. for doubling time median of OTUs with available data over all plots < 20%) and were excluded from the analysis. Community weighted means were calculated using OTUs numbers as an approximation for relative abundances.<br>We complete these traits with community-levels characteristics. For bacteria, we quantified the relative abundance (approximated as OTU number) of groups commonly classified as oligotrophs (Acidobacteria, Verrucomicrobia, Planctomycetes) to copiotrophs (Actinobacteria, Betaproteobacteria, Gammaproteobacteria, Proteobacteria, Bacteroidetes).<br>For fungi, we calculated the relative abundance of pathotroph fungi among all fungi.<br>Finally, we also measured the PLFA-based fungal-bacteria ratio from 2011 and 2014; data that were first published by (Boeddinghaus et al., 2019). This was done by sampling two g of field moist soil (from the same composite samples as described above) for lipid extraction and fractionation following the alkaline methylation method described in (Frostegård et al., 1991). Samples were measured by gas chromatography (AutoSystem XL. PerkinElmer Inc., Massachusetts, USA) using a flame ionization detector, an HP-5 capillary column and helium as the carrier gas. Fatty acid nomenclature used was described by (Å Frostegård et al., 1993). Total bacterial PLFAs were calculated as the sum of Gram positive (a15:0, i15:0, i16:0 and i17:0) and Gram negative (cy17:0 and cy19:0)(Ruess and Chamberlain, 2010) plus the FAME 16:1ω?7 which is widespread in bacteria in general. Fungal biomass was represented by the PLFA 18:2ω?6.,9(Ruess and Chamberlain, 2010) | |
| Protists | Genus-level protist trait data was extracted from (Dumack et al., 2020) and completed for size class data by K. Dumack. Size classes were coded numerically: 1 (<10micron), 2 (11-30microns), 3 (31-50microns), 4 (>51 microns). Trophic levels were classified as plant parasites, bacterivores, or secondary consumers (non-plant parasites, eukaryvores, omnivores).<br>Primary consumers (plant parasites) had only one trait combination (size class 1) so we characterised these communities only by the relative abundance of plant parasites among all protists. | Bonkowski, Dumack, Fiore-Donno<br>Ref: (Dumack et al., 2020) |

**Methods Table 4.** Methods for the acquisition of environmental covariates.

| Environmental covariate | Methods | Owners / references |
|---|---|---|

| | | |
|---|---|---|
| pH and soil texture | Composite samples were taken in 2011 in all plots, by mixing 14 mineral topsoil samples (0–10 cm, using a split tube manual soil corer with 5 cm diameter). Ten g of sieved and air-dried soil were mixed with 25 ml 0.01 M CaCl2 solution and shook for 2 hours. Afterwards the pH of the soil suspension was measured using a glass electrode. The pH of each sample was measured twice (pH 1 and pH 2). We determined soil texture by separating soil particles into sand (2–0.063 mm), silt (0.063–0.002 mm) and clay (<0.002 mm) by sieving and sedimentation (DIN-ISO 11277). | Schöning, Trumbore, Schrumpf<br><br>Open datasets from Biodiversity Exploratories: Schöning et al., 2015a, Schöning et al., 2015b, 2013 |
| TWI | The topographic Wetness Index (TWI) combines measures of upslope contributing area (determining the amount of water received from upslope areas) and slope (determining the loss of water from the site to downslope areas), and has been shown in previous analyses to be a better predictor than local humidity measures. It is defined as ln(a/tanB), where a is the specific catchment area (cumulative upslope area which drains through a Digital Elevation Model (DEM, http://www.bkg.bund.de) cell, divided by per unit contour length) and tanB is the slope gradient in radians calculated over a local region surrounding the cell of interest 88,89. TWI was calculated from raster DEM data with a cell size of 25 m for all plots, using GIS tools (flow direction and flow accumulation tools of the hydrology toolset and raster calculator). The TWI measure used was the average value for a 4 × 4 window centred on the plot, i.e. 16 DEM cells corresponding to an area of 100 m × 100 m. | Le Provost, Manning<br>Ref: (Le Provost et al., 2021b)<br><br>Dataset from the Exploratories: Le Provost et al., 2021a |
| Mean annual temperature | Temperature measured 2 mm above ground level in each plot, then aggregated at the year level and averaged between 2008 and 2015. | Biodiversity Exploratories Instrumentation team<br><br>Open datasets from Biodiversity Exploratories: Hänsel et al., 2019 |

**Methods Table 5.** Measurement of ecosystem functions. Most data was extracted from an aggregated ecosystem functions datasets from the Biodiversity Exploratories *(Schenk et al., 2021)*; original datasets and their current status (publicly available or not yet) are listed below.

| Function "bundle" | Function name | Methods | Data owners and methods |
|---|---|---|---|
| Nitrogen cycle | DEA (denitrification enzyme activity) | Soil samples were taken from each plot in the beginning of May 2011 as a mixed sample from 14 soil cores of the top horizon (0-10 cm). | Berner, Boeddinghaus, Kandeler, Marhan,<br>Ref: (Keil et al., 2015; Smith and Tiedje, 1979) |

| | | | |
|---|---|---|---|
| | | Denitrification enzyme activity (DEA) was measured according to (Keil et al., 2015; Smith and Tiedje, 1979). | Open datasets from Biodiversity Exploratories: Kandeler et al., 2017a |
| | Activity of urease | Soil samples were taken from each plot in the beginning of May 2011 as a mixed sample from 14 soil cores of the top horizon (0-10 cm).<br><br>Urease activity was determined by incubating 1 g of fresh soil with 1.5 ml of 0.08 M substrate (urea) solution at 37 °C for 2 h (Schinner et al., 1996). Released ammonium was extracted with 12 ml of a 1 M potassium chloride/0.01 M hydrochloric solution and determined by a modified Berthelot reaction. | Berner, Boeddinghaus, Kandeler, Marhan,<br>Ref: (Schinner et al., 1996)<br><br>Open datasets from Biodiversity Exploratories: Kandeler et al., 2017a |
| | amoA_AOA (abundance of ammonia oxidation gene of archaea) | Soil samples were taken from each plot in 2011 and 2014 as a mixed sample from 14 soil cores of the top horizon (0-10 cm).<br>The abundance of the different functional genes (nifH, amoA) is quantified via real-time qPCR analysis and averaged across years | Kovacevic, Schloter, Schulz, Stempfhuber<br>Ref: (Barbara Stempfhuber et al., 2014)<br><br>Open datasets from Biodiversity Exploratories: Kovacevic et al., 2019; Stempfhuber and Schloter, 2017 |
| | amoA_AOB (abundance of ammonia oxidation gene of bacteria) | Soil samples were taken from each plot in 2011 and 2014 as a mixed sample from 14 soil cores of the top horizon (0-10 cm).<br>The abundance of the different functional genes (nifH, amoA) is quantified via real-time qPCR analysis and averaged across years | Schloter, Stempfhuber<br>Ref: (Barbara Stempfhuber et al., 2014)<br><br>Open datasets from Biodiversity Exploratories: Kovacevic et al., 2019; Stempfhuber and Schloter, 2017 |
| | nifH (abundance of nitrogen fixation gene in soil bacteria) | Soil samples were taken from each plot in 2011 as a mixed sample from 14 soil cores of the top horizon (0-10 cm).<br>The abundance of the different functional genes (nifH, amoA) is quantified via real-time qPCR analysis. and averaged across years | Schloter, Stempfhuber<br><br>Open datasets from Biodiversity Exploratories: Stempfhuber and Schloter, 2017 |
| | nxrA Nitrobacter-like NOB (abundance of nitrite oxidation gene of Nitrobacterbacteria) | Soil samples were taken from each plot in 2014 as a mixed sample from 14 soil cores of the top horizon (0-10 cm).<br>The abundance of ammonia nitrite oxidizing Nitrobacterbacteria and archaea was estimated using respective amoA gene, while NS-like and NB-like NOBs were targeted by primer sets for 16S rRNA genes for NS and nxrA primers genes specific for NB(Ollivier et al., 2013). | Kovacevic, Schulz, Schloter, Stempfhuber<br><br>Ref: (Ollivier et al., 2013)<br><br>Open datasets from Biodiversity Exploratories: Kovacevic et al., 2019 |
| | 16S rRNA Nitrospira-like NOB (abundance of nitrite oxidizing bacteria) | Soil samples were taken from each plot in 2014 as a mixed sample from 14 soil cores of the top horizon (0-10 cm).<br>The abundance of nitrite oxidizing Nitrospira was estimated using *16S rRNA gene primer* specific for Nitrospira(Ollivier et al., 2013). | Kovacevic, Schulz, Schloter, Stempfhuber<br><br>Ref: (Ollivier et al., 2013) |

| | | | |
|---|---|---|---|
| | | | Open datasets from Biodiversity Exploratories: Kovacevic et al., 2019 |
| | Potential_nitrification (nitrite accumulation over time) | Soil samples were taken from each plot in 2011 and 2014 as a a mixed sample from 14 soil cores of the top horizon (0-10 cm).<br>Ammonium and nitrate concentrations are measured after CaCl2 extraction, Potential nitrification is determined according to (Hoffmann et al., 2007) and averaged across years. | Stempfhuber, Schloter<br>Ref: (Hoffmann et al., 2007)<br><br>Open datasets from Biodiversity Exploratories: Stempfhuber and Schloter, 2017, 2016 |
| Decomposition | Dung decomposition (rate of dung removal by dung beetles) | Five dung samples were placed on each site in 2014. We used dung with a fresh weight of approx. 220.7 (± 19.9) g of cow, 34.4 (± 3.8) g of horse, 50.5 (± 3.6) g of sheep, 32.6 (± 1.6) g of deer, 14.5 (± 1.4) g of fox and 47.6 (± 2.4) g of wild boar. All dung samples have been placed on cellulose paper. After 48 h dung samples of removal experiments were collected, transferred into small paper bags, labelled (date, site-ID, dung type) and stored in a freezer at −20 °C. Removal samples were transferred into drying ovens and kept there at 60 °C for at least five days. Afterwards the dry weight for each dung sample was weighed (Mettler Toledo "EL 2001" (± 0.01 g), Columbus, Ohio) and noted for further calculations. | Frank, Blüthgen<br><br>Ref: (Frank et al., 2017)<br>Open datasets from Biodiversity Exploratories: Frank and Blüthgen, 2017 |
| | Root.decomposition | Decomposition measured of fine roots (<2 mm) within the upper 10 cm of the mineral soil in 2012. with standardized herbaceous roots. Root material was let to decompose in litterbags with mesh size of 100 micrometers µm during 6 months (until October 2012), and mass loss was determined.<br>Standardised herbaceous roots were used as control material. | Solly, Schrumpf, Klötzing, Schöning<br><br>Open datasets from Biodiversity Exploratories: Solly and Schöning, 2013 |
| | Litter.decomposition | Around 1.5 g of dry plant material collected in each EPsplot in spring 2012 were put back in their original location in January 2013 inside the climatic station. The above ground vegetation has been removed, and bags were fixed to the ground with nails or wood sticks. 10 replicated bags in each plot, 5 of them have been collected two months later, and the 5 last ones two more months later. The dry biomass remaining in each bag were measured to calculate daily decomposition rates. | Fischer, Grassein<br><br>Open datasets from Biodiversity Exploratories: Fischer and Grassein, 2015 |
| Carbon cycle | Xylosidase | Activity of the soil enzyme beta-xylosidase (hemicellulose degradation). Soil samples were taken from each plot in the beginning of May 2011 as a a mixed sample from 14 soil cores of the top horizon (0-10 cm). Activities were determined according to Marx et al. (2001) as described in detail in (Berner et al., 2011), using fluorescent 4-methylumbelliferone substrates (4-MUF; Sigma-Aldrich, St. Louis, USA) and a buffered solution (pH 6.1). | Berner, Kandeler, Marhan, Boeddinghaus<br>Ref: (Berner et al., 2011; Marx et al., 2001)<br><br>Open datasets from Biodiversity Exploratories: Kandeler et al., 2017a |
| | N-Acetyl beta Glucosaminidase | Activity of the soil enzyme beta-N-acetylglucosaminidase (chitin degradation). Soil samples were taken from each plot in the beginning of May 2011 as a a mixed sample from | Berner, Kandeler, Marhan, Boeddinghaus<br>Ref: (Berner et al., 2011; Marx et al., 2001) |

| | | | |
|---|---|---|---|
| | | 14 soil cores of the top horizon (0-10 cm). Activities were determined according to (Marx et al., 2001) as described in detail in (Berner et al., 2011), using fluorescent 4-methylumbelliferone substrates (4-MUF; Sigma-Aldrich, St. Louis, USA) and a buffered solution (pH 6.1). | Open datasets from Biodiversity Exploratories: Kandeler et al., 2017a |
| | Beta Glucosidase | Activity of the soil enzyme beta-glucosidase (cellulose degradation). Soil samples were taken from each plot in the beginning of May 2011 as a a mixed sample from 14 soil cores of the top horizon (0-10 cm). Activities were determined according to (Marx et al., 2001) as described in detail in (Berner et al., 2011), using fluorescent 4-methylumbelliferone substrates (4-MUF; Sigma-Aldrich, St. Louis, USA) and a buffered solution (pH 6.1). | Berner, Kandeler, Marhan, Boeddinghaus<br>Ref: (Berner et al., 2011; Marx et al., 2001)<br><br>Open datasets from Biodiversity Exploratories: Kandeler et al., 2017a |
| Respiration | Respiration | Soil respiration was measured from late June to July in 2018 and 2019, using the soda-lime adsorption absorption method with an open and static chamber to determine soil CO2 efflux. Soda-lime, i.e. mainly Ca(OH)2 and NaOH, was used as the adsorption absorption material. We installed four chambers, forming a square of 10 m side length (around the weather station), and one bottom-sealed trap served as control. The mass of soda-lime for each measurement was ~12 g/d and the exposure time was three days. Aboveground vegetation was carefully clipped and removed from the installation area and PVC rings was were plugged down to 1 cm soil depth. Soda-lime and lidstraps were installed five to six days after vegetation clipping. Just before installation, we re-wetted the soda-lime to compensate for the initial moisture content of about 18%, since CO2 needs to be hydrated before reacting with soda-lime. The $CO_2$ flux is calculated from the dry soda-lime mass differences considering i) the exposure time, ii) the area of the chamber, iii) the coefficient of 1.69 to account for the water lost during the reaction and iv) after correcting with controls (bottom-sealed) traps. The soil efflux is calculated by the equation (Grogan, 1998; Keith and Wong, 2006): Rs = [(WGsample-WGblank)/CA] × [24/T] × [12/44] ×1.69 where Rs is the soil respiration in [$gCO_2$-C $m^{-2}$ $d^{-1}$], WG is the weight gain [g], CA is the chamber basal area in [$m^{-2}$], T is the implementation time in [h], 12/44 is the ratio of Carbon atomic mass over Carbon dioxide molecular mass and 1.69 compensates for the $H_2O$ formed during $CO_2$ sorption and lost during drying. | Apostolakis, Schöning, Schrumpf, Klötzing, Trumbore<br><br>Refs: (Grogan, 1998; Keith and Wong, 2006)<br><br>Open datasets from Biodiversity Exploratories: Apostolakis et al., 2020 |
| Biomass | Plant aboveground biomass production | On two subplots of 1.5 m x 2 m size (varying position every year between 2009-2017), aboveground biomass was harvested between mid-May and mid-June by clipping the vegetation at a height of 5 cm. Samples were June, dried by 80 °C for 48 hours and weighted. On each subplots the inside of the frame - thrown 4 times by chance on each subplot - was harvested. After drying and weighting, an arithmetic mean of biomass per m2 for each EP plot was calculated. Data was then averaged across years<br>2009-2017. | Boch, Fischer, Minker, Prati, Schäfer<br><br>Open datasets from Biodiversity Exploratories: Boch et al., 2017; Fischer et al., 2017; Prati et al., 2013; Schäfer et al., 2018, 2016a, 2016b; Schmitt et al., 2012a, 2012b, 2011 |

**Supplementary results**

**Table S 1.** Trait-specific hypothesis testing: expected response of each trait to resource availability (fertilisation) and disturbance (mowing/grazing); overall coverage (number of individual with available trait data / total number of individual for each plot, median (min-max)); number of taxa with data available / total number of taxa; Expected correlation with land-use intensity; test of the hypothesised response. P-values were corrected for multiple testing (false detection rate, n = 47).

Empty cells indicate that no specific response to the corresponding driver is expected.
Number of taxa with available trait data is considered after extrapolation (see Methods Table 3). Total number of taxa includes taxa identified only at higher level (e.g. Genus sp.).
*for bacteria, includes genera from which data was extrapolated from other genera in the same Order. Colours indicate trophic level (pale to dark colours) and position above (blue) or belowground (brown).

| Trophic guild | Trait | Expected response to resource availability (fertilisation) | Expected response to disturbance (mowing / grazing) | Plot-level coverage, i.e. proportion (in abundance) with species available trait data)<br><br>median (min-max) | Number of taxa with available trait data /number of taxa identified; number of plots with available data | Expectation: high trait values correspond to fast or slow strategies | Observed response to LUI (slope estimate; 95% confidence interval; pvalue | Response as expected? |
|---|---|---|---|---|---|---|---|---|
| Plants (primary producers) | Specific leaf area | High resource availability favours fast-growing species, with rapid resource capture and fast turnover of organs. This translates into high SLA, which allows rapid growth and high resource acquisition but short-lived leaves and hence poor conservation of resources (Grime, 1979; Lavorel and Garnier, 2002; Wright et al., 2004). | | 99 (61-100) | Taxa: 283/362<br><br>Plots: 150/150 | Fast | 0.31 (0.17 - 0.46)<br><br>$P < 0.001$ | Yes |
| | Seed mass | | Seed mass reflects a trade-off between colonisation ability | 99 (61-100) | Taxa: 334/362 | Slow | -0.31 (-0.44 – -0.18) | Yes |

| | | | | | | | | |
|---|---|---|---|---|---|---|---|---|
| | | | and seedling survival (Díaz et al., 2016; Lavorel and Garnier, 2002). Colonisation ability is particularly important at high disturbance to allow for recolonisation. Further, low seed mass is usually associated with short plant height (Díaz et al., 2016) which is selected by grazing and mowing disturbance. | | Plots: 150/150 | | P < 0.001 | |
| | Leaf dry matter content | LDMC is a correlate of specific leaf area (negative correlation), associated with slow growth and good resource conservation. It is a negatively associated with soil fertility (Hodgson et al., 2011). | | 100 (61-100) | Taxa: 330/362<br><br>Plots: 150/150 | Slow | -0.28 (-0.42 – -0.13)<br><br>P < 0.001 | Yes |
| | Leaf nitrogen | High resource availability translates into high leaf nutrient content of fast-growing species (Lavorel and Garnier, 2002). | | 98 (61-100) | Taxa: 252/362<br><br>Plots: 150/150 | Fast | 0.46 (0.32 – 0.59)<br><br>P < 0.001 | Yes |
| | Leaf phosphorus | High resource availability translates into high leaf nutrient content of fast-growing species (Lavorel and Garnier, 2002). | | 97 (57-100) | Taxa: 197/362<br><br>Plots: 150/150 | Fast | 0.51 (0.38 – 0.64)<br><br>P < 0.001 | Yes |
| | Root tissue density | Low resource availability favours slow-growing roots with slow turnover that better conserve nutrients (Bergmann et al., 2020; Weigelt et al., 2021). | | 95 (27-100) | Taxa: 231/362<br>Plots: 150/150 | Slow | -0.22 (-0.37 – -0.07)<br><br>P = 0.006 | Yes |
| Lepidoptera (primary consumers, aboveground) | Flight period | | An early emergence (i.e. long flight season) allows for growth and reproduction before the start of the | 100 (83-100) | Taxa: 90/97<br><br>Plots: 136/150 | Fast | 0.31 (0.21 – 0.42)<br><br>P < 0.001 | Yes |

| | | | | | | | | |
|---|---|---|---|---|---|---|---|---|
| | | | disturbance (mowing) (Börschig et al., 2013). | | | | | |
| | Generations per year | | High reproductive rates can compensate for mortality due to disturbance and favour fast recolonisation (Börschig et al., 2013).<br>In addition disturbance selects for early flowering plants which provide resources earlier in the year and promote multiple generations per year (Börschig et al., 2013). | 100 (83-100) | Taxa: 88/97<br><br>Plots: 136/150 | Fast | 0.21 (0.06 - 0.37)<br><br>P = 0.014 | Yes |
| | Hibernation stage | | A later hibernation stage allows individuals to reproduce before early disturbances and better recolonise a habitat (Börschig et al., 2013). | 100 (83-100) | Taxa: 90/97<br><br>Plots: 136/150 | Fast | 0.27 (0.15 – 0.39)<br><br>P < 0.001 | Yes |
| | Size (wing size) | | Larger wings translate into higher dispersal abilities (which facilitates recolonisation) but larger body sizes makes it more difficult to survive disturbance (Birkhofer et al., 2017; Hanson et al., 2016). | 100 (83-100) | Taxa: 90/97<br><br>Plots: 136/150 | Fast or slow | -0.07 (-0.22 – 0.08)<br><br>P = 0.42 | Inconclusive |
| | Generalism | Fertilisation reduces plant diversity (Le Provost et al., 2021b; Socher et al., 2012), which tends to reduce host availability for specialist species (Chisté et al., 2018). | | 95 (75-100) | Taxa: 79/97<br><br>Plots: 136/150 | Fast | 0.30 (0.15 – 0.45)<br><br>P = 0.001 | Yes |
| Arthropods (primary | Body size | Large resource availability promotes fast pace of life (r strategy), which is usually related | Smaller body size makes it easier to hide and escape the disturbance (Birkhofer et al., | 100 (0.99-100) | Taxa: 797/803 | Slow | -0.24 (-0.4 – -0.09)<br><br>P = 0.004 | Yes |

| | | | | | | | | |
|---|---|---|---|---|---|---|---|---|
| consumers, aboveground) | | to smaller body size (Pianka, 1970). | 2017, 2015b; Simons et al., 2016). | | Plots: 150/150 | | | |
| | Feeding generalism | Fertilisation reduces plant diversity (Le Provost et al., 2021b; Socher et al., 2012), which tends to reduce host availability for specialist species (Chisté et al., 2018). | Generalists are expected to respond less strongly to disturbance in terms of land-use intensity (Simons et al., 2016). | 95 (61-99) | Taxa: 621/803<br><br>Plots: 150/150 | Fast | 0.34 (0.21 – 0.47)<br><br>P < 0.001 | Yes |
| | Dispersal ability | | Higher dispersal ability makes it easier to recolonise after disturbance (Birkhofer et al., 2017, 2015b; Simons et al., 2016). | 100 (0.99-100) | Taxa: 784/803<br><br>Plots: 150/150 | Fast | 0.37 (0.25 – 0.49)<br><br>P < 0.001 | Yes |
| | Generations per year | More resources might allow more generations if resources are limiting. | Faster reproduction makes it easier to recover from disturbance | 83 (41-98) | Taxa: 202/803<br><br>Plots: 150/150 | Fast | 0.56 (0.45 – 0.67)<br><br>P < 0.001 | Yes |
| Arthropods (secondary consumers, aboveground) | Body size | | Smaller body size makes it easier to hide and escape the disturbance (Birkhofer et al., 2017, 2015b; Simons et al., 2016). | 100 (100-100) | Taxa: 240/240<br><br>Plots: 150/150 | Slow | 0.03 (-0.12 – 0.19)<br><br>P = 0.76 | Inconclusive |
| | Dispersal ability | | Higher dispersal ability makes it easier to recolonise after disturbance (Birkhofer et al., 2017, 2015b; Simons et al., 2016). | 100 (0.76-100) | Taxa: 239/240<br><br>Plots: 150/150 | Fast | 0.33 (0.19 – 0.47)<br><br>P < 0.001 | Yes |
| Birds (secondary tertiary consumers) | Body mass | | Mowing decreases the abundance of large invertebrates on average, but creates resource "flushes" after just after mowing when invertebrate prey are easier to find, resulting in higher foraging efficiency after | 100 (100-100) | Taxa: 24/24<br><br>Plots: 145/150 | Slow | 0.23 (0.08 – 0.37)<br><br>P = 0.005 | No |

| | | | | | | | | |
|---|---|---|---|---|---|---|---|---|
| | | | mowing (Devereux et al., 2006) but overall higher resource availability disparities in time.<br>In addition, mowing also increases mortality and destroys nests of ground-nesting species (Frawley and Best, 1992; MacDonald, 2006).<br>Both mechanisms should promote 'fast' species which are able to recover after disturbance and to rapidly exploit available resources; these are characterised by rapid reproductive traits, typically associated to small size due to energy allocation (Sibly et al., 2012). | | | | | |
| | Incubation time | | | 100 (100-100) | Taxa: 24/24<br><br>Plots: 145/150 | Slow | 0.12 (-0.03 – 0.27)<br><br>P = 0.20 | Inconclusive |
| | Maximum number of offspring | | | 100 (100-100) | Taxa: 24/24<br><br>Plots: 145/150 | Fast | -0.12 (-0.27 – 0.03)<br><br>P = 0.17 | Inconclusive |
| | Generation length | | | 100 (100-100) | Taxa: 24/24<br><br>Plots: 145/150 | Slow | 0.19 (0.04 – 0.34)<br><br>P = 0.023 | No |
| Bats (tertiary consumers, aboveground) | Body mass | | Mowing decreases the abundance of large invertebrates on average, but creates resource "flushes" after just after mowing when | 100 (100-100) | Taxa: 11/11<br><br>Plots: 148/150 | Slow | -0.07 (-0.18 – 0.03)<br><br>P = 0.25 | Inconclusive |

| | | | | | | | | |
|---|---|---|---|---|---|---|---|---|
| | | | invertebrate prey are easier to find, resulting in higher foraging efficiency after mowing (Devereux et al., 2006) but overall higher resource availability disparities in time. Both mechanisms should promote 'fast' species which are able to recover after disturbance and to rapidly exploit available resources. | | | | | |
| | Maximum longevity | | | 100 (62-100) | Taxa: 10/11<br><br>Plots: 148/150 | Slow | -0.01 (-0.12 – 0.11)<br><br>P = 0.91 | Inconclusive |
| | Number of offspring | | | 100 (62-100) | Taxa: 10/11<br><br>Plots: 148/150 | Fast | -0.06 (-0.17 – 0.05)<br><br>P = 0.39 | Inconclusive |
| Protists (plant pathogens i.e. primary consumers) | Relative abundance | High resource availability promotes fast-growing plants which have lower amounts of constitutive defences than slow-growing species; and fast-growing species support higher herbivory rates than slow-growing species (Endara and Coley, 2011). This is likely to promote high abundance of herbivores and pathogens. Empirical evidence supports this; protist pathogens have been shown to respond positively to intensive management (Fiore-Donno et al., 2020b). | | non applicable | Taxa: 9 genera<br><br>Plots: 150/150 | Fast | 0.46 (0.32 – 0.59)<br><br>P < 0.001 | Yes |

| | | | | | | | | |
|---|---|---|---|---|---|---|---|---|
| Bacteria and fungi | Bacterial cell volume | Alternative hypotheses: a. small cells are more efficient for diffusive uptake; or b. for a given substrate demand, large radius compensates low substrate concentrations (Westoby et al., 2021). | | 42 (27-72) | Taxa: 371/1155*<br><br>Plots: 150/150 | fast (a) or slow (b) | -0.30 (-0.41 – -0.19)<br><br>P < 0.001 | Yes (hypothesis B) |
| | Bacterial oligotroph:copiotroph ratio | By definition, oligotrophic bacteria survive better at low resource availability, while copiotrophic bacteria reproduce faster at high resources availability (Barnett et al., 2021; Fierer et al., 2012; Leff et al., 2015). | | 50 (29-70) | Taxa: 827/1155*<br><br>Plots: 150/150 | slow | -0.16 (-0.27 – -0.03)<br><br>P = 0.022 | Yes |
| | Bacterial genome size | Bacteria with large genomes are capable to use a larger array of resources in low amounts; thus they are more ecologically successful in environments where resources are scarce but diverse and where there is little penalty for slow growth (Konstantinidis and Tiedje, 2004; Leff et al., 2015). | | 54 (40-79) | Taxa: 685/1155*<br><br>Plots: 150/150 | slow | -0.15 (-0.22 – -0.08)<br><br>P < 0.001 | Yes |
| | Fungi:bacteria ratio | Fungi dominate in soils with low resource availability and are associated with "slow" plant communities (Boeddinghaus et al., 2019; de Vries et al., 2012, 2006). | | (non applicable) | Taxa: non applicable<br><br>Plots: 150/150 | slow | -0.28 (-0.40 – -0.19)<br><br>P < 0.001 | Yes |
| | Proportion of fungal pathotrophs among all fungi | High resource availability promote fast-growing plants which have lower amounts of constitutive defences than slow-growing species; and fast-growing species support higher herbivory rates than slow-growing species (Endara and Coley, 2011) which promotes fungal pathogens (Lekberg et al., 2021; Liu et al., 2021). | | (non applicable) | Taxa: non applicable<br><br>Plots: 150/150 | fast | 0.37 (0.24 – 0.50)<br><br>P < 0.001 | Yes |

| | | | | | | | | |
|---|---|---|---|---|---|---|---|---|
| | | Alternatively, pathogens at high intensity might derive from organic amendments such as slurry. | | | | | | |
| Protists (bacterivores) | Only one trait, cell size | Low resource availability promotes slow turnover and conservative strategies with larger cell size in protists (Cavalier-Smith, 1980; Lüftenegger et al., 1985). Effect might be indirect via bacterial abundance. | | 85 (54-98) | Taxa: 26/30 genera<br><br>Plots: 150/150 | slow | -0.29 (-0.44 – -0.15)<br><br>$P < 0.001$ | Yes |
| Protists (secondary consumers) | Only one trait, cell size | Low resource availability promotes slow turnover and conservative strategies with larger cell size in protists (Cavalier-Smith, 1980; Lüftenegger et al., 1985). Effect might be indirect via bacterial and other protists's abundance. | | 98 (79-100) | Taxa: 31/32 genera<br><br>Plots: 150/150 | slow | -0.39 (-0.51 – -0.26)<br><br>$P < 0.001$ | Yes |
| Arthropods (primary consumers, belowground) | Body size | Large resource availability promotes fast pace of life (r strategy), which is usually related to smaller body size (Pianka, 1970). Effect might be indirect through consumption of plant roots. | In most arthropods, smaller body size makes it easier to hide and escape the disturbance (Birkhofer et al., 2017, 2015b; Simons et al., 2016). | 100 (100-100) | Taxa: 109/109<br><br>Plots: 136/150 | slow | -0.24 (-0.41 – -0.11)<br><br>$P = 0.002$ | Yes |
| | Generalism | Fertilisation reduces plant diversity (Le Provost et al., 2021b; Socher et al., 2012), which tends to reduce host availability for specialist species (Chisté et al., 2018). | | 100 (92-100) | Taxa: 107/109<br><br>Plots: 111/150 | fast | -0.04 (-0.23 – 0.14)<br><br>$P = 0.74$ | Inconclusive |
| | Dispersal ability | | High dispersal ability makes it easier to evade disturbance and to recolonise after disturbance | 100 (92-100) | Taxa: 107/109 | fast | 0.15 (-0.02 – 0.31)<br><br>$P = 0.12$ | Inconclusive |

| | | | | | | | | |
|---|---|---|---|---|---|---|---|---|
| | | | (Birkhofer et al., 2017, 2015b; Simons et al., 2016). | | Plots: 136/150 | | | |
| Collembola (omnivores, belowground) | Body size | Large resource availability promotes fast pace of life (r strategy), which is usually related to smaller body size (Pianka, 1970). Effect might be indirect through consumption of plant roots and lower trophic levels. | In most arthropods, smaller body size makes it easier to hide and escape the disturbance (Birkhofer et al., 2017, 2015b; Simons et al., 2016). | 100 (91-100) | Taxa: 63/64<br><br>Plots: 140/150 | Slow | 0.02 (-0.13 -- 0.18)<br><br>P = 0.84 | Inconclusive |
| | Depth preference | Species dwelling in deeper horizons are usually considered to have 'faster' traits due to resource availability and soil pore size constraining body size (V. Wolters, and R. Saifutdinov., pers. comm and (Petersen, 1980)) | | 100 (91-100) | Taxa: 63/64<br><br>Plots: 140/150 | Fast | -0.01 (-0.16 – 0.14)<br><br>P = 0.93 | Inconclusive |
| | Voltinism | | Fast reproduction (multivoltine species) allows for more reproduction before disturbance and rapid recolonisation after disturbance. | 100 (0-100) | Taxa: 55/64<br><br>Plots: 138/150 | Fast | -0.09 (-0.24 – 0.06)<br><br>P = 0.34 | Inconclusive |
| | Reproduction type: sexual | | Parthenogenetic species are usually considered as "r" strategists (Petersen, 1980), good at colonising new territories, with especially large population sizes at early succession stages (Chauvat et al., 2007). Indeed, thelytoky (parthenogenetic reproduction of only females) is an advantage for colonization, since there is no need of energy for partner searching. | 100 (91-100) | Taxa: 63/64<br><br>Plots: 140/150 | Slow | 0.00 (-0.15 -- 0.16)<br><br>P = 0.96 | Inconclusive |
| | Habitat specificity | More specialised species (in order of specialisation: non-specialised, | | 63 (21-100) | Taxa: 51/56 | slow | -0.07 (-0.23 -- 0.09) | Inconclusive |

| | | | | | | | | |
|---|---|---|---|---|---|---|---|---|
| Oribatid mites (omnivores, belowground) | | soil, surface, litter) are usually considered to have 'faster' traits (V. Wolters, and R. Saifutdinov., pers. comm and (Petersen, 1980)) | | | Plots: 129/150 | | P = 0.49 | |
| | Feeding specialisation | Low resource availability favours high relative fungal availability and species richness, providing resources for fungivorous species which are the most specialised (V. Wolters and A Zaytsev, pers. comm). | | 63 (21-100) | Taxa: 51/56<br><br>Plots: 129/150 | slow | | Inconclusive |
| | Reproduction type: sexual | | As for Collembola, parthenogenetic species are considered as faster than sexually reproducting species due to higher colonisation abilities and faster reproduction. | 63 (21-100) | Taxa: 51/56<br><br>Plots: 129/150 | slow | 0.00 (-0.14 – 0.15)<br><br>P = 0.96 | Inconclusive |
| | Days to maturity | | Shorter maturation time are related to faster reproduction, which makes it easier to recolonise after a disturbance. | 63 (21-100) | Taxa: 51/56<br><br>Plots: 129/150 | slow | -0.05 (-0.21 – 0.1)<br><br>P = 0.60 | Inconclusive |
| | Body mass | Large resource availability promotes fast pace of life (r strategy), which is usually related to smaller body size (Pianka, 1970). Effect might be indirect through consumption of plant roots and lower trophic levels. | In most arthropods, smaller body size makes it easier to hide and escape the disturbance (Birkhofer et al., 2017, 2015a; Simons et al., 2016). | 63 (21-100) | Taxa: 51/56<br><br>Plots: 129/150 | slow | -0.17 (-0.30 – -0.03)<br><br>P = 0.03 | Yes |
| Other arthropods (secondary consumers, belowground) | Body size | | In most arthropods, smaller body size makes it easier to hide and escape the disturbance (Birkhofer et al., 2017, 2015a; Simons et al., 2016). | 100 (100-100) | Taxa: 205/205<br><br>Plots: 150/150 | slow | -0.02 (-0.16 – 0.1)<br><br>P = 0.76 | Inconclusive |

| | | | | | | | | |
|---|---|---|---|---|---|---|---|---|
| | Dispersal ability | | Higher dispersal ability makes it easier to recolonise after disturbance (Birkhofer et al., 2017, 2015a; Simons et al., 2016). | 100 (100-100) | Taxa: 205/205<br><br>Plots: 150/150 | fast | 0.09 (-0.06 – 0.24)<br><br>P = 0.32 | Inconclusive |

**Table S 2.** Identification of species and guild-level slow-fast axes. For taxa-level PCA, missing traits values were imputed using the mice package (mean method). Community-level PCAs only include sites for which data was available, and taxa with no trait data were usually removed (see Methods Table 3 for a few cases where we imputed trait data). Land -use intensity, fertilisation and disturbance (combined index of grazing and mowing) were added as supplementary variables.

| Guild | Species-level PCA | Community-level PCA | Comment |
|---|---|---|---|
| Vascular plants | 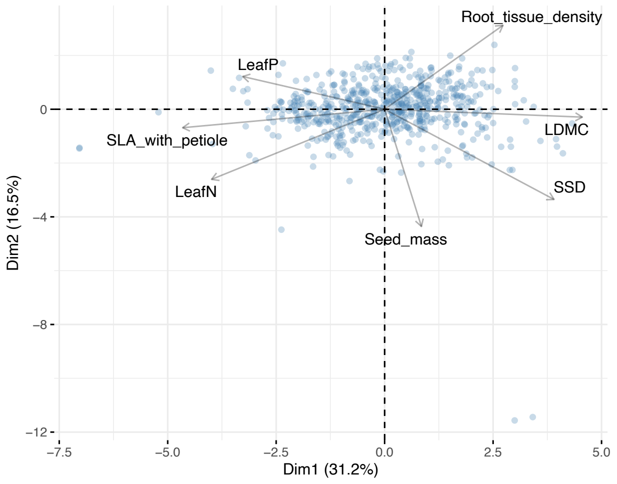  | 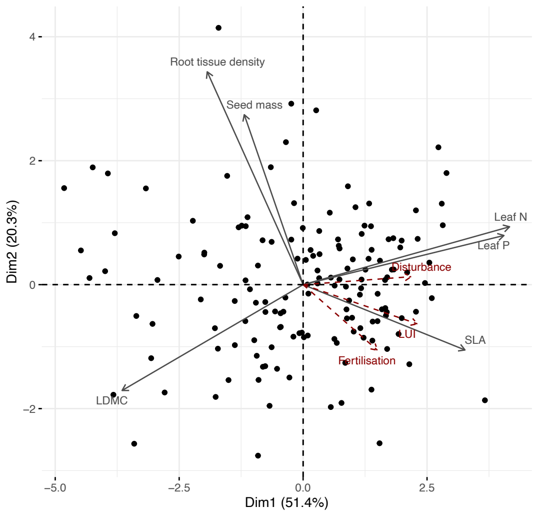  | |

| | | | |
|---|---|---|---|
| Lepidoptera | 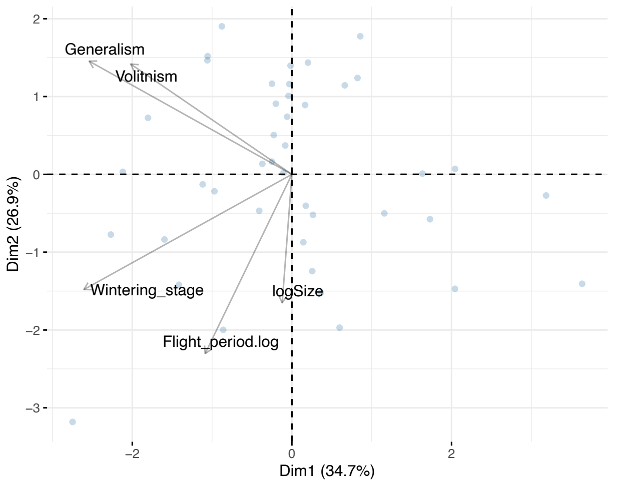 | 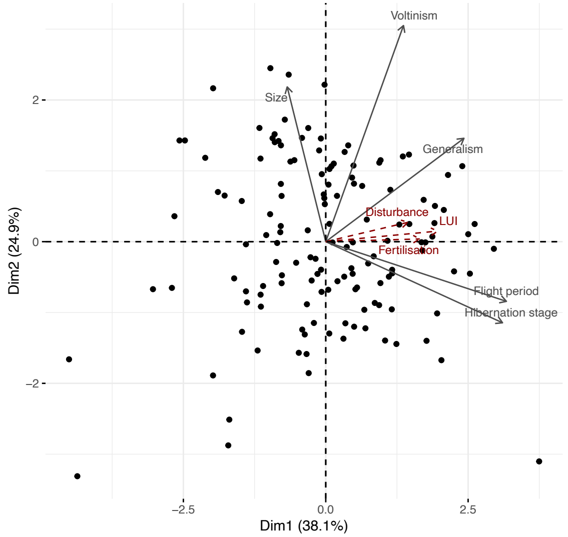 | Partial: unexpected response from body size. Body size was measured as wing length, which is an indicator of overall body size (which is expected to be a "slow" trait; smaller size helps to survive disturbance) but also of dispersal ability ("fast" trait, as larger wings promote recolonisation after disturbance). Both effects might cancel each other leading to no response of size. |
| Arthropods (primary consumers, aboveground) | 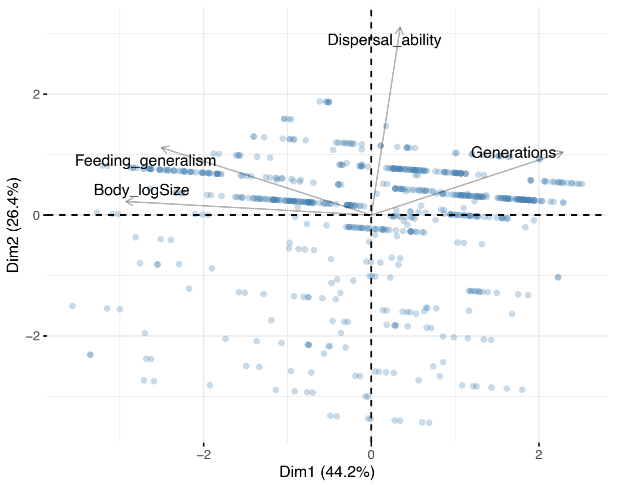 | 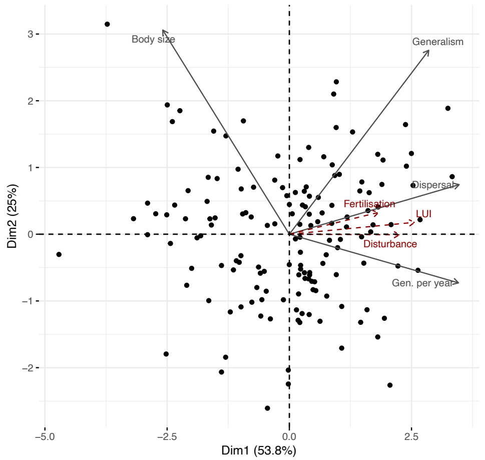 | |

<table>
<tr>
<td>Arthropods (secondary consumers, aboveground)</td>
<td>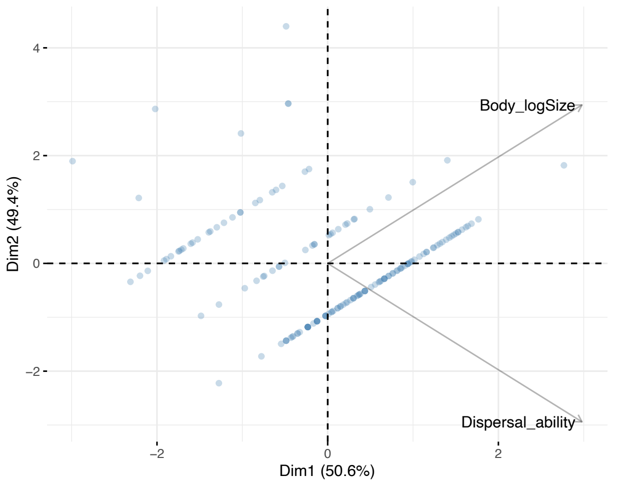
</td>
<td>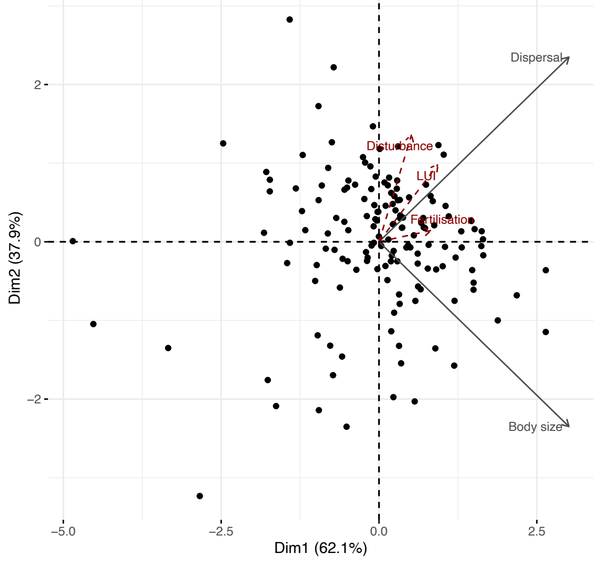
</td>
<td>Dispersal and body size are slightly confounded because larger body size increases dispersal abilities. However, these two traits are opposed on the second axis of the PCA, which we use as our slow-fast index.</td>
</tr>
<tr>
<td>Birds (tertiary consumers)</td>
<td>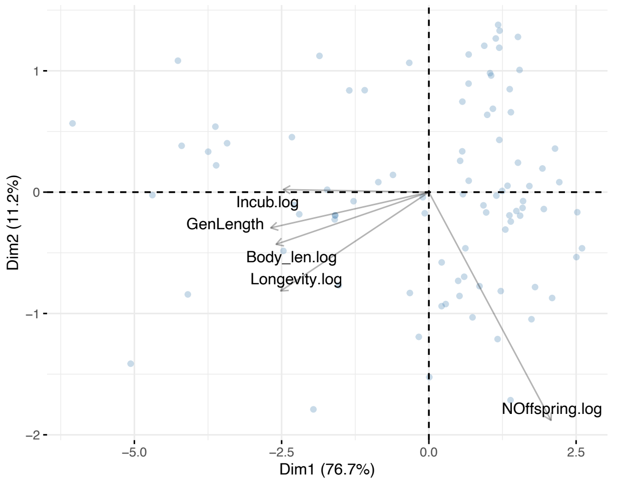
</td>
<td>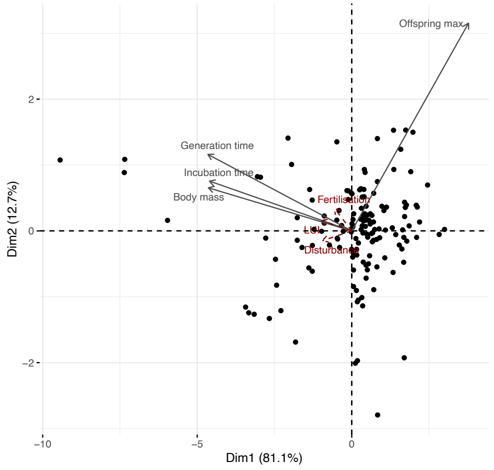
</td>
<td></td>
</tr>
</table>

| | | | |
|---|---|---|---|
| Bats (tertiary consumers) | 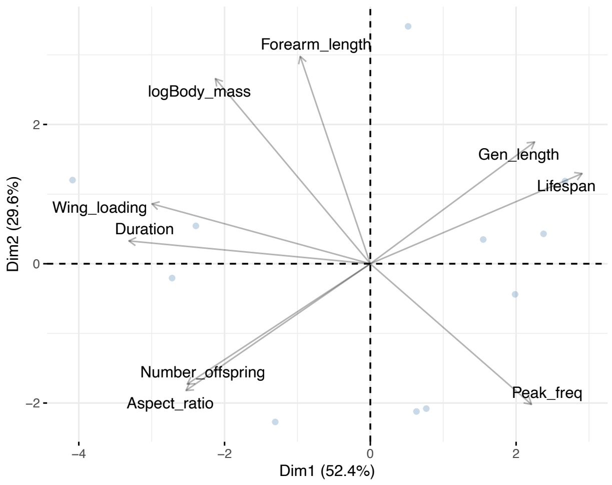  | 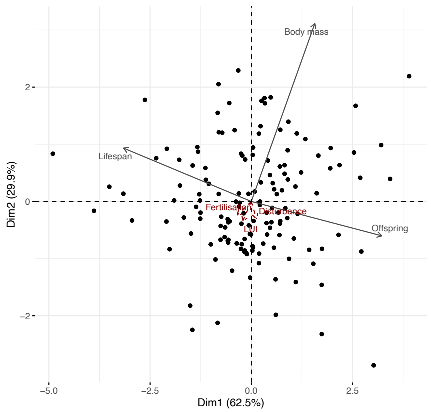  | Partial<br><br>Body mass is usually considered as a 'slow' trait, and is expected to be positively correlated to lifespan and negatively to the number of offspring (trade-off between survival and reproduction).<br>Hibernation saves resources and leads, in hibernating bats (all species observed in our study), to a correlation between number of offspring and body mass (Wilkinson 2002), leading to the results observed here. |
| Protists (plant pathogens) | Not applicable: only one trait | Not applicable: only one trait | |

| | | |
|---|---|---|
| Micro-organisms (bacteria and fungi) | Not applicable: mix of species- and community- level traits | 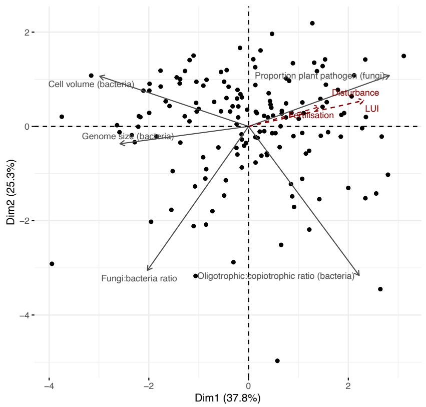 |
| Protists (bacterivores) | Not applicable: only one trait | Not applicable: only one trait |
| Protists (secondary consumers) | Not applicable: only one trait | Not applicable: only one trait |
| Arthropods (primary consumers, belowground) | 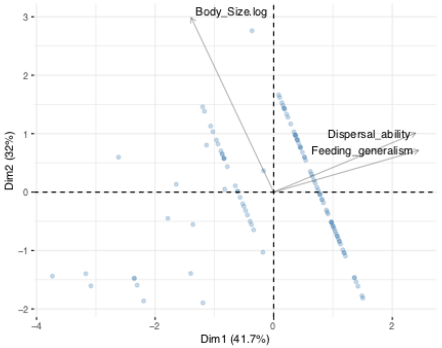 | 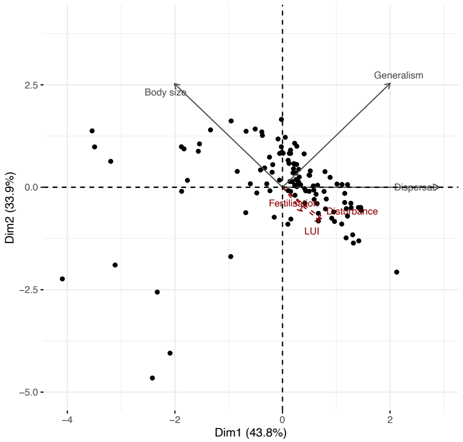 |

| | | |
|---|---|---|
| Collembola | 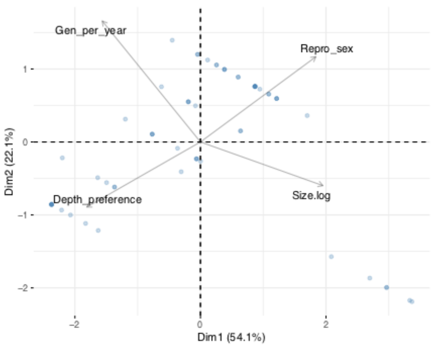 | 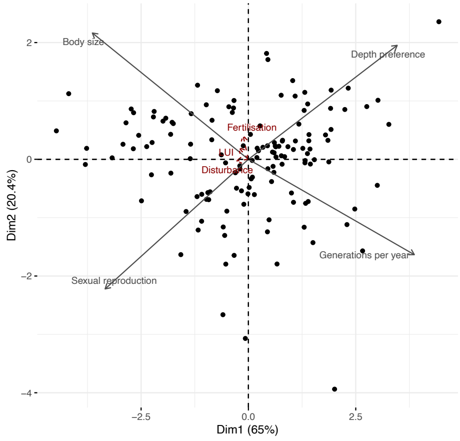 |
| Oribatid mites | 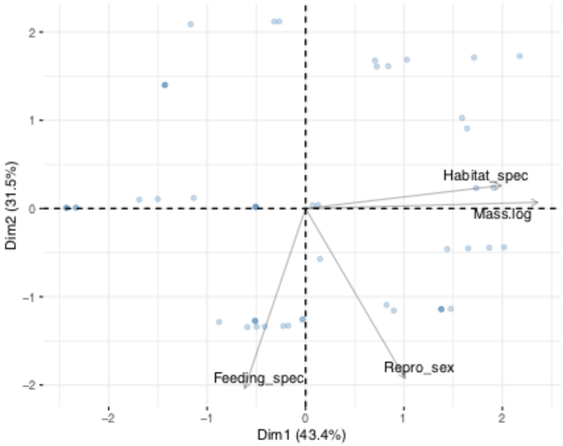 | 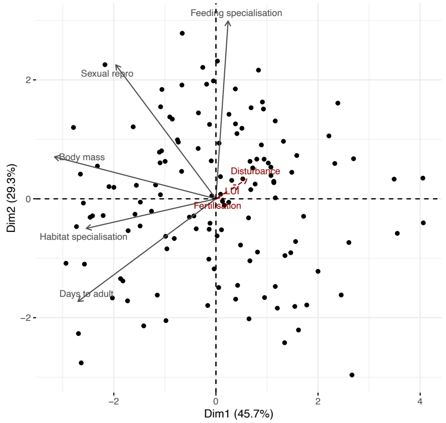 |

| | | |
|---|---|---|
| Arthropods (secondary consumers, belowground) | 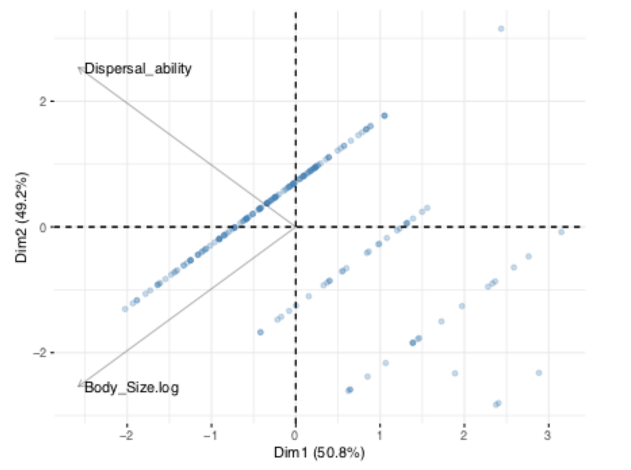  | 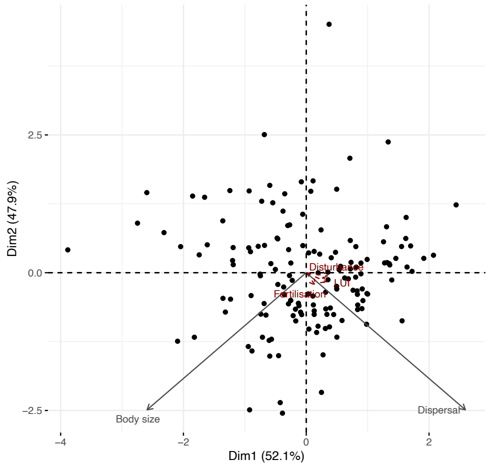  |

**Table S 3.** SEM path parameters for the belowground model (fitted with lavaan, boostrapped with 300 iterations). Colours indicate trophic level (pale to dark colours).

| **Fit indices** | P-value = 0.98 | RMSEA = 0.00 | CFI = 1.00 | BIC = 2441 | |
|---|---|---|---|---|---|
| **Regression Slopes** | | | | | |
| **Left-hand side variable** | **Right-hand side variable** | **Estimate** | **Standard error** | **z** | **p** |
| Plant slow-fast axis | ~ land use intensity | 0.51 | 0.07 | 6.91 | .000 |
| Above-ground arthropods (primary consumers) slow-fast axis | ~ land use intensity | 0.50 | 0.07 | 7.36 | .000 |
| Above-ground arthropods (primary consumers) slow-fast axis | ~ Plant slow-fast axis | 0.21 | 0.07 | 2.97 | .003 |
| Lepidoptera (primary consumers) slow-fast axis | ~ land use intensity | 0.34 | 0.09 | 3.81 | .000 |

| | | | | | |
|---|---|---|---|---|---|
| Lepidoptera (primary consumers) slow-fast axis | ~ Plant slow-fast axis | 0.29 | 0.09 | 3.40 | .001 |
| Above-ground arthropods (secondary consumers) slow-fast axis | ~ land use intensity | 0.09 | 0.12 | 0.72 | .473 |
| Above-ground arthropods (secondary consumers) slow-fast axis | ~ Plant slow-fast axis | 0.00 | 0.09 | 0.04 | .965 |
| Above-ground arthropods (secondary consumers) slow-fast axis | ~ Lepidoptera (primary consumers) slow-fast axis | 0.09 | 0.11 | 0.80 | .421 |
| Above-ground arthropods (secondary consumers) slow-fast axis | ~ Above-ground arthropods (primary consumers) slow-fast axis | 0.20 | 0.11 | 1.93 | .053 |
| Birds (tertiary consumers) slow-fast axis | ~ land use intensity | -0.25 | 0.11 | -2.39 | .017 |
| Birds (tertiary consumers) slow-fast axis | ~ Plant slow-fast axis | -0.11 | 0.09 | -1.18 | .237 |
| Birds (tertiary consumers) slow-fast axis | ~ Lepidoptera (primary consumers) slow-fast axis | -0.07 | 0.10 | -0.63 | .530 |
| Birds (tertiary consumers) slow-fast axis | ~ Above-ground arthropods (primary consumers) slow-fast axis | 0.24 | 0.13 | 1.87 | .061 |
| Birds (tertiary consumers) slow-fast axis | ~ Above-ground arthropods (secondary consumers) slow-fast axis | 0.00 | 0.08 | 0.03 | .975 |
| Bats (tertiary consumers) slow-fast axis | ~ land use intensity | 0.01 | 0.12 | 0.12 | .905 |
| Bats (tertiary consumers) slow-fast axis | ~ Plant slow-fast axis | 0.16 | 0.11 | 1.46 | .144 |
| Bats (tertiary consumers) slow-fast axis | ~ Lepidoptera (primary consumers) slow-fast axis | -0.01 | 0.11 | -0.09 | .931 |
| Bats (tertiary consumers) slow-fast axis | ~ Above-ground arthropods (primary consumers) slow-fast axis | -0.23 | 0.10 | -2.30 | .022 |
| Bats (tertiary consumers) slow-fast axis | ~ Above-ground arthropods (secondary consumers) slow-fast axis | -0.05 | 0.09 | -0.54 | .591 |
| **Intercepts** | | | | | |

| | | | | | |
|---|---|---|---|---|---|
| Plant slow-fast axis | | -0.00 | 0.07 | -0.00 | 1.000 |
| Above-ground arthropods (primary consumers) slow-fast axis | | 0.00 | 0.06 | 0.00 | 1.000 |
| Lepidoptera (primary consumers) slow-fast axis | | -0.00 | 0.07 | -0.00 | .996 |
| Above-ground arthropods (secondary consumers) slow-fast axis | | 0.00 | 0.08 | 0.00 | 1.000 |
| Birds (tertiary consumers) slow-fast axis | | -0.00 | 0.08 | -0.03 | .977 |
| Bats (tertiary consumers) slow-fast axis | | -0.00 | 0.09 | -0.02 | .983 |
| LUI | | 0 (fixed parameter) | | | |
| **Residual Variances** | | | | | |
| Plant slow-fast axis | | 0.73 | 0.08 | 8.99 | .000 |
| Above-ground arthropods (primary consumers) slow-fast axis | | 0.60 | 0.07 | 9.06 | .000 |
| Lepidoptera (primary consumers) slow-fast axis | | 0.67 | 0.08 | 7.91 | .000 |
| Above-ground arthropods (secondary consumers) slow-fast axis | | 0.89 | 0.13 | 6.83 | .000 |
| Birds (tertiary consumers) slow-fast axis | | 0.91 | 0.22 | 4.23 | .000 |
| Bats (tertiary consumers) slow-fast axis | | 0.94 | 0.15 | 6.49 | .000 |
| LUI | | 1 (fixed parameter) | | | |
| **Residual Variances** | | | | | |
| Birds (tertiary consumers) slow-fast axis | Bats (tertiary consumers) slow-fast axis | 0.02 | 0.07 | 0.25 | .805 |

**Table S 4.** SEM path parameters for the belowground model (fitted with lavaan, boostrapped with 300 iterations). Colours indicate trophic level (pale to dark colours).

| **Fit indices** | P-value = 0.76 | RMSEA = 0.00 | CFI = 1.00 | BIC = 3243 | |
|---|---|---|---|---|---|
| **Regression Slopes** | | | | | |
| **Left-hand side variable** | **Right-hand side variable** | **Estimate** | **Standard error** | **z** | **p** |
| Plant slow-fast axis | ~ land use intensity | 0.51 | 0.07 | 7.48 | .000 |
| Protists (pathotrophs. i.e.e primary consumers) slow-fast axis | ~ land use intensity | 0.31 | 0.08 | 3.65 | .000 |
| Protists (pathotrophs. i.e.e primary consumers) slow-fast axis | ~ Plant slow-fast axis | 0.34 | 0.08 | 4.10 | .000 |
| Bacteria and fungi slow-fast axis | ~ land use intensity | 0.34 | 0.06 | 5.71 | .000 |
| Bacteria and fungi slow-fast axis | ~ Plant slow-fast axis | 0.41 | 0.09 | 4.54 | .000 |
| Protists (bacterivores) slow-fast axis | ~ land use intensity | 0.41 | 0.10 | 4.19 | .000 |
| Protists (bacterivores) slow-fast axis | ~ Plant slow-fast axis | 0.13 | 0.10 | 1.30 | .193 |
| Protists (bacterivores) slow-fast axis | ~ Bacteria and fungi slow-fast axis | -0.29 | 0.07 | -3.98 | .000 |
| Protists (secondary consumers) slow-fast axis | ~ land use intensity | 0.26 | 0.10 | 2.61 | .009 |
| Protists (secondary consumers) slow-fast axis | ~ Plant slow-fast axis | 0.03 | 0.12 | 0.23 | .821 |
| Protists (secondary consumers) slow-fast axis | ~ Bacteria and fungi slow-fast axis | 0.30 | 0.12 | 2.42 | .015 |
| Protists (secondary consumers) slow-fast axis | ~ Protists (bacterivores) slow-fast axis | 0.04 | 0.08 | 0.57 | .566 |
| Oribatid mites (omnivores) slow-fast axis | ~ land use intensity | 0.04 | 0.14 | 0.32 | .748 |
| Oribatid mites (omnivores) slow-fast axis | ~ Plant slow-fast axis | 0.10 | 0.10 | 1.04 | .300 |
| Oribatid mites (omnivores) slow-fast axis | ~ Bacteria and fungi slow-fast axis | 0.10 | 0.11 | 0.98 | .325 |

| | | | | | |
|---|---|---|---|---|---|
| Oribatid mites (omnivores) slow-fast axis | ~ Protists (bacterivores) slow-fast axis | -0.03 | 0.08 | -0.33 | .742 |
| Oribatid mites (omnivores) slow-fast axis | ~ Protists (secondary consumers) slow-fast axis | -0.05 | 0.10 | -0.51 | .610 |
| Collembola (omnivores) slow-fast axis | ~ land use intensity | -0.04 | 0.12 | -0.31 | .759 |
| Collembola (omnivores) slow-fast axis | ~ Plant slow-fast axis | 0.15 | 0.11 | 1.38 | .167 |
| Collembola (omnivores) slow-fast axis | ~ Bacteria and fungi slow-fast axis | -0.22 | 0.12 | -1.79 | .073 |
| Collembola (omnivores) slow-fast axis | ~ Protists (bacterivores) slow-fast axis | 0.08 | 0.08 | 1.01 | .311 |
| Collembola (omnivores) slow-fast axis | ~ Protists (secondary consumers) slow-fast axis | 0.01 | 0.11 | 0.10 | .924 |
| Arthropods (secondary consumers) slow-fast axis | ~ land use intensity | 0.17 | 0.08 | 2.04 | .041 |
| Arthropods (secondary consumers) slow-fast axis | ~ Oribatid mites slow-fast axis | 0.15 | 0.09 | 1.72 | .085 |
| Arthropods (secondary consumers) slow-fast axis | ~ Collembola slow-fast axis | -0.07 | 0.08 | -0.99 | .324 |
| Arthropods (secondary consumers) slow-fast axis | ~ Protists (bacterivores) slow-fast axis | -0.13 | 0.08 | -1.69 | .091 |
| Arthropods (secondary consumers) slow-fast axis | ~ Protists (secondary consumers) slow-fast axis | -0.13 | 0.08 | -1.65 | .098 |
| **Intercepts** | | | | | |
| Plant slow-fast axis | | -0.00 | 0.07 | -0.00 | 1.000 |
| Protists (primary consumers) slow-fast axis | | 0.00 | 0.06 | 0.00 | 1.000 |
| Bacteria and fungi fast-slow axis | | 0.00 | 0.06 | 0.00 | 1.000 |
| Protists (bacterivores) slow-fast axis | | 0.00 | 0.07 | 0.00 | 1.000 |
| Protists (secondary consumers) slow-fast axis | | -0.00 | 0.07 | -0.00 | 1.000 |
| Oribatid mites (omnivores) slow-fast axis | | 0.02 | 0.09 | 0.18 | .858 |
| Collembola (omnivores) slow-fast axis | | -0.01 | 0.08 | -0.08 | .935 |

| | | | | | |
|---|---|---|---|---|---|
| Above-ground arthropods (secondary consumers) slow-fast axis | | -0.00 | 0.09 | -0.03 | .973 |
| LUI | | 0 (fixed parameter) | | | |
| **Residual Variances** | | | | | |
| Plant slow-fast axis | | 0.73 | 0.08 | 9.37 | .000 |
| Protists (primary consumers) slow-fast axis | | 0.68 | 0.07 | 9.73 | .000 |
| Bacteria and fungi fast-slow axis | | 0.57 | 0.09 | 6.57 | .000 |
| Protists (bacterivores) slow-fast axis | | 0.85 | 0.11 | 7.67 | .000 |
| Protists (secondary consumers) slow-fast axis | | 0.73 | 0.07 | 10.18 | .000 |
| Oribatid mites (omnivores) slow-fast axis | | 0.95 | 0.11 | 8.95 | .000 |
| Collembola (omnivores) slow-fast axis | | 0.95 | 0.13 | 7.20 | .000 |
| Above-ground arthropods (secondary consumers) slow-fast axis | | 0.93 | 0.14 | 6.80 | .000 |
| LUI | | 1 (fixed parameter) | | | |
| **Residual Variances** | | | | | |
| Protists (primary consumers) slow-fast axis | Bacteria and fungi fast-slow axis | 0.16 | 0.05 | 3.10 | .002 |
| Protists (primary consumers) slow-fast axis | Above-ground arthropods (secondary consumers) slow-fast axis | 0 (fixed parameter) | | | |

**Table S 5.** Comparison of the effect of multiple drivers on the ecosystem functions slow-fast axis, obtained from linear models with the function slow-fast axis as a response and the indicated variables as explanatory variables. P-values were corrected for multiple testing within these different model (i.e. correction for false discovery rates, R function p.adjust, n = 5).

| **Model** | **Slope estimate (+/- standard error)** | **P-value** | **Adj $R^2$** |
|---|---|---|---|
| Functions slow-fast axis ~ entire community slow-fast axis (based on guild-level PCA) | 0.39 (0.04) | < 0.001 | 33% |
| Functions slow-fast axis ~ entire community slow-fast axis (based on individual traits) | 0.49 (0.05) | < 0.001 | 38% |
| Functions slow-fast axis ~ plants slow-fast axis | 0.30 (0.05) | < 0.001 | 19% |
| Functions slow-fast ~ microbial slow-fast | 0.41 (0.06) | < 0.001 | 23% |
| Functions slow-fast ~ land-use intensity | 0.62 (0.09) | < 0.001 | 26% |
| Functions slow-fast ~ taxonomic multidiversity | -0.42 (0.09) | < 0.001 | 11% |

**Table S 6.** Variance partitioning of land use intensity and multivariate trait community weighted mean across sampling plot and years. The partitioning was done using the varpart function (package vegan). Only groups with more than one sampling year are included. For bacteria and fungi, year 2017 was excluded because only one trait was available (% pathogen fungi) which did not allow to properly partition the variance.

| **Variable** | **Variance attributed to the plot** | **Variance attributed to the year** | **Residuals** |
|---|---|---|---|
| LUI | 0.70 | <0.01 | 0.30 |
| Birds | 0.25 | 0.18 | 0.60 |
| Bats | 0.43 | <0.01 | 0.57 |
| Plants | 0.52 | 0.01 | 0.47 |

| **Variable** | **Variance attributed to the plot** | **Variance attributed to the year** | **Residuals** |
|---|---|---|---|
| Arthropods (above-ground, secondary consumers) | 0.09 | 0.01 | 0.90 |
| Arthropods (above-ground, primary consumers) | 0.23 | 0.04 | 0.72 |
| Bacteria and fungi | 0.68 | 0.03 | 0.24 |
| Protists (secondary consumers) | 0.22 | <0.01 | 0.78 |

| | | | | | | | |
|---|---|---|---|---|---|---|---|
| Arthropods (below-ground, secondary consumers) | 0.10 | 0.03 | 0.87 | Protists (bacterivores) | 0.15 | 0.08 | 0.68 |
| Arthropods (below-ground, primary consumers) | 0.17 | 0.06 | 0.79 | Protists (pland pathogens) | 0.38 | 0.15 | 0.3 |

**Figure S 1.** Correlation between guild-level PCA axes or single traits and land use intensity. Three axes were retained when more than two traits were available; otherwise only two were retained. For protists, only one trait was available, and the correlation between the CWM of this trait and LUI is shown. The fast-slow axis is always the first PC axis, except for arthropod (secondary consumers) above-ground, for which it was PC axis 2. The axes were transformed were needed (inversed sign) so that higher axis values indicate "faster" strategies. P-values were corrected for false detection rates (***: P < 0.001, **: P < 0.01, *: P< 0.05, n.s.: P > 0.05)

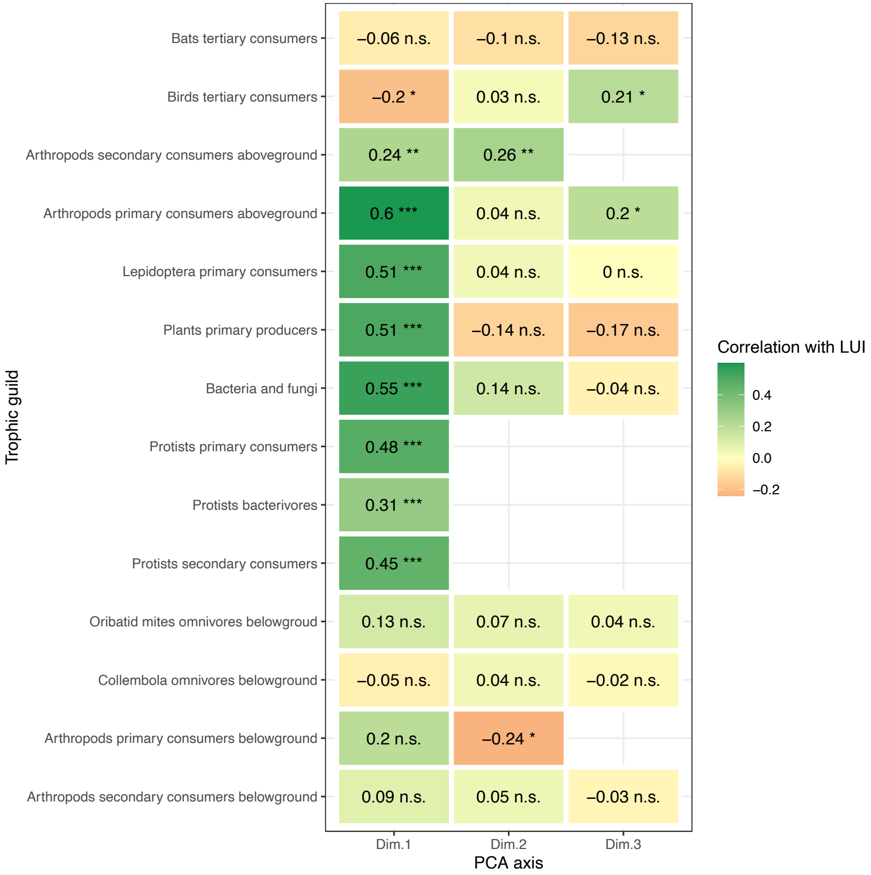

**Figure S 2.** Theoretical SEMs for above-and below ground trophic levels

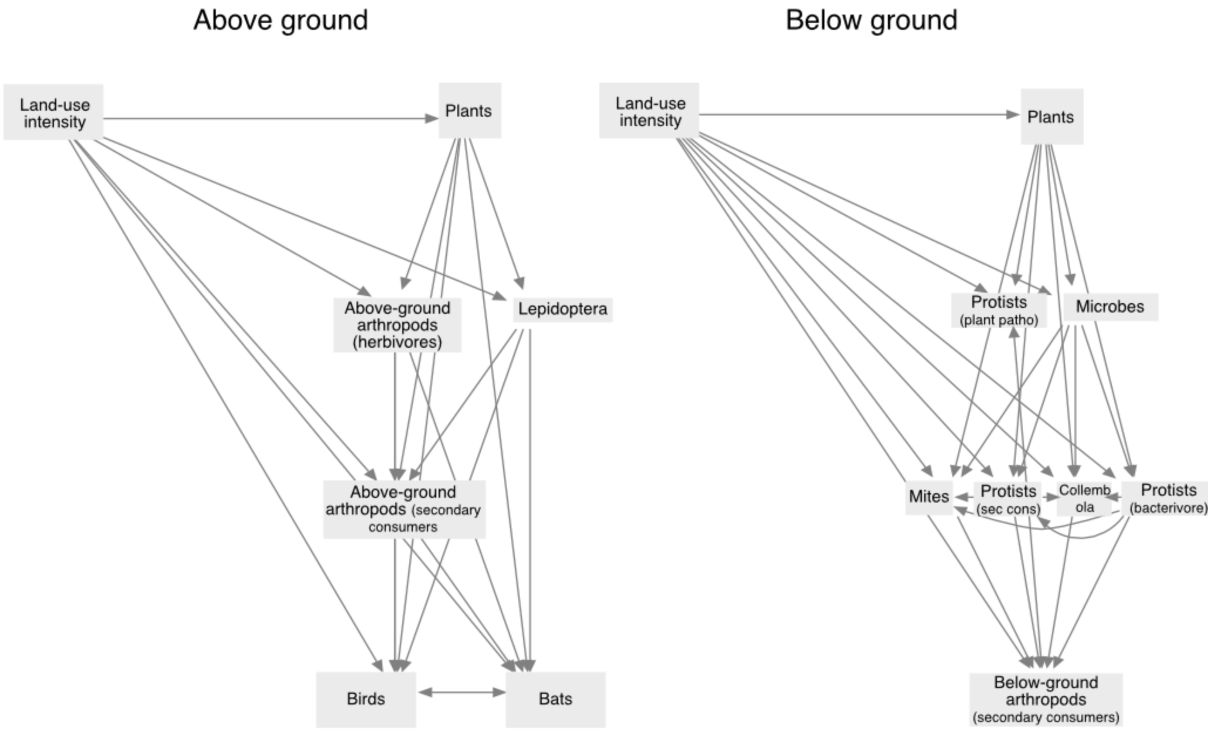

**Figure S 3.** Whole community level slow-fast axis, based on all traits. In contrast to the results shown in Fig. 2, where the PCA was conducted on guild-level slow-fast axes, the PCA was conducted on all traits CWM. Each trait was weighted as 1/n, with n the number of traits available for the group, so that all groups are weighted equally. Traits expected to be “slow” are coded in blue, “fast” in green

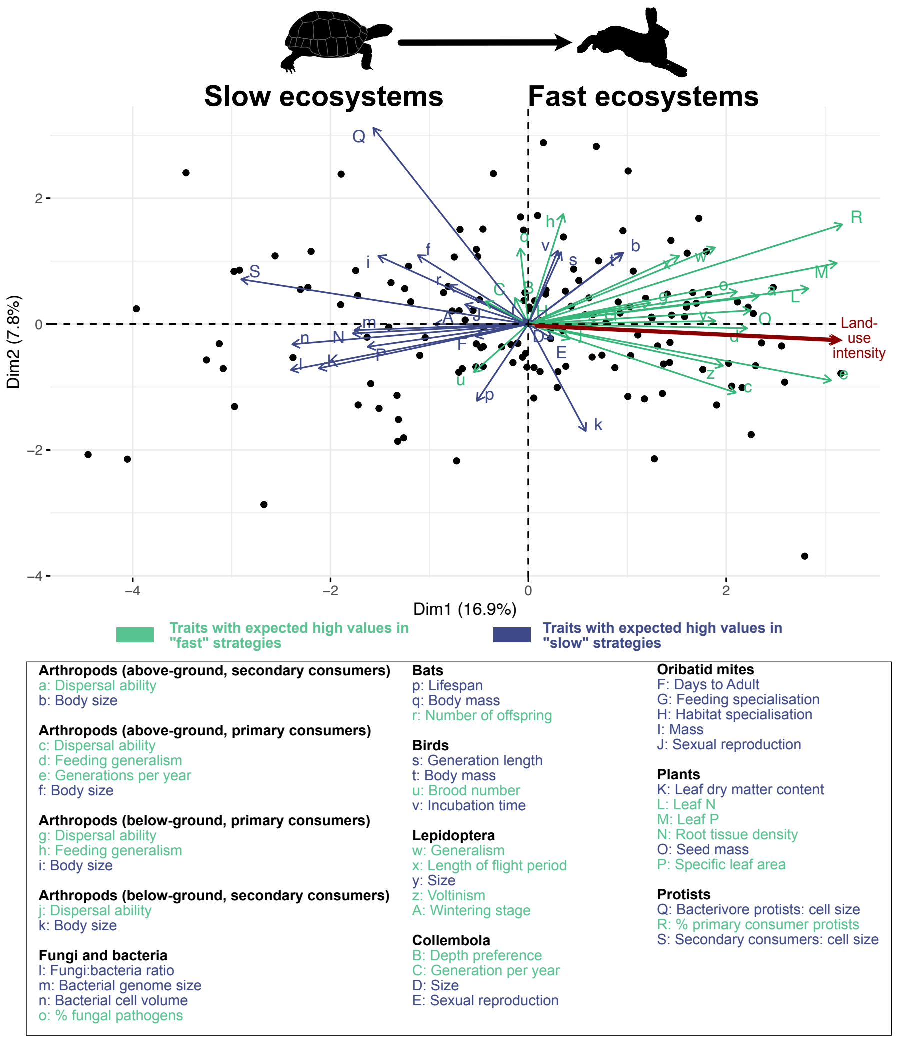

**Figure S 4.** Direct and indirect links between land- use intensity, the functional traits slow-fast axis and the ecosystem function slow-fast axis. **Functional trait slow-fast axes was measured as the first axis of a PCA with all selected traits as variables (Figure S 3, PC1), rather than individual guild slow-fast axes.**

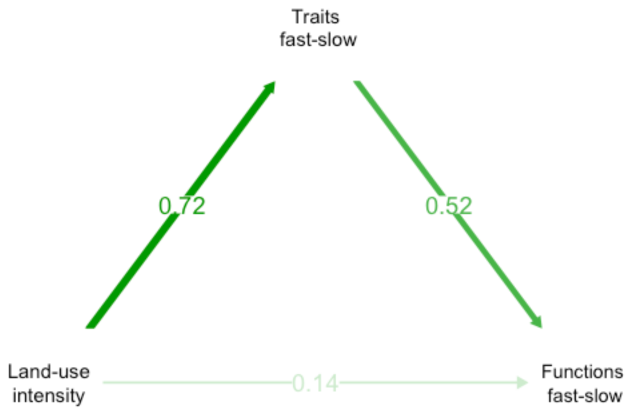

### Sensitivity analyses

We tested the sensitivity of our results to different parameters.

**Role of species turnover versus abundance gradients in driving the results**

In addition, we conducted a few additional analyses to assess whether our results were due to species turnover (i.e. species with certain sets of traits disappearing from one end of the land-use intensity gradient to the other) or driven mostly by changes in the abundance few species. For this, we first calculated the turnover and nestedness components of the abundance-based bray-Curtis beta diversity (package betapart, function beta.multi.abund). This was done both for the whole land-use intensity gradient (including all plots) and between the 10 highest-LUI and lowest-LUI plots. This indicated that for most guilds, taxonomic turnover was the main component of species abundance dissimilarities (**Table S 7**).
In addition, we also conducted all the analyses on non-weighted community trait values (i.e. all species present in each plot across the multiple sampling years were given equal abundance of 1). These analyses overall shower weaker, but consistent, results as shown in the main text (**Table S 8**, **Table S 9**, **Table S 10**, **Table S 11**, **Table S 12**, **Figure S 5**, **Figure S 6**, **Figure S 7**, **Figure S 8**). This indicates that both abundance changes and species turnover were responsible for the slow-fast community responses observed at the level of the entire community.

#### Correction for environmental covariates

We also ran the analyses without conducting the environmental corrections on the traits or functions. The community-level results were mostly similar to the main results, except for soil bacterial and fungal communities whose traits strongly differed across regions (**Table S 13**). The functioning of the system was also strongly affected by region (**Figure S 12**). However, the entire-community slow-fast axis was still identified (**Figure S 9**, **Figure S 10**) and also drove the ecosystem functions slow-fast axis, although less strongly than in the main results (**Figure S 13**, **Table S 14**, **Table S 17**). The structure SEM to quantify the relative importance of direct and trophically-mediated LUI effects on the different trophic levels was not as well supported by the data as in the main results, probably due to the effect of the environmental covariates that were not accounted for (**Figure S 11**, **Table S 15**, **Table S 16**).

#### Exclusion of body size data

While there were strong hypothetical reasons to expect body size to be a key trait driving fast-slow variation at the community level, it can respond to a range of drivers and drive life history variation in the absence of other trait responses. As a result, the effect of body mass is often removed before identifying life history trade-offs. We therefore conducted analyses in which we excluded all body size data (resulting in the exclusion of bacterivores and predator protists as other trait data were not available for these groups). We were still able to identify a strong guild-level slow-fast axis for most groups, except collembola. This resulted in somewhat weaker, but still consistent results regarding the synchrony of slow-fast axes across guilds and the effect of the whole community slow-fast axis on ecosystem functioning (**Figure S 14**, **Figure S 15**, **Table S 18**)

**Table S 7.** Abundance-based nestedness (balanced variation) and turnover (abundance gradient) components of Bray-Curtis dissimilarity. The components of dissimilarity were calculated either across all plots, or by first aggregating the 10 highest- and lowest-LUI plots and calculating the dissimilarity between the two groups. Dissimilarities were then calculated using the beta.multi.abund function (package betapart). Colours indicate trophic level (pale to dark colours) and position above (blue) or belowground (brown).

| **Guild** | **Dissimilarity across all plots** | | | **Dissimilarity between 10 highest- and lowest- intensity plots** | | |
|---|---|---|---|---|---|---|
| | Turnover (balanced variation) | Nestedness (abundance gradient) | Overall dissimilarity | Turnover (balanced variation) | Nestedness (abundance gradient) | Turnover (balanced variation) |
| Vascular plants (primary producers) | 0.98 | 0.002 | 0.983 | 0.861 | 0.029 | 0.890 |
| Lepidoptera (primary consumers) | 0.957 | 0.029 | 0.986 | 0.513 | 0.141 | 0.654 |

| | | | | | | |
|---|---|---|---|---|---|---|
| Other arthropods (primary consumers, aboveground) | 0.973 | 0.011 | 0.984 | 0.864 | 0.018 | 0.882 |
| Arthropods (secondary consumers, aboveground) | 0.971 | 0.014 | 0.984 | 0.727 | 0.126 | 0.852 |
| Birds (secondary consumers) | 0.968 | 0.020 | 0.987 | 0.767 | 0.060 | 0.826 |
| Bats | 0.912 | 0.073 | 0.985 | 0.518 | 0.019 | 0.537 |
| Protists (plant pathogens) | 0.884 | 0.106 | 0.990 | 0.894 | 0.034 | 0.928 |
| Micro-organisms (bacteria and fungi) | Non applicable (mix of community- and species-level traits) | | | Non applicable (mix of community- and species-level traits) | | |
| Protists (bacterivores) | 0.960 | 0.0196 | 0.980 | 0.676 | 0.041 | 0.717 |
| Protists (secondary consumers) | 0.955 | 0.025 | 0.980 | 0.710 | 0.145 | 0.855 |
| Arthropods (primary consumers, belowground) | 0.971 | 0.019 | 0.990 | 0.864 | 0.034 | 0.898 |
| Collembola (omnivores) | 0.910 | 0.017 | 0.987 | 0.622 | 0.169 | 0.790 |
| Oribatid mites (omnivores) | 0.964 | 0.021 | 0.985 | 0.622 | 0.169 | 0.790 |
| Arthropods (secondary consumers, belowground) | 0.973 | 0.012 | 0.984 | 0.812 | 0.049 | 0.861 |

**Table S 8.** Identification of guild-level slow-fast axes. **Community-level trait data (CWM) was not weighted by taxa abundance.**

| **Guild** | **Community-level PCA** | **Comment** |
|---|---|---|
| Vascular plants (primary producers) | 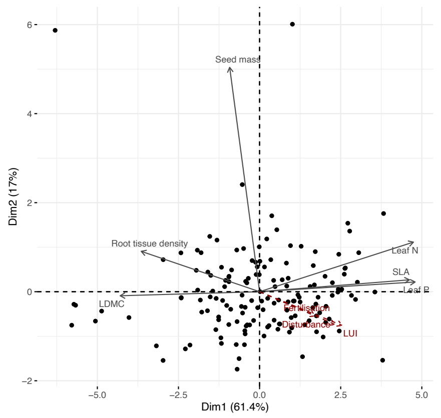  | |
| Lepidoptera (primary consumers) | 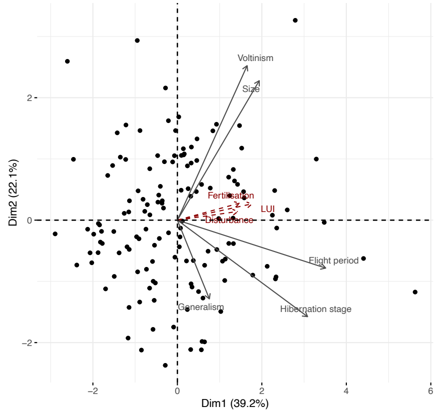  | Partial: no response from body size. Body size was measured as wing length, which is an indicator of overall body size (which is expected to be a "slow" trait; smaller size helps to survive disturbance) but also of dispersal ability ("fast" trait, as larger wings promote recolonisation after disturbance). Both effects might cancel each other leading to no response of size. |

| | | |
|---|---|---|
| Other arthropods (primary consumers, aboveground) | 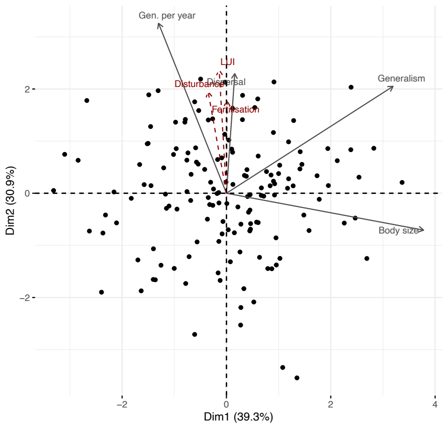 | The slow-fast axis is the second axis of the PCA |
| Arthropods (secondary consumers, aboveground) | 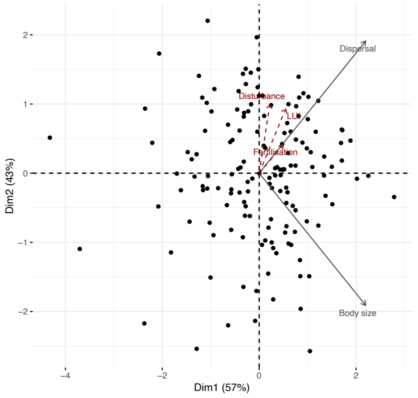 | Dispersal and body size are slightly confounded because larger body size increases dispersal abilities. However, these two traits are opposed on the second axis of the PCA, which we use as our slow-fast index. |

| | | |
|---|---|---|
| Birds (tertiary consumers) | 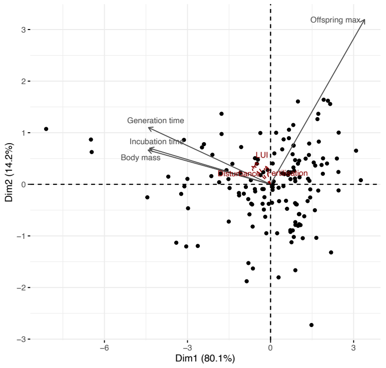 | |
| Bats (tertiary consumers) | 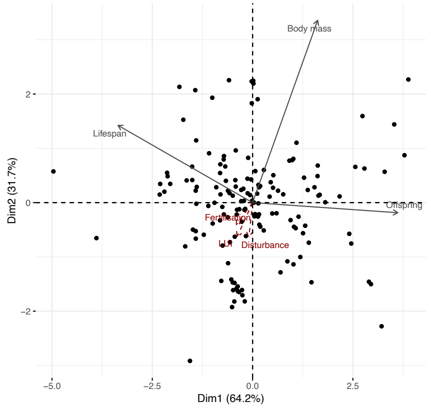 | Partial<br><br>Body mass is usually considered as a 'slow' trait, and is expected to be positively correlated to lifespan and negatively to the number of offspring (trade-off between survival and reproduction). Hibernation saves resources and leads, in hibernating bats (most of the species observed in our study), to a correlation between number of offspring and body mass (Wilkinson 2002), leading to the results observed here. |
| Protists (plant pathogens, ie primary consumers) | Only one trait | |

<table>
<tr><td>Bacteria and fungi</td><td>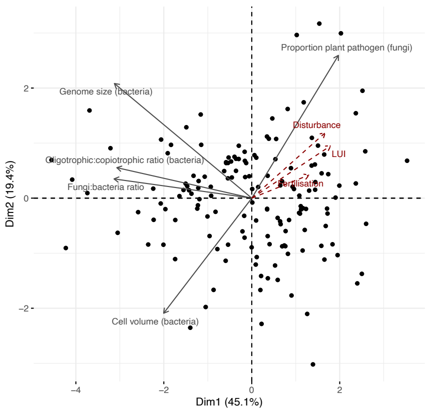
</td><td></td></tr>
<tr><td>Protists (bacterivores)</td><td>Only one trait</td><td></td></tr>
<tr><td>Protists (secondary consumers)</td><td>Only one trait</td><td></td></tr>
<tr><td>Arthropods (primary consumers, belowground)</td><td>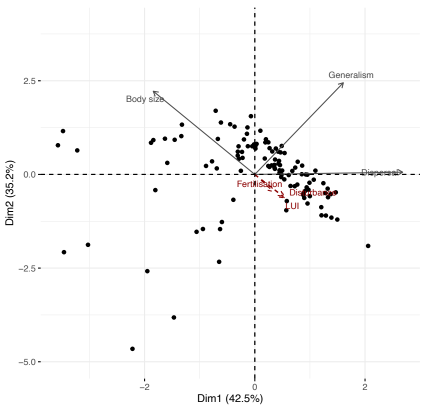
</td><td></td></tr>
</table>

| | |
|---|---|
| Collembola (omnivores) | 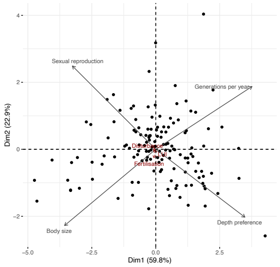  |

| | |
|---|---|
| Oribatid mites (omnivores) | 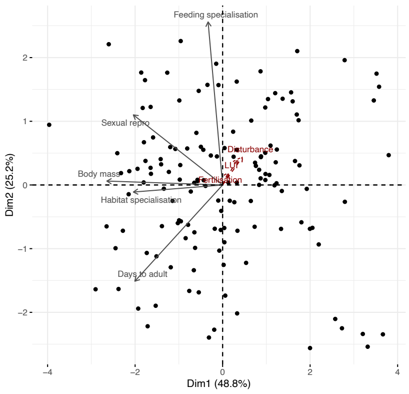 |

| | |
|---|---|
| Arthropods (secondary consumers, belowground) | 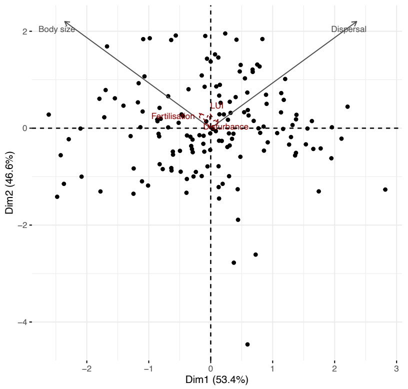  |

**Figure S 5.** Correlation between guild-level PCA axes or single traits and land use intensity. Three axes were retained when more than two traits were available; otherwise only two were retained. For protists, only one trait was available, and the correlation between the CWM of this trait and LUI is shown. The fast-slow axis is always the first PC axis, except for arthropod (primary and secondary consumers) above-ground, for which it was PC axis 2. The axes were transformed were needed (inversed sign) so that higher axis values indicate "faster" strategies. P-values were corrected for false detection rates (***: P < 0.001, **: P < 0.01, *: P< 0.05, n.s.: P > 0.05). **Community-level trait data (CWM) was not weighted by taxa abundance.**

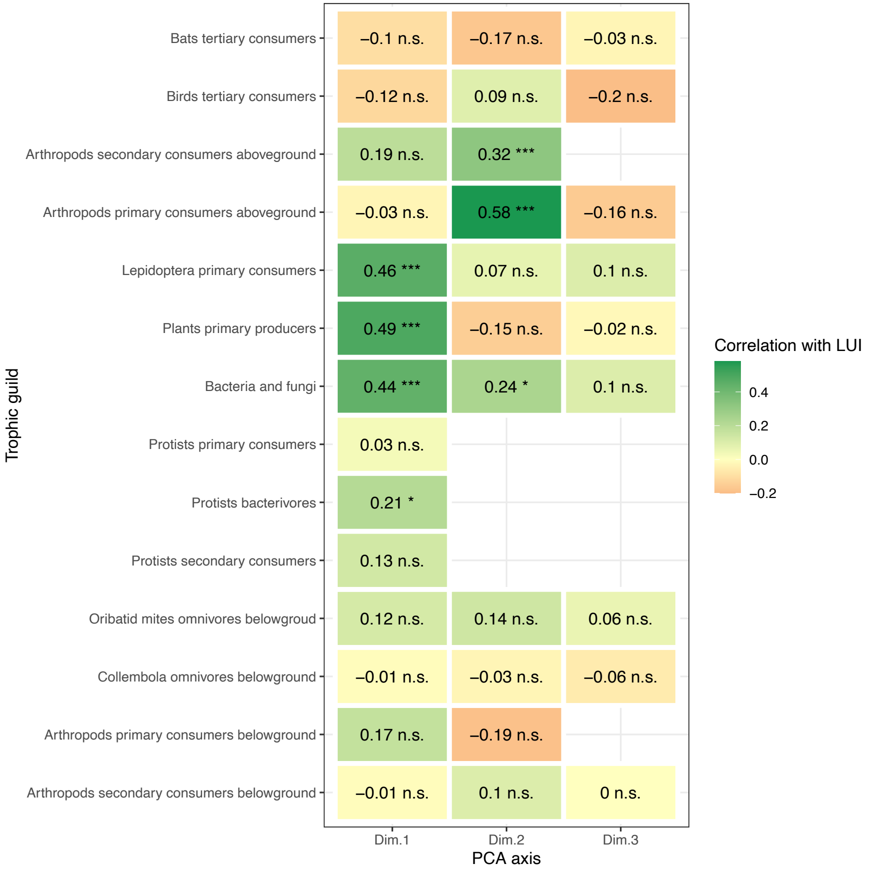

**Table S 9.** Trait-specific hypothesis testing: expected response of each trait to resource availability and disturbance (see **Table S 1** for detailed hypotheses); test of the hypothesised response**. Community-level trait data (CWM) was not weighted by taxa abundance.** Colours indicate trophic level (pale to dark colours) and position above (blue) or belowground (brown).

| **Guild** | **Trait** | **Expectation: fast or slow trait** | **Slope estimate (trait CWM ~LUI)** | **Response as expected?** |
|---|---|---|---|---|
| Vascular plants (primary producers) | Specific Leaf area | Fast | 0.36 (0.21 – 0.51)<br>P < 0.001 | Yes |
| | Seed mass | Slow | -0.21 (-0.35 -- 0.07)<br>P = 0.01 | Yes |
| | Leaf dry matter content | Slow | -0.31 (-0.45 -- -0.17)<br>P < 0.001 | Yes |
| | Leaf nitrogen | Fast | 0.42 (0.28 – 0.56)<br>P < 0.001 | Yes |
| | Leaf phosphorus | Fast | 0.51 (0.38 – 0.64)<br>P < 0.001 | Yes |
| | Root tissue density | Slow | -0.34 (-0.47 -- -0.2)<br>P < 0.001 | Yes |
| Lepidoptera | Flight period | Fast | 0.27 (0.16 – 0.38)<br>P < 0.001 | Yes |
| | Generations per year | Fast | 0.34 (0.18 – 0.48)<br>P < 0.001 | Yes |
| | Hibernation stage | Fast | 0.27 (0.14 – 0.39)<br>P < 0.001 | Yes |
| | Size (wing size) | Fast or slow | 0.15 (0.02 – 0.28)<br>P = 0.05 | Inconclusive |

| | | | | |
|---|---|---|---|---|
| | Generalism | Fast | 0.14 (0.0 – 0.29)<br>P = 0.10 | Inconclusive |
| Arthropods (primary consumers, aboveground) | Body size | Slow | -0.12 (-0.28 – 0.04)<br>P = 0.20 | Inconclusive |
| | Feeding generalism | Fast | 0.27 (0.13 – 0.42)<br>P < 0.001 | Yes |
| | Dispersal ability | Fast | 0.17 (0.03 – 0.32)<br>P < 0.04 | Yes |
| | Generations per year | Fast | 0.47 (0.35 – 0.58)<br>P < 0.001 | Yes |
| Arthropods (secondary consumers, aboveground) | Body size | Slow | -0.06 (-0.22 – 0.09)<br>P = 0.52 | Inconclusive |
| | Dispersal ability | Fast | 0.34 (0.19 – 0.49)<br>P < 0.001 | Yes |
| Birds (secondary consumers) | Body mass | Slow | 0.17 (0.02 – 0.32)<br>P = 0.04 | No |
| | Incubation time | Slow | 0.07 (-0.09 – 0.22)<br>P = 0.51 | Inconclusive |
| | Maximum number of offspring | Fast | -0.03 (-0.18 – 0.13)<br>P = 0.80 | Inconclusive |
| | Generation time | Slow | 0.13 (-0.02 – 0.28)<br>P = 0.15 | Inconclusive |
| Bats | Body mass | Slow | -0.15 (-0.28 – 0.03)<br>P = 0.03 | Yes |
| | Maximum longevity | Slow | 0.02 (-0.13 – 0.17)<br>P = 0.82 | Inconclusive |

| | | | | |
|---|---|---|---|---|
| | Number of offspring | Fast | -0.08 (-0.22 – 0.05)<br>P = 0.31 | Inconclusive |
| Protists (plant pathogens) | Relative abundance | Fast | Not applicable (based on relative abundances) | |
| Micro-organisms (bacteria and fungi) | bacterial cell volume | fast (a) or slow (b) | -0.27 (-0.42 -- -0.12)<br>P = 0.0015 | Yes (hypothesis B) |
| | Bacterial oligotroph:copiotroph ratio | slow | -0.21 ( -0.34 -- -0.08)<br>P = 0.0036 | Yes |
| | Bacterial genome size | slow | -0.16 (-0.28 -- -0.03)<br>P = 0.01 | Yes |
| | Fungi:bacteria ratio | slow | Not applicable (based on relative abundances) | |
| | Proportion of fungal pathotrophs among all fungi | fast | 0.37 (0.24 – 0.5)<br>P < 0.001 | Not applicable (based on relative abundances) |
| Protists (bacterivores) | Only one trait, cell size | slow | -0.19 (-0.33 – -0.05)<br>P = 0.02 | Yes |
| Protists (secondary consumers) | Only one trait, cell size | slow | -0.12 (-0.26 -- 0.02)<br>P = 0.14 | Inconclusive |
| Arthropods (primary consumers, belowground) | Body size | slow | -0.23 (-0.38 -- -0.07)<br>P = 0.01 | Yes |
| | Generalism | fast | -0.06 (-0.24 – 0.12)<br>P = 0.60 | Inconclusive |
| | Dispersal ability | fast | 0.14 (-0.02 – 0.31)<br>P = 0.15 | Inconclusive |

| | | | | |
|---|---|---|---|---|
| Collembola | Body size | slow | -0.02 (-0.13 – 0.14)<br>P = 0.82 | Inconclusive |
| | Depth preference | fast | -0.00 (-0.16 – 0.15)<br>P = 0.95 | Inconclusive |
| | Voltinism | fast | -0.06 (-0.21 – 0.1)<br>P = 0.57 | Inconclusive |
| | Reproduction type: sexual | slow | -0.0 (-0.15 – 0.16)<br>P = 0.96 | Inconclusive |
| Oribatid mites | Habitat specificity | slow | -0.06 (-0.22 – 0.11)<br>P = 0.54 | Inconclusive |
| | Reproduction type: sexual | slow | -0.04 (-0.2 – 0.11)<br>P = 0.72 | Inconclusive |
| | Days to maturity | slow | -0.10 (-0.25 – 0.05)<br>P = 0.30 | Inconclusive |
| | Body mass | slow | -0.15 (-0.29 – 0.0)<br>P = 0.09 | Inconclusive |
| Arthropods (secondary consumers, belowground) | Body size | slow | 0.06 (-0.07 – 0.19)<br>P = 0.51 | Inconclusive |
| | Dispersal ability | fast | 0.06 (-0.09 – 0.21)<br>P = 0.56 | Inconclusive |

**Figure S 6.** Synchronised slow-fast trait response of individual guilds is strongly related to land-use intensity. **Community-level trait data (CWM) was not weighted by taxa abundance**. The variables included in the PCA are the slow-fast axes of each guild. Land-use intensity, added as a supplementary variable, was strongly associated with axis 1. Belowground guilds are shown in brown, aboveground guilds in blue.

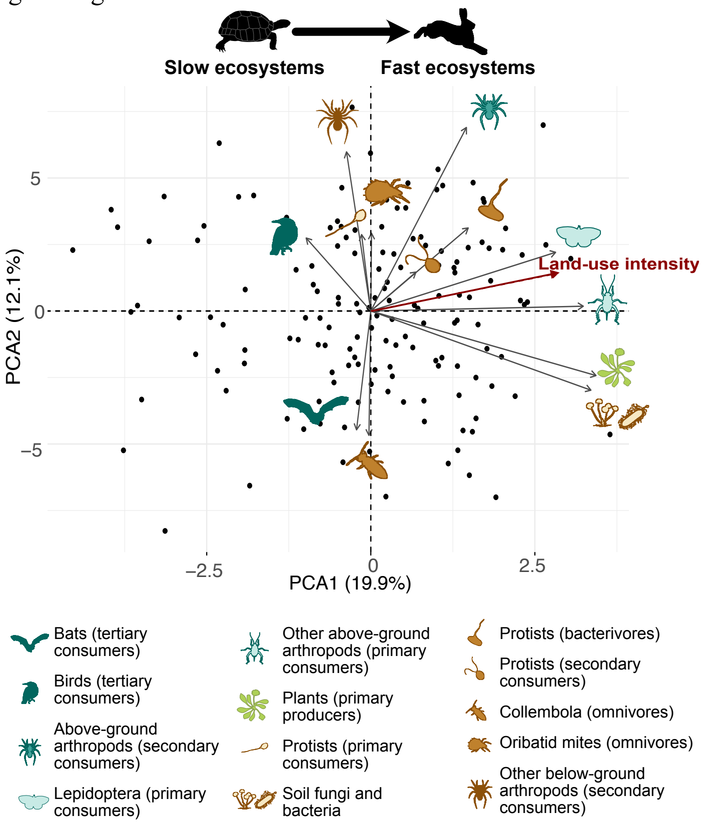

**Figure S 7.** Direct and trophically mediated effects of land-use intensity on the slow-fast axis of different trophic levels. **Community-level trait data (CWM) was not weighted by taxa abundance**. a. Full SEMs including all guilds. Two independent models were fitted for below- and aboveground guilds; plants being included in both. b. Average direct, indirect and total LUI effects on each trophic level (averaged from the full SEM). c. Decreasing direct, indirect and total LUI effects with trophic level. Each dot represents the estimated effect (+/- standard error) of an individual guild in the full SEM.

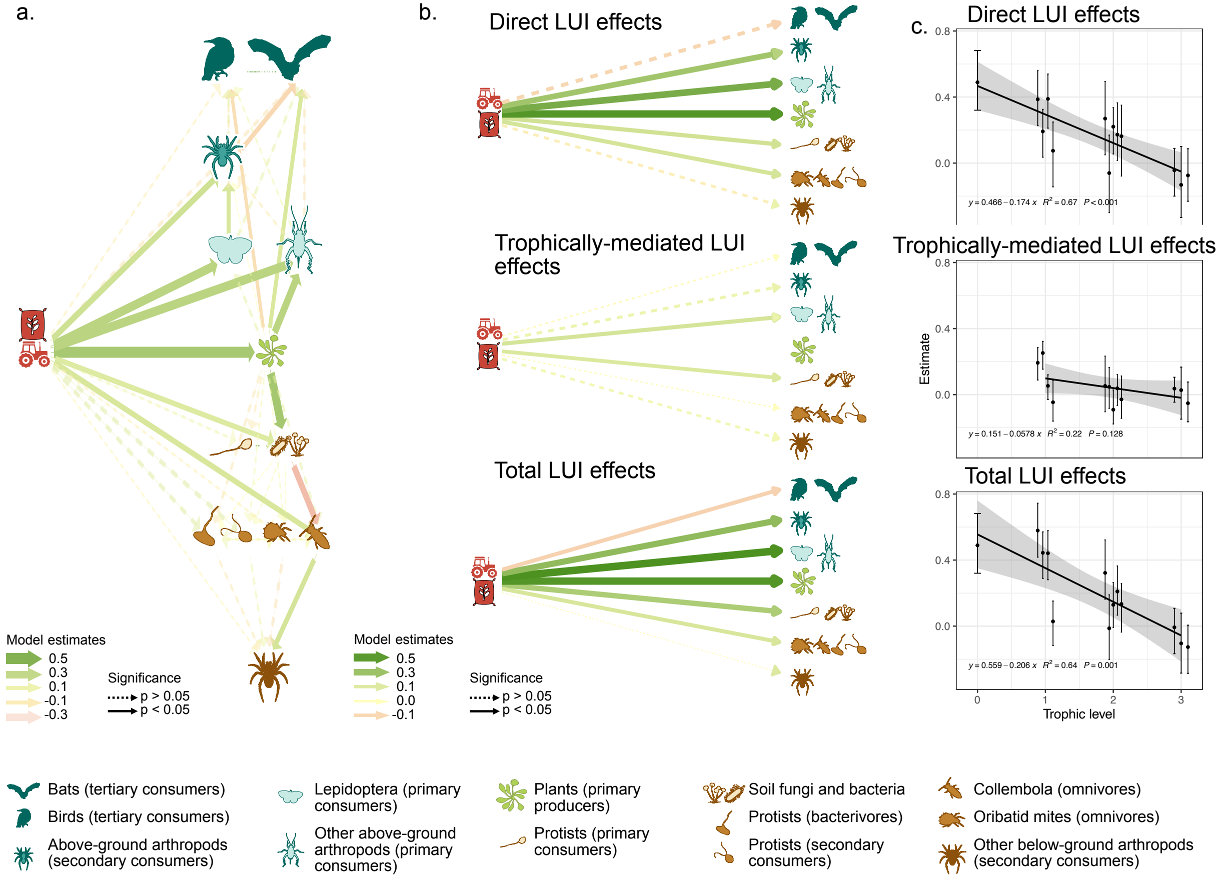

**Table S 10.** SEM path parameters for the belowground model (fitted with lavaan, boostrapped with 300 iterations). **Community-level trait data (CWM) was not weighted by taxa abundance.** Colours indicate trophic level (pale to dark colours).

| **Fit indices** | P-value = 0.37 | RMSEA = 0.0 | CFI = 1.00 | BIC = 2443 | |
|---|---|---|---|---|---|
| **Regression Slopes** | | | | | |
| **Left-hand side variable** | **Right-hand side variable** | **Estimate** | **Standard error** | **z** | **p** |
| Plant slow-fast axis | ~ land use intensity | 0.49 | 0.08 | 5.91 | .000 |
| Above-ground arthropods (primary consumers) slow-fast axis | ~ land use intensity | 0.39 | 0.08 | 4.78 | .000 |
| Above-ground arthropods (primary consumers) slow-fast axis | ~ Plant slow-fast axis | 0.39 | 0.08 | 4.95 | .000 |
| Lepidoptera (primary consumers) slow-fast axis | ~ land use intensity | 0.39 | 0.09 | 4.37 | .000 |
| Lepidoptera (primary consumers) slow-fast axis | ~ Plant slow-fast axis | 0.11 | 0.07 | 1.49 | .137 |
| Above-ground arthropods (secondary consumers) slow-fast axis | ~ land use intensity | 0.27 | 0.11 | 2.37 | .018 |
| Above-ground arthropods (secondary consumers) slow-fast axis | ~ Plant slow-fast axis | 0.00 | 0.09 | 0.01 | .990 |
| Above-ground arthropods (secondary consumers) slow-fast axis | ~ Lepidoptera (primary consumers) slow-fast axis | 0.23 | 0.11 | 1.98 | .048 |
| Above-ground arthropods (secondary consumers) slow-fast axis | ~ Above-ground arthropods (primary consumers) slow-fast axis | -0.08 | 0.10 | -0.79 | .429 |
| Birds (tertiary consumers) slow-fast axis | ~ land use intensity | -0.07 | 0.09 | -0.88 | .380 |
| Birds (tertiary consumers) slow-fast axis | ~ Plant slow-fast axis | -0.16 | 0.08 | -2.00 | .045 |
| Birds (tertiary consumers) slow-fast axis | ~ Lepidoptera (primary consumers) slow-fast axis | -0.10 | 0.10 | -1.03 | .303 |

| | | | | | |
|---|---|---|---|---|---|
| Birds (tertiary consumers) slow-fast axis | ~ Above-ground arthropods (primary consumers) slow-fast axis | 0.06 | 0.09 | 0.72 | .470 |
| Birds (tertiary consumers) slow-fast axis | ~ Above-ground arthropods (secondary consumers) slow-fast axis | 0.12 | 0.08 | 1.56 | .118 |
| Bats (tertiary consumers) slow-fast axis | ~ land use intensity | -0.13 | 0.11 | -1.25 | .213 |
| Bats (tertiary consumers) slow-fast axis | ~ Plant slow-fast axis | 0.21 | 0.09 | 2.43 | .015 |
| Bats (tertiary consumers) slow-fast axis | ~ Lepidoptera (primary consumers) slow-fast axis | 0.12 | 0.09 | 1.36 | .173 |
| Bats (tertiary consumers) slow-fast axis | ~ Above-ground arthropods (primary consumers) slow-fast axis | -0.12 | 0.10 | -1.20 | .230 |
| Bats (tertiary consumers) slow-fast axis | ~ Above-ground arthropods (secondary consumers) slow-fast axis | -0.19 | 0.08 | -2.25 | .024 |
| **Intercepts** | | | | | |
| Plant slow-fast axis | | 0.00 | 0.07 | 0.00 | 1.000 |
| Above-ground arthropods (primary consumers) slow-fast axis | | -0.00 | 0.07 | -0.00 | 1.000 |
| Lepidoptera (primary consumers) slow-fast axis | | 0.01 | 0.08 | 0.11 | .915 |
| Above-ground arthropods (secondary consumers) slow-fast axis | | -0.00 | 0.07 | -0.02 | .980 |
| Birds (tertiary consumers) slow-fast axis | | -0.01 | 0.09 | -0.10 | .918 |
| Bats (tertiary consumers) slow-fast axis | | -0.01 | 0.09 | -0.11 | .911 |
| LUI | | 0 (fixed parameter) | | | |
| **Residual Variances** | | | | | |
| Plant slow-fast axis | | 0.76 | 0.10 | 7.28 | .000 |
| Above-ground arthropods (primary consumers) slow-fast axis | | 0.54 | 0.06 | 8.64 | .000 |

| | | | | | |
|---|---|---|---|---|---|
| Lepidoptera (primary consumers) slow-fast axis | | 0.77 | 0.14 | 5.47 | .000 |
| Above-ground arthropods (secondary consumers) slow-fast axis | | 0.85 | 0.09 | 9.71 | .000 |
| Birds (tertiary consumers) slow-fast axis | | 0.94 | 0.16 | 5.81 | .000 |
| Bats (tertiary consumers) slow-fast axis | | 0.92 | 0.13 | 7.22 | .000 |
| LUI | | 1 (fixed parameter) | | | |
| **Residual Variances** | | | | | |
| Birds (tertiary consumers) slow-fast axis | Bats (tertiary consumers) slow-fast axis | -0.11 | 0.08 | -1.35 | .176 |

**Table S 11.** SEM path parameters for the belowground model (fitted with lavaan, boostrapped with 300 iterations). **Community-level trait data (CWM) was not weighted by taxa abundance**. Note the weaker model fit than in the main results. Colours indicate trophic level (pale to dark colours).

| **Fit indices** | P-value = 0.48 | RMSEA = 0.02 | CFI =1 | BIC = 3371 | |
|---|---|---|---|---|---|
| **Regression Slopes** | | | | | |
| **Left-hand side variable** | **Right-hand side variable** | **Estimate** | **Standard error** | **z** | **p** |
| Plant slow-fast axis | ~ land use intensity | 0.49 | 0.07 | 6.75 | .000 |
| Protists (pathotrophs. i.e.e primary consumers) slow-fast axis | ~ land use intensity | 0.07 | 0.11 | 0.69 | .491 |
| Protists (pathotrophs. i.e.e primary consumers) slow-fast axis | ~ Plant slow-fast axis | -0.09 | 0.12 | -0.79 | .432 |
| Bacteria and fungi slow-fast axis | ~ land use intensity | 0.19 | 0.06 | 3.07 | .002 |
| Bacteria and fungi slow-fast axis | ~ Plant slow-fast axis | 0.51 | 0.07 | 7.47 | .000 |
| Protists (bacterivores) slow-fast axis | ~ land use intensity | 0.17 | 0.09 | 1.92 | .055 |
| Protists (bacterivores) slow-fast axis | ~ Plant slow-fast axis | 0.05 | 0.10 | 0.45 | .650 |
| Protists (bacterivores) slow-fast axis | ~ Bacteria and fungi slow-fast axis | 0.03 | 0.10 | 0.33 | .742 |
| Protists (secondary consumers) slow-fast axis | ~ land use intensity | 0.16 | 0.11 | 1.52 | .129 |
| Protists (secondary consumers) slow-fast axis | ~ Plant slow-fast axis | -0.11 | 0.10 | -1.07 | .285 |
| Protists (secondary consumers) slow-fast axis | ~ Bacteria and fungi slow-fast axis | 0.00 | 0.12 | 0.02 | .981 |
| Protists (secondary consumers) slow-fast axis | ~ Protists (bacterivores) slow-fast axis | 0.10 | 0.08 | 1.26 | .209 |
| Oribatid mites (omnivores) slow-fast axis | ~ land use intensity | 0.22 | 0.07 | 3.10 | .002 |

| | | | | | |
|---|---|---|---|---|---|
| Oribatid mites (omnivores) slow-fast axis | ~ Plant slow-fast axis | 0.11 | 0.10 | 1.09 | .275 |
| Oribatid mites (omnivores) slow-fast axis | ~ Bacteria and fungi slow-fast axis | -0.33 | 0.11 | -2.87 | .004 |
| Oribatid mites (omnivores) slow-fast axis | ~ Protists (bacterivores) slow-fast axis | 0.03 | 0.08 | 0.33 | .742 |
| Oribatid mites (omnivores) slow-fast axis | ~ Protists (secondary consumers) slow-fast axis | -0.04 | 0.09 | -0.41 | .684 |
| Collembola (omnivores) slow-fast axis | ~ land use intensity | -0.06 | 0.12 | -0.50 | .617 |
| Collembola (omnivores) slow-fast axis | ~ Plant slow-fast axis | 0.04 | 0.10 | 0.42 | .672 |
| Collembola (omnivores) slow-fast axis | ~ Bacteria and fungi slow-fast axis | 0.02 | 0.11 | 0.20 | .840 |
| Collembola (omnivores) slow-fast axis | ~ Protists (bacterivores) slow-fast axis | 0.05 | 0.09 | 0.63 | .528 |
| Collembola (omnivores) slow-fast axis | ~ Protists (secondary consumers) slow-fast axis | 0.04 | 0.07 | 0.65 | .514 |
| Arthropods (secondary consumers) slow-fast axis | ~ land use intensity | -0.04 | 0.08 | -0.56 | .575 |
| Arthropods (secondary consumers) slow-fast axis | ~ Oribatid mites slow-fast axis | 0.24 | 0.08 | 3.00 | .003 |
| Arthropods (secondary consumers) slow-fast axis | ~ Collembola slow-fast axis | -0.14 | 0.08 | -1.63 | .103 |
| Arthropods (secondary consumers) slow-fast axis | ~ Protists (bacterivores) slow-fast axis | 0.08 | 0.08 | 0.97 | .334 |
| Arthropods (secondary consumers) slow-fast axis | ~ Protists (secondary consumers) slow-fast axis | -0.09 | 0.07 | -1.24 | .216 |
| **Intercepts** | | | | | |
| Plant slow-fast axis | | 0.00 | 0.07 | 0.00 | 1.000 |
| Protists (primary consumers) slow-fast axis | | 0.00 | 0.09 | 0.00 | 1.000 |
| Bacteria and fungi fast-slow axis | | 0.00 | 0.06 | 0.00 | 1.000 |

| | | | | | |
|---|---|---|---|---|---|
| Protists (bacterivores) slow-fast axis | | -0.00 | 0.07 | -0.00 | 1.000 |
| Protists (secondary consumers) slow-fast axis | | 0.00 | 0.09 | 0.00 | 1.000 |
| Oribatid mites (omnivores) slow-fast axis | | -0.00 | 0.08 | -0.03 | .978 |
| Collembola (omnivores) slow-fast axis | | 0.00 | 0.09 | 0.00 | .996 |
| Above-ground arthropods (secondary consumers) slow-fast axis | | 0.00 | 0.07 | 0.01 | .994 |
| LUI | | 0 (fixed parameter) | | | |
| **Residual Variances** | | | | | |
| Plant slow-fast axis | | 0.76 | 0.11 | 7.12 | .000 |
| Protists (primary consumers) slow-fast axis | | 0.99 | 0.39 | 2.55 | .011 |
| Bacteria and fungi fast-slow axis | | 0.60 | 0.09 | 7.00 | .000 |
| Protists (bacterivores) slow-fast axis | | 0.95 | 0.12 | 7.94 | .000 |
| Protists (secondary consumers) slow-fast axis | | 0.96 | 0.17 | 5.50 | .000 |
| Oribatid mites (omnivores) slow-fast axis | | 0.91 | 0.10 | 9.44 | .000 |
| Collembola (omnivores) slow-fast axis | | 0.99 | 0.13 | 7.54 | .000 |
| Above-ground arthropods (secondary consumers) slow-fast axis | | 0.91 | 0.10 | 8.96 | .000 |
| LUI | | 1 (fixed parameter) | | | |
| **Residual Coariances** | | | | | |
| Protists (primary consumers) slow-fast axis | Bacteria and fungi fast-slow axis | -0.09 | 0.05 | -1.76 | .079 |

| | | |
|---|---|---|
| Protists (primary consumers) slow-fast axis | | 0 (fixed parameter) |

**Figure S 8.** Direct and indirect links between land- use intensity, functional traits slow-fast axis and ecosystem function slow-fast axis. **In contrast to Fig. S4, community-level trait data (CWM) was not weighted by taxa abundance**

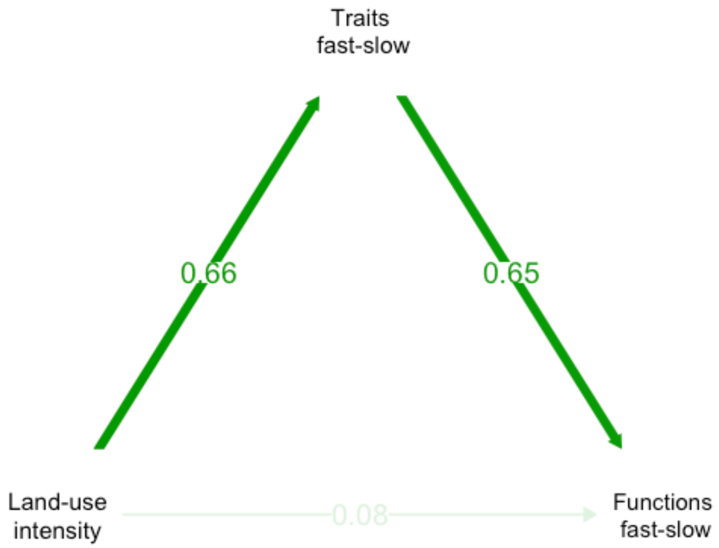

**Table S 12.** Comparison of the effect of multiple drivers on the ecosystem functions slow-fast axis, obtained from linear models with the function slow-fast axis as a response and the indicated variables as explanatory variables. **Community-level trait data (CWM) was not weighted by taxa abundance.** P-values were corrected for multiple testing within these different model (i.e. correction for false discovery rates, R function p.adjust, n = 5).

| **Model** | **Slope estimate (+/- standard error)** | **P-value** | **Adj $R^2$** |
|---|---|---|---|
| Functions slow-fast ~ plants slow-fast | 0.36 (0.05) | P < 0.001 | 32% |
| Functions slow-fast ~ microbial slow-fast | 0.41 (0.06) | P < 0.001 | 51% |
| Functions slow-fast ~ whole community slow-fast | 0.58 (0.05) | P < 0.001 | 46% |
| Functions slow-fast ~ taxonomic multidiversity | -0.42 (0.09) | P < 0.001 | 11% |
| Functions slow-fast ~ land-use intensity | 0.62 (0.09) | P < 0.001 | 26% |

**Table S 13.** Identification of guild-level slow-fast axes. **Community-level traits (CWM) were not corrected for environmental covariates.** Different colors show the different regions of the Exploratories.

| Guild | Community-level PCA | Comment |
|---|---|---|
| Vascular plants (primary producers) | 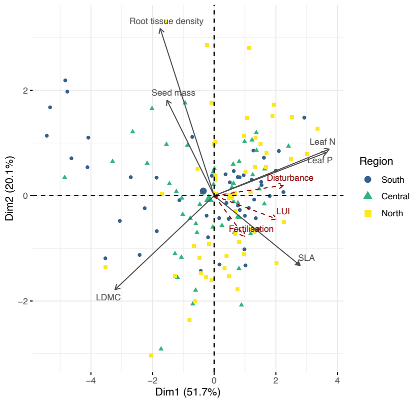 | |
| Lepidoptera (primary consumers) | 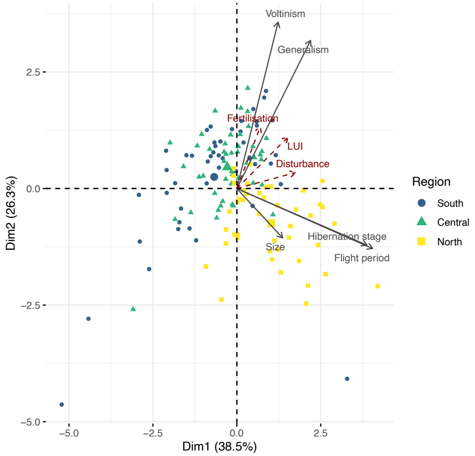 | Partial: no response from body size. Body size was measured as wing length, which is an indicator of overall body size (which is expected to be a "slow" trait; smaller size helps to survive disturbance) but also of dispersal ability ("fast" trait, as larger wings promote recolonisation after disturbance). Both effects might cancel each other leading to no response of size. |

| | | |
|---|---|---|
| Other arthropods (primary consumers, aboveground) | 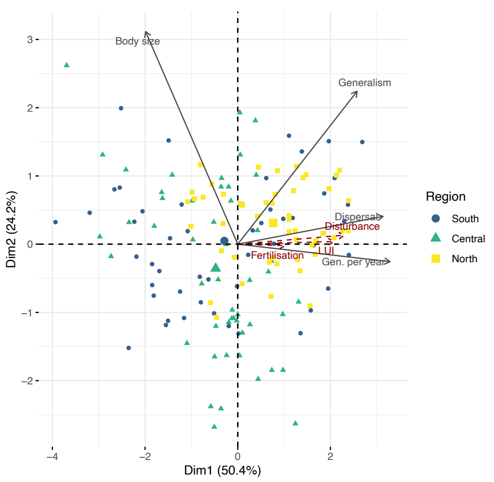 | |
| Arthropods (secondary consumers, aboveground) | 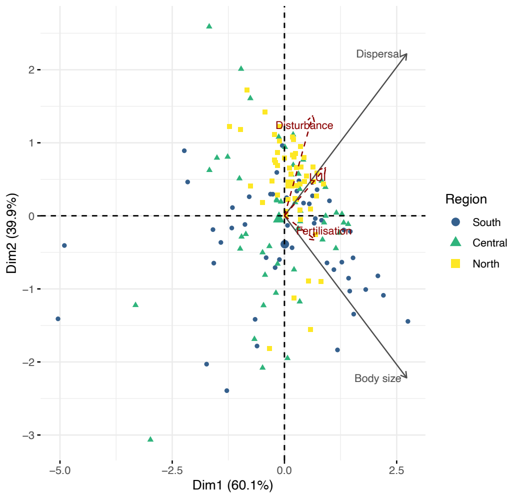 | Dispersal and body size are slightly confounded because larger body size increases dispersal abilities. However, these two traits are opposed on the second axis of the PCA, which we use as our slow-fast index. |

<table>
<tr><td>Birds (tertiary consumers)</td><td>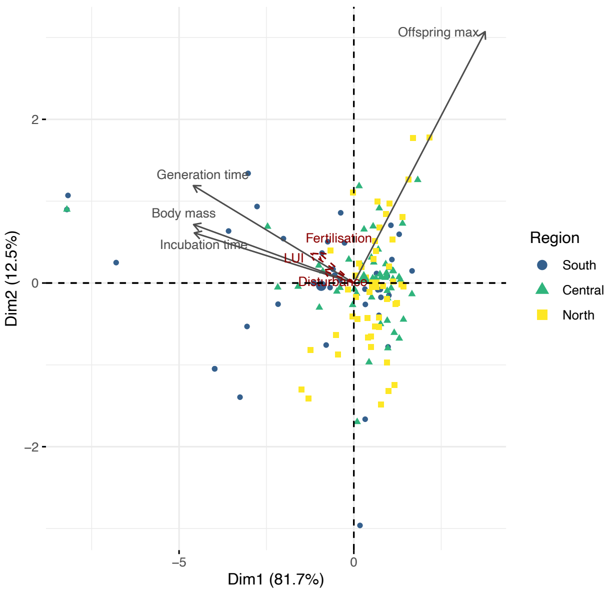
</td><td></td></tr>
<tr><td>Bats (tertiary consumers)</td><td>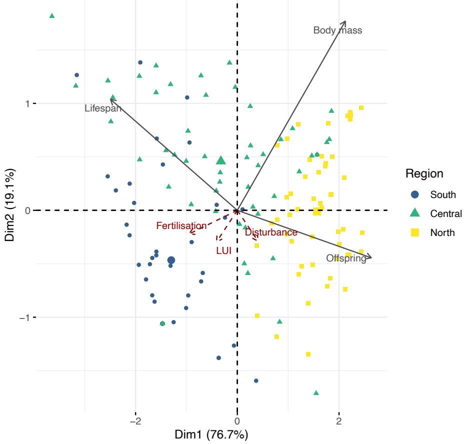
</td><td>Partial<br><br>Body mass is usually considered as a ‘slow’ trait, and is expected to be positively correlated to lifespan and negatively to the number of offspring (trade-off between survival and reproduction). Hibernation saves resources and leads, in hibernating bats (most of the species observed in our study), to a correlation between number of offspring and body mass (Wilkinson 2002), leading to the results observed here.</td></tr>
</table>

| | |
|---|---|
| Protists (plant pathogens, ie primary consumers) | Only one trait |
| Bacteria and fungi | 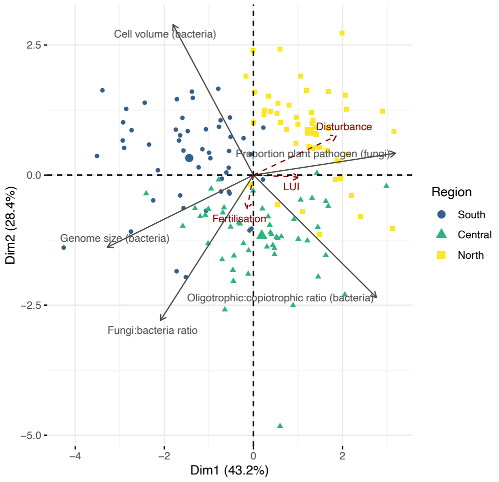 |
| Protists (bacterivores) | Only one trait |
| Protists (secondary consumers) | Only one trait |

| | |
|---|---|
| Arthropods (primary consumers, belowground) | 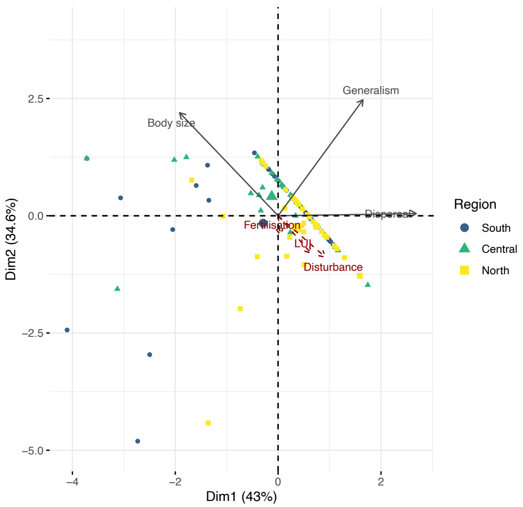 |
| Collembola (omnivores) | 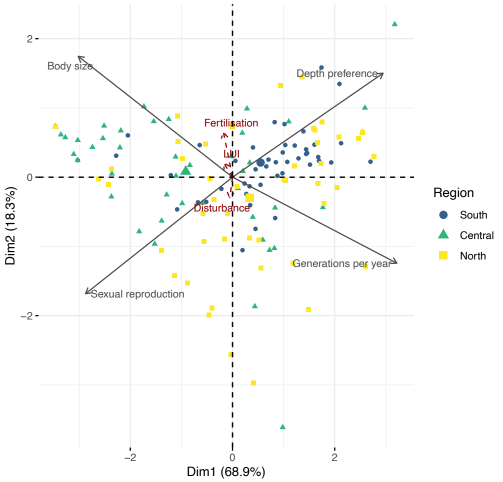 |

| | |
|---|---|
| Oribatid mites (omnivores) | 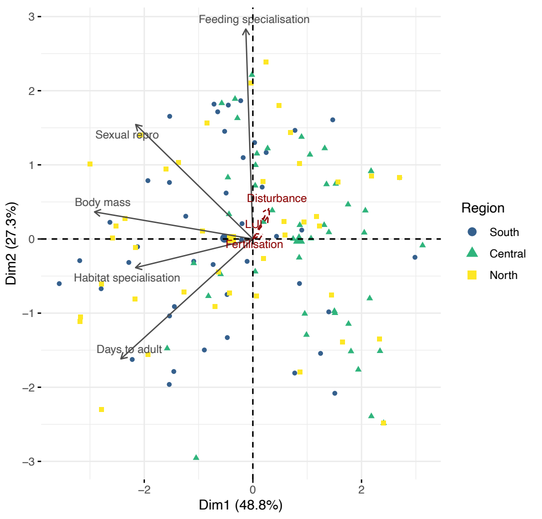 |
| Arthropods (secondary consumers, belowground) | 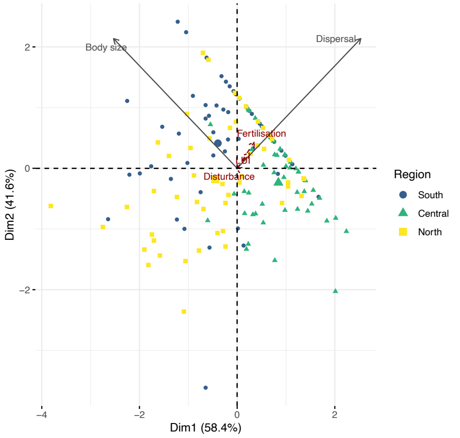 |

**Table S 14.** Trait-specific hypothesis testing: expected response of each trait to resource availability and disturbance (see **Table S 1** for detailed hypotheses). **Community-level traits (CWM) were not corrected for environmental covariates.** Colours indicate trophic level (pale to dark colours) and position above (blue) or belowground (brown).

| **Guild** | **Trait** | **Expectation: fast or slow trait** | **Slope estimate (trait CWM ~LUI)** | **Response as expected?** |
|---|---|---|---|---|
| Vascular plants (primary producers) | Specific Leaf area | Fast | 0.31 (0.16– 0.46)<br>P < 0.001 | Yes |
| | Seed mass | Slow | -0.31 (-0.46 – -0.15)<br>P < 0.001 | Yes |
| | Leaf dry matter content | Slow | -0.29 (-0.45– -0.14)<br>P < 0.001 | Yes |
| | Leaf nitrogen | Fast | 0.46 (0.31 – 0.60)<br>P < 0.001 | Yes |
| | Leaf phosphorus | Fast | 0.50 (0.36 – 0.64)<br>P < 0.001 | Yes |
| | Root tissue density | Slow | -0.28 (-0.44 – -0.13)<br>P = 0.001 | Yes |
| Lepidoptera | Flight period | Fast | 0.26 (0.1 – 0.42)<br>P = 0.004 | Yes |
| | Generations per year | Fast | 0.25 (0.1 – 0.41)<br>P = 0.004 | Yes |
| | Hibernation stage | Fast | 0.25 (0.09 – 0.41)<br>P = 0.006 | Yes |
| | Size (wing size) | Fast or slow | -0.12 (-0.28 – 0.04)<br>P = 0.21 | Inconclusive |

| | | | | |
|---|---|---|---|---|
| | Generalism | Fast | 0.31 (0.16 – 0.47)<br>P = 0.001 | Yes |
| Arthropods (primary consumers, aboveground) | Body size | Slow | -0.26 (-0.42 – -0.1)<br>P = 0.003 | Yes |
| | Feeding generalism | Fast | 0.39 (0.24 – 0.54)<br>P < 0.001 | Yes |
| | Dispersal ability | Fast | 0.28 (0.13 – 0.44)<br>P = 0.001 | Yes |
| | Generations per year | Fast | 0.54 (0.4 – 0.68)<br>P = 0.001 | Yes |
| Arthropods (secondary consumers, aboveground) | Body size | Slow | 0.08 (-0.08 – 0.25)<br>P = 0.40 | Inconclusive |
| | Dispersal ability | Fast | 0.32 (0.17 – 0.48)<br>P < 0.001 | Yes |
| Birds (secondary consumers) | Body mass | Slow | 0.31 (0.15 – 0.47)<br>P < 0.001 | No |
| | Incubation time | Slow | 0.18 (0.02 – 0.34)<br>P = 0.05 | Inconclusive |
| | Maximum number of offspring | Fast | -0.15 (-0.31 – 0.01)<br>P = 0.11 | No |
| | Generation time | Slow | 0.27 (0.11 – 0.42)<br>P = 0.003 | No |
| Bats | Body mass | Slow | -0.17 (-0.33 – -0.01)<br>P = 0.062 | Inconclusive |
| | Maximum longevity | Slow | 0.06 (-0.1 – 0.23)<br>P = 0.54 | Inconclusive |

| | | | | |
|---|---|---|---|---|
| | Number of offspring | Fast | -0.16 (-0.32 – 0.01)<br>P = 0.098 | Inconclusive |
| Protists (plant pathogens) | Relative abundance | Fast | 0.46 (0.32 – 0.61)<br>P < 0.001 | Yes |
| Micro-organisms (bacteria and fungi) | bacterial cell volume | fast (a) or slow (b) | -0.31 (-0.46 – -0.15)<br>P < 0.001 | Yes (hypothesis B) |
| | Bacterial oligotroph:copiotroph ratio | slow | -0.01 (-0.15 – 0.17)<br>P = 0.95 | Inconclusive |
| | Bacterial genome size | slow | -0.04 (-0.21 – 0.12)<br>P = 0.71 | Inconclusive |
| | Fungi:bacteria ratio | slow | -0.24 (-0.4 – -0.08)<br>P = 0.006 | Yes |
| | Proportion of fungal pathotrophs among all fungi | fast | 0.31 (0.15 – 0.46)<br>P < 0.001 | Yes |
| Protists (bacterivores) | Only one trait, cell size | slow | -0.37 (-0.52 – -0.21)<br>P < 0.001 | Yes |
| Protists (secondary consumers) | Only one trait, cell size | slow | -0.30 (-0.46 – -0.15)<br>P < 0.001 | Yes |
| Arthropods (primary consumers, belowground) | Body size | slow | -0.25 (-0.41 – -0.09)<br>P = 0.006 | Yes |
| | Generalism | fast | -0.09 (-0.28 – 0.1)<br>P = 0.44 | Inconclusive |
| | Dispersal ability | fast | 0.15 (-0.01 – 0.32)<br>P = 0.11 | Inconclusive |

| | | | | |
|---|---|---|---|---|
| Collembola | Body size | slow | 0.01 (-0.13 – 0.21)<br>P = 0.73 | Inconclusive |
| | Depth preference | fast | -0.01 (-0.18 – 0.16)<br>P = 0.95 | Inconclusive |
| | Voltinism | fast | -0.09 (-0.26 – 0.08)<br>P = 0.40 | Inconclusive |
| | Reproduction type: sexual | slow | 0.01 (-0.16 – 0.18)<br>P = 0.97 | Inconclusive |
| Oribatid mites | Habitat specificity | slow | 0.03 (-0.2 – 0.14)<br>P = 0.78 | Inconclusive |
| | Reproduction type: sexual | slow | 0 (-0.17 – 0.17)<br>P = 0.97 | Inconclusive |
| | Days to maturity | slow | -0.06 (-0.23 – 0.11)<br>P = 0.59 | Inconclusive |
| | Body mass | slow | -0.16 (-0.33 – 0)<br>P = 0.09 | Inconclusive |
| Arthropods (secondary consumers, belowground) | Body size | slow | -0.01 (-0.17 – 0.15)<br>P = 0.95 | Inconclusive |
| | Dispersal ability | fast | 0.11 (-0.05 – 0.27)<br>P = 0.26 | Inconclusive |

**Figure S 9.** Correlation between guild-level PCA axes or single traits and land use intensity. Three axes were retained when more than two traits were available; otherwise only two were retained. For protists, only one trait was available, and the correlation between the CWM of this trait and LUI is shown. The fast-slow axis is always the first PC axis, except for arthropod (secondary consumers) above-ground, for which it was PC axis 2. The axes were transformed were needed (inversed sign) so that higher axis values indicate "faster" strategies. P-values were corrected for false detection rates (***: P < 0.001, **: P < 0.01, *: P< 0.05, n.s.: P > 0.05). **Community-level traits (CWM) were not corrected for environmental covariates**.

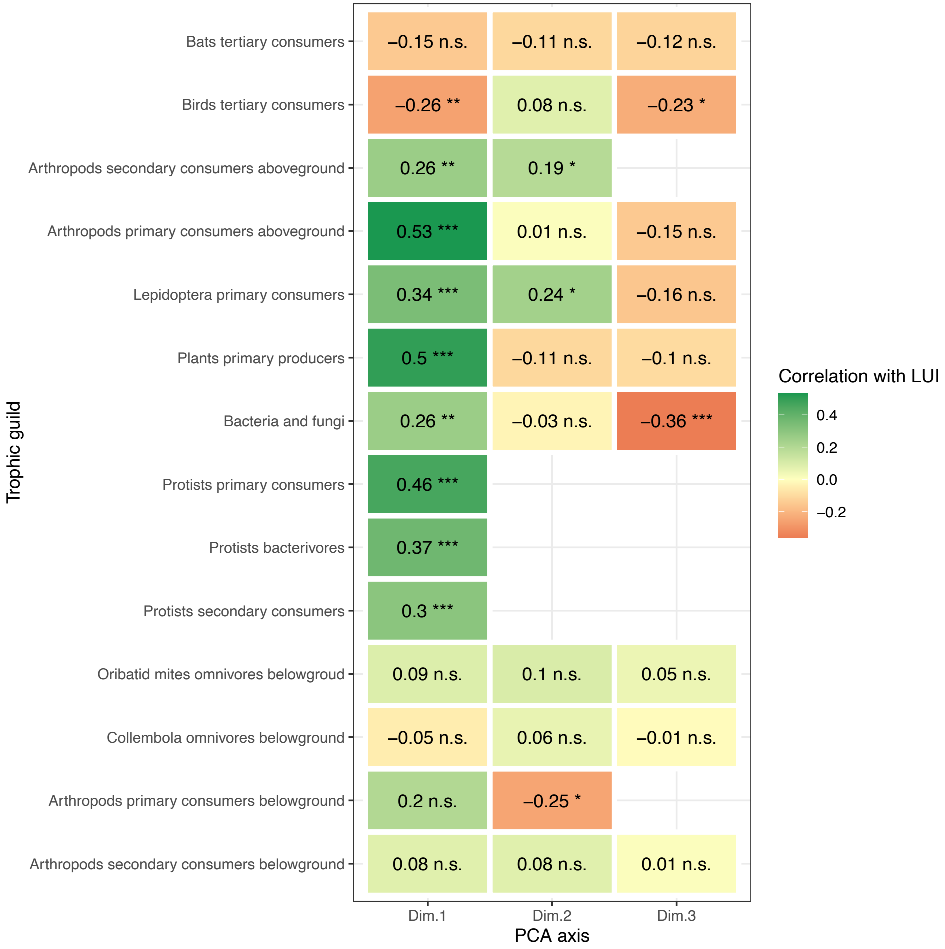

**Figure S 10.** Synchronised slow-fast trait response of individual guilds is strongly related to land-use intensity. **Community-level traits (CWM) were not corrected for environmental covariates**. The variables included in the PCA are the slow-fast axes of each guild. Land-use intensity, added as a supplementary variable, was strongly associated with axis 1. Belowground guilds are shown in brown, aboveground guilds in blue.

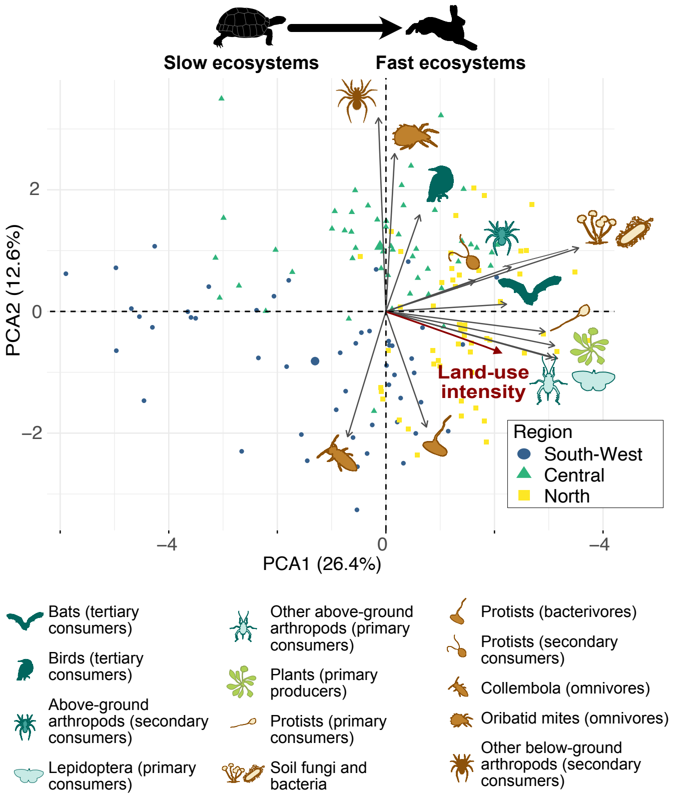

**Figure S 11.** Direct and trophically mediated effects of land-use intensity on the slow-fast axis of different trophic levels**. Community-level traits (CWM) were not corrected for environmental covariates.** a. Full SEMs including all guilds. Two independent models were fitted for below- and aboveground guilds; plants being included in both. b. Average direct, indirect and total LUI effects on each trophic level (averaged from the full SEM). c. Decreasing direct, indirect and total LUI effects with trophic level. Each dot represents the estimated effect (+/- standard error) of an individual guild in the full SEM.

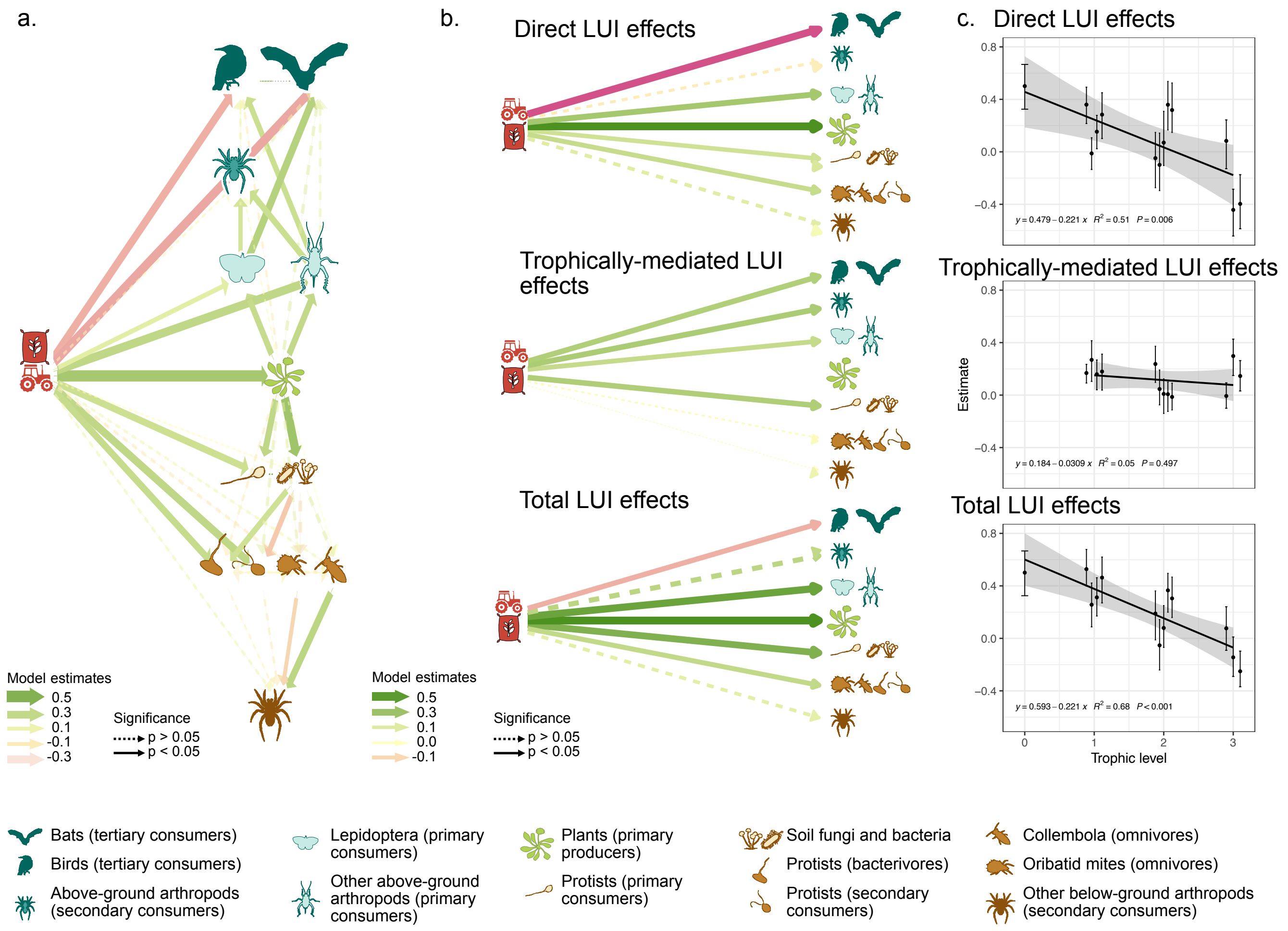

**Table S 15.** SEM path parameters for the belowground model (fitted with lavaan, boostrapped with 300 iterations). **Community-level traits (CWM) were not corrected for environmental covariates.** Note that the model fit is low, likely due to the influence of environmental variables that were not accounted for. Colours indicate trophic level (pale to dark colours).

| **Fit indices** | P-value = 0.02 | RMSEA = 0.18 | CFI = 0.98 | BIC = 2404.58 |
|---|---|---|---|---|

**Regression Slopes**

| **Left-hand side variable** | **Right-hand side variable** | **Estimate** | **Standard error** | **z** | **p** |
|---|---|---|---|---|---|
| Plant slow-fast axis | ~ land use intensity | 0.50 | 0.07 | 6.71 | .000 |
| Above-ground arthropods (primary consumers) slow-fast axis | ~ land use intensity | 0.36 | 0.07 | 5.18 | .000 |
| Above-ground arthropods (primary consumers) slow-fast axis | ~ Plant slow-fast axis | 0.34 | 0.06 | 5.79 | .000 |
| Lepidoptera (primary consumers) slow-fast axis | ~ land use intensity | 0.15 | 0.07 | 2.27 | .023 |
| Lepidoptera (primary consumers) slow-fast axis | ~ Plant slow-fast axis | 0.32 | 0.09 | 3.37 | .001 |
| Above-ground arthropods (secondary consumers) slow-fast axis | ~ land use intensity | -0.05 | 0.11 | -0.42 | .674 |
| Above-ground arthropods (secondary consumers) slow-fast axis | ~ Plant slow-fast axis | 0.02 | 0.08 | 0.29 | .768 |
| Above-ground arthropods (secondary consumers) slow-fast axis | ~ Lepidoptera (primary consumers) slow-fast axis | 0.24 | 0.08 | 3.00 | .003 |
| Above-ground arthropods (secondary consumers) slow-fast axis | ~ Above-ground arthropods (primary consumers) slow-fast axis | 0.29 | 0.10 | 2.78 | .005 |
| Birds (tertiary consumers) slow-fast axis | ~ land use intensity | -0.40 | 0.10 | -3.91 | .000 |
| Birds (tertiary consumers) slow-fast axis | ~ Plant slow-fast axis | -0.09 | 0.08 | -1.17 | .242 |

| | | | | | |
|---|---|---|---|---|---|
| Birds (tertiary consumers) slow-fast axis | ~ Lepidoptera (primary consumers) slow-fast axis | 0.08 | 0.09 | 0.91 | .362 |
| Birds (tertiary consumers) slow-fast axis | ~ Above-ground arthropods (primary consumers) slow-fast axis | 0.29 | 0.14 | 2.10 | .035 |
| Birds (tertiary consumers) slow-fast axis | ~ Above-ground arthropods (secondary consumers) slow-fast axis | 0.06 | 0.09 | 0.69 | .489 |
| Bats (tertiary consumers) slow-fast axis | ~ land use intensity | -0.44 | 0.09 | -4.72 | .000 |
| Bats (tertiary consumers) slow-fast axis | ~ Plant slow-fast axis | 0.16 | 0.09 | 1.92 | .054 |
| Bats (tertiary consumers) slow-fast axis | ~ Lepidoptera (primary consumers) slow-fast axis | 0.41 | 0.08 | 5.33 | .000 |
| Bats (tertiary consumers) slow-fast axis | ~ Above-ground arthropods (primary consumers) slow-fast axis | 0.12 | 0.09 | 1.38 | .169 |
| Bats (tertiary consumers) slow-fast axis | ~ Above-ground arthropods (secondary consumers) slow-fast axis | 0.11 | 0.08 | 1.45 | .148 |
| **Intercepts** | | | | | |
| Plant slow-fast axis | | 0.00 | 0.08 | 0.00 | 1.000 |
| Above-ground arthropods (primary consumers) slow-fast axis | | -0.00 | 0.07 | -0.00 | 1.000 |
| Lepidoptera (primary consumers) slow-fast axis | | 0.01 | 0.08 | 0.10 | .922 |
| Above-ground arthropods (secondary consumers) slow-fast axis | | -0.00 | 0.08 | -0.02 | .981 |
| Birds (tertiary consumers) slow-fast axis | | -0.01 | 0.08 | -0.06 | .950 |
| Bats (tertiary consumers) slow-fast axis | | -0.01 | 0.07 | -0.18 | .858 |
| LUI | | 0 (fixed parameter) | | | |
| **Residual Variances** | | | | | |
| Plant slow-fast axis | | 0.74 | 0.10 | 7.81 | .000 |

| | | | | | |
|---|---|---|---|---|---|
| Above-ground arthropods (primary consumers) slow-fast axis | | 0.63 | 0.06 | 10.39 | .000 |
| Lepidoptera (primary consumers) slow-fast axis | | 0.80 | 0.12 | 6.72 | .000 |
| Above-ground arthropods (secondary consumers) slow-fast axis | | 0.82 | 0.11 | 7.23 | .000 |
| Birds (tertiary consumers) slow-fast axis | | 0.85 | 0.23 | 3.64 | .000 |
| Bats (tertiary consumers) slow-fast axis | | 0.69 | 0.08 | 8.40 | .000 |
| LUI | | 1 (fixed parameter) | | | |
| **Residual Variances** | | | | | |
| Birds (tertiary consumers) slow-fast axis | Bats (tertiary consumers) slow-fast axis | 0.09 | 0.06 | 1.56 | .120 |

**Table S 16.** SEM path parameters for the belowground model (fitted with lavaan, boostrapped with 300 iterations). **Community-level traits (CWM) were not corrected for environmental covariates.** Colours indicate trophic level (pale to dark colours).

| **Fit indices** | P-value = 0.4 | RMSEA = 0.02 | CFI = 1.00 | BIC = 3270.13 |
|---|---|---|---|---|

**Regression Slopes**

| **Left-hand side variable** | **Right-hand side variable** | **Estimate** | **Standard error** | **z** | **p** |
|---|---|---|---|---|---|
| Plant slow-fast axis | ~ land use intensity | 0.50 | 0.08 | 6.10 | .000 |
| Protists (pathotrophs. i.e.e primary consumers) slow-fast axis | ~ land use intensity | 0.28 | 0.09 | 3.24 | .001 |
| Protists (pathotrophs. i.e.e primary consumers) slow-fast axis | ~ Plant slow-fast axis | 0.36 | 0.09 | 3.78 | .000 |
| Bacteria and fungi slow-fast axis | ~ land use intensity | -0.01 | 0.06 | -0.19 | .850 |
| Bacteria and fungi slow-fast axis | ~ Plant slow-fast axis | 0.54 | 0.11 | 5.06 | .000 |
| Protists (bacterivores) slow-fast axis | ~ land use intensity | 0.36 | 0.09 | 3.95 | .000 |
| Protists (bacterivores) slow-fast axis | ~ Plant slow-fast axis | 0.15 | 0.12 | 1.29 | .195 |
| Protists (bacterivores) slow-fast axis | ~ Bacteria and fungi slow-fast axis | -0.27 | 0.08 | -3.28 | .001 |
| Protists (secondary consumers) slow-fast axis | ~ land use intensity | 0.32 | 0.10 | 3.35 | .001 |
| Protists (secondary consumers) slow-fast axis | ~ Plant slow-fast axis | -0.09 | 0.10 | -0.92 | .358 |
| Protists (secondary consumers) slow-fast axis | ~ Bacteria and fungi slow-fast axis | 0.27 | 0.10 | 2.85 | .004 |
| Protists (secondary consumers) slow-fast axis | ~ Protists (bacterivores) slow-fast axis | -0.10 | 0.09 | -1.15 | .249 |
| Oribatid mites (omnivores) slow-fast axis | ~ land use intensity | 0.07 | 0.11 | 0.63 | .529 |

| | | | | | |
|---|---|---|---|---|---|
| Oribatid mites (omnivores) slow-fast axis | ~ Plant slow-fast axis | 0.00 | 0.10 | 0.05 | .963 |
| Oribatid mites (omnivores) slow-fast axis | ~ Bacteria and fungi slow-fast axis | 0.16 | 0.09 | 1.70 | .089 |
| Oribatid mites (omnivores) slow-fast axis | ~ Protists (bacterivores) slow-fast axis | -0.06 | 0.09 | -0.61 | .544 |
| Oribatid mites (omnivores) slow-fast axis | ~ Protists (secondary consumers) slow-fast axis | -0.04 | 0.10 | -0.40 | .687 |
| Collembola (omnivores) slow-fast axis | ~ land use intensity | -0.04 | 0.10 | -0.40 | .687 |
| Collembola (omnivores) slow-fast axis | ~ Plant slow-fast axis | -0.04 | 0.10 | -0.40 | .687 |
| Collembola (omnivores) slow-fast axis | ~ Bacteria and fungi slow-fast axis | -0.04 | 0.10 | -0.40 | .687 |
| Collembola (omnivores) slow-fast axis | ~ Protists (bacterivores) slow-fast axis | -0.04 | 0.10 | -0.40 | .687 |
| Collembola (omnivores) slow-fast axis | ~ Protists (secondary consumers) slow-fast axis | -0.04 | 0.10 | -0.40 | .687 |
| Arthropods (secondary consumers) slow-fast axis | ~ land use intensity | 0.08 | 0.09 | 0.89 | .376 |
| Arthropods (secondary consumers) slow-fast axis | ~ Oribatid mites slow-fast axis | 0.37 | 0.07 | 5.39 | .000 |
| Arthropods (secondary consumers) slow-fast axis | ~ Collembola slow-fast axis | -0.20 | 0.08 | -2.47 | .014 |
| Arthropods (secondary consumers) slow-fast axis | ~ Protists (bacterivores) slow-fast axis | -0.09 | 0.08 | -1.12 | .261 |
| Arthropods (secondary consumers) slow-fast axis | ~ Protists (secondary consumers) slow-fast axis | -0.05 | 0.08 | -0.55 | .581 |
| **Intercepts** | | | | | |
| Plant slow-fast axis | | -0.00 | 0.07 | -0.00 | 1.000 |
| Protists (primary consumers) slow-fast axis | | 0.00 | 0.07 | 0.00 | 1.000 |
| Bacteria and fungi fast-slow axis | | -0.00 | 0.06 | -0.00 | 1.000 |

| | | | | | |
|---|---|---|---|---|---|
| Protists (bacterivores) slow-fast axis | | 0.00 | 0.07 | 0.00 | 1.000 |
| Protists (secondary consumers) slow-fast axis | | 0.00 | 0.08 | 0.00 | 1.000 |
| Oribatid mites (omnivores) slow-fast axis | | 0.01 | 0.09 | 0.11 | .913 |
| Collembola (omnivores) slow-fast axis | | -0.01 | 0.08 | -0.10 | .921 |
| Above-ground arthropods (secondary consumers) slow-fast axis | | -0.01 | 0.07 | -0.07 | .944 |
| LUI | | 0 (fixed parameter) | | | |
| **Residual Variances** | | | | | |
| Plant slow-fast axis | | 0.74 | 0.09 | 8.43 | 0 |
| Protists (primary consumers) slow-fast axis | | 0.74 | 0.09 | 7.97 | .000 |
| Bacteria and fungi fast-slow axis | | 0.68 | 0.07 | 10.51 | .000 |
| Protists (bacterivores) slow-fast axis | | 0.71 | 0.08 | 9.41 | .000 |
| Protists (secondary consumers) slow-fast axis | | 0.81 | 0.11 | 7.10 | .000 |
| Oribatid mites (omnivores) slow-fast axis | | 0.83 | 0.09 | 9.14 | .000 |
| Collembola (omnivores) slow-fast axis | | 0.96 | 0.09 | 10.39 | .000 |
| Above-ground arthropods (secondary consumers) slow-fast axis | | 0.95 | 0.10 | 9.51 | .000 |
| LUI | | 1 (fixed parameter) | | | |
| **Residual Coariances** | | | | | |
| Protists (primary consumers) slow-fast axis | Bacteria and fungi fast-slow axis | 0.19 | 0.05 | 3.78 | .000 |
| Protists (primary consumers) slow-fast axis | | 0 (fixed parameter) | | | |

**Figure S 12.** Identification of ecosystem functions slow-fast axis. **Functions were not corrected for environmental covariates.** For all functions, high values are expected to be related to “fast” ecosystem functioning. Opposite response of dung decomposition compared to others is explained by the negative effect of Land-use intensity (especially mowing) on the abundance and activity of dung beetles (Frank et al., 2017).

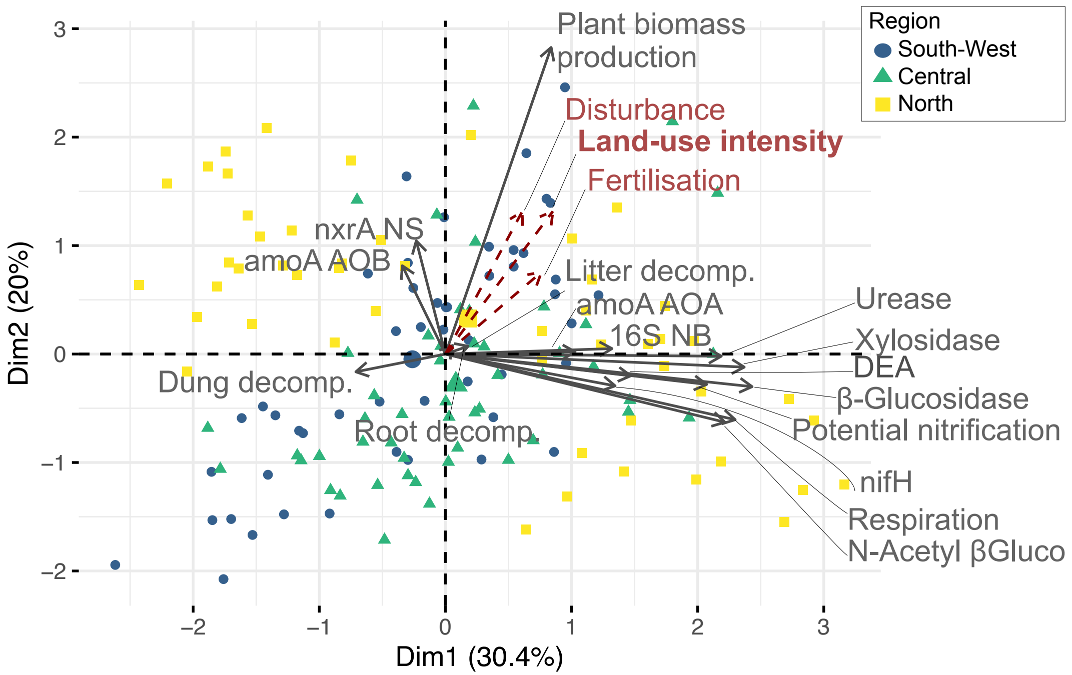

**Figure S 13.** Direct and indirect links between land- use intensity, functional traits slow-fast axis and ecosystem function slow-fast axis. **In contrast to Fig. S4, functions and community-level trait data were not corrected for environmental covariates.**

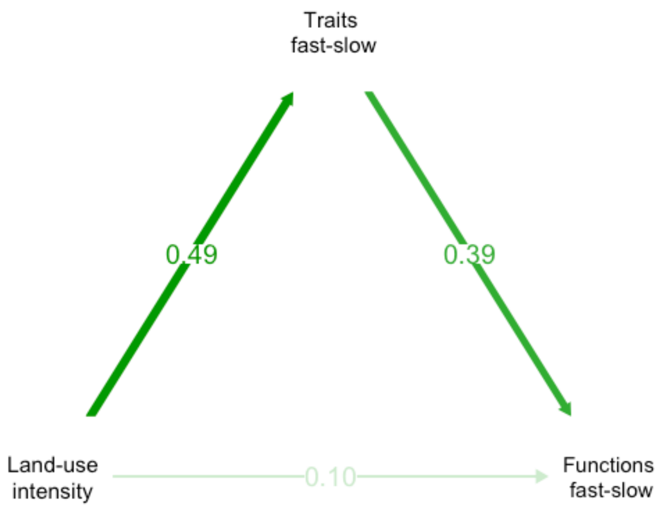

**Table S 17.** Comparison of the effect of multiple drivers on the ecosystem functions slow-fast axis, obtained from linear models with the function slow-fast axis as a response and the indicated variables as explanatory variables. **Functions and community-level trait data were not corrected for environmental covariates.** P-values were corrected for multiple testing within these different model (i.e. correction for false discovery rates, R function p.adjust, n = 5).

| **Model** | **Slope estimate (+/- standard error)** | **P-value** | **Adj $R^2$** |
|---|---|---|---|
| Functions slow-fast ~ plants slow-fast | 0.17 (0.06) | P < 0.001 | 5% |
| Functions slow-fast ~ microbial slow-fast | 0.25 (0.06) | P = 0.003 | 9% |
| Functions slow-fast ~ ecosystem-level slow-fast | 0.29 (0.05) | P < 0.001 | 19% |

| | | | |
|---|---|---|---|
| Functions slow-fast ~ taxonomic multidiversity | -0.37 (0.1) | P < 0.001 | 8% |
| Functions slow-fast ~ land-use intensity | 0.35 (0.1) | P < 0.001 | 8% |

**Table S 18.** Identification of guild-level slow-fast axes. **Analyses excluded all size- and body mass-related traits.**

Birds

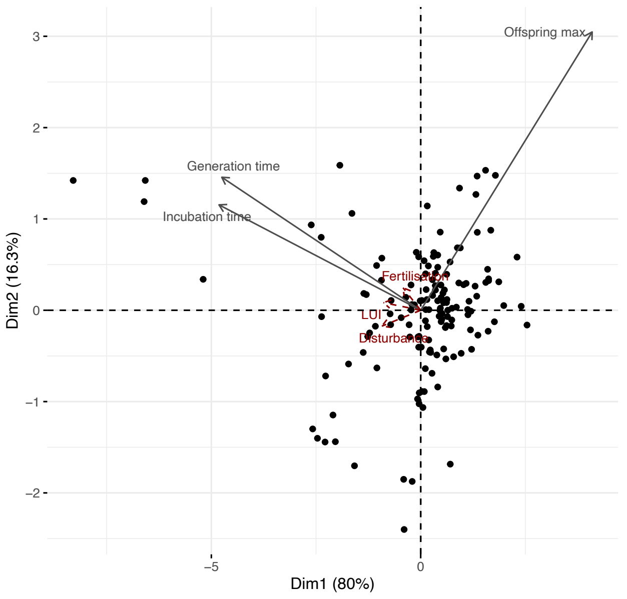

PC1 = slow-fast axis

Bats

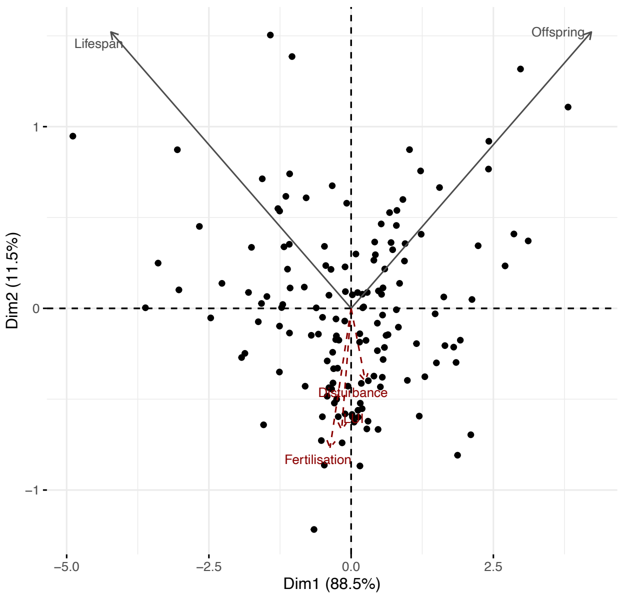

PC1 = slow-fast axis

| Arthropods, above-ground (secondary consumers) | Arthropods, above-ground (primary consumers) |
| --- | --- |
| Only one trait left (dispersal) -> no PCA | 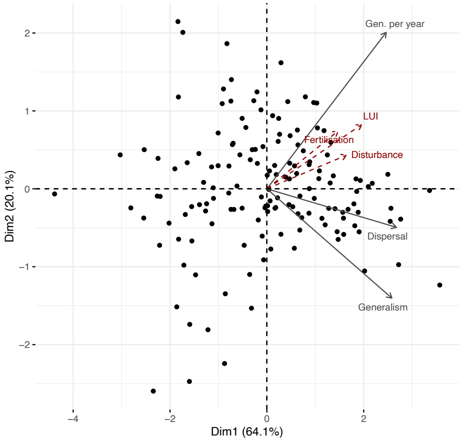<br>PC1 = slow-fast axis |
| Plants | Lepidoptera (primary consumers)<br>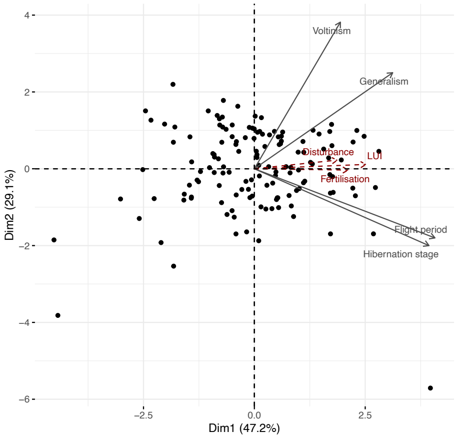 |

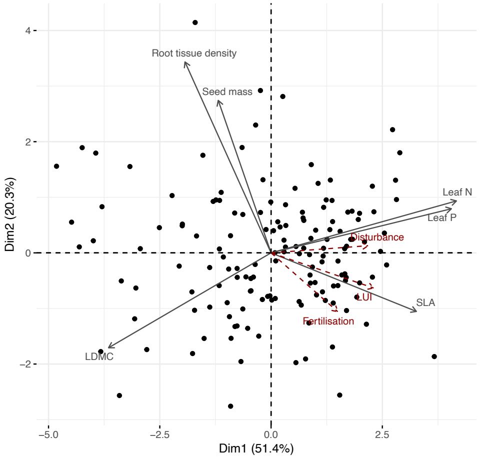

PC1 = slow-fast axis

PC1 = slow-fast axis

Fungi and bacteria

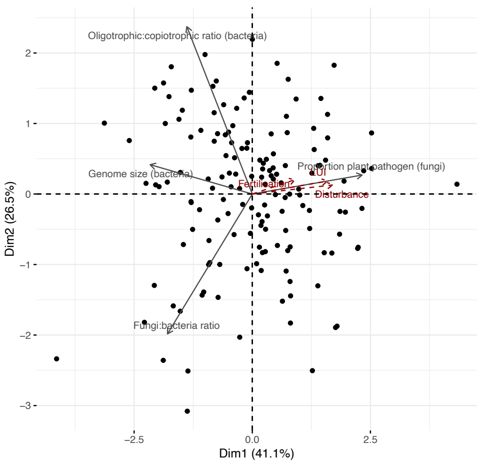

Protists, pathogens

Only one trait -> no PCA

| | |
|---|---|
| PC1 = slow-fast axis | |
| Protists, bacterivores<br><br>No trait left -> not included | Protists, secondary consumers<br><br>No trait left -> not included |
| Oribatid mites<br><br>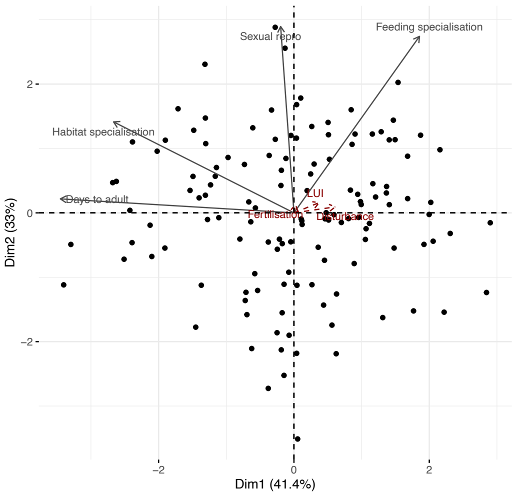<br><br>No slow-fast axis identified -> cannot be included in further analyses | Collembola<br><br>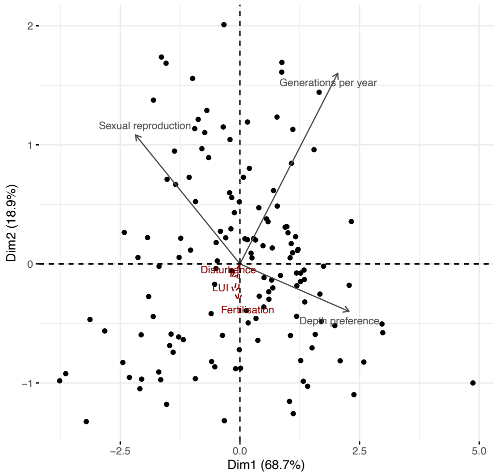<br><br>PC1 = slow-fast axis |
| Arthropods, below-ground (secondary consumers)<br><br>Only one trait left (dispersal) -> no PCA | |

**Figure S 14.** Synchronised slow-fast trait response of individual guilds is strongly related to land-use intensity **Analyses excluded all size- and body mass-related traits**. The variables included in the PCA are the slow-fast axes of each guild. Land-use intensity, added as a supplementary variable, was strongly associated with axis 1. Belowground guilds are shown in brown, aboveground guilds in blue.

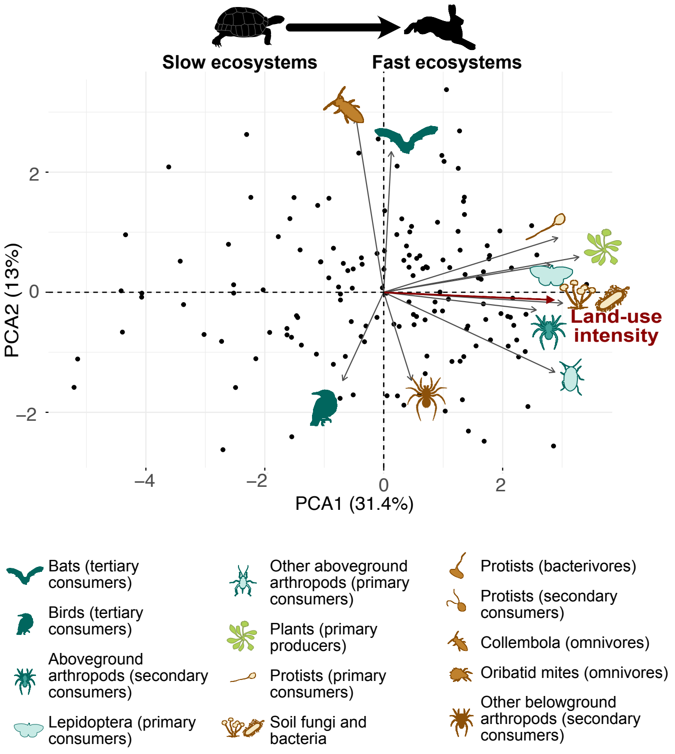

**Figure S 15.** Direct and trophically mediated effects of land-use intensity on the slow-fast axis of different trophic levels**. Analyses excludes size- and body mass- related traits.** a. Full SEMs including all guilds. Two independent models were fitted for below- and aboveground guilds; plants being included in both. b. Average direct, indirect and total LUI effects on each trophic level (averaged from the full SEM). c. Decreasing direct, indirect and total LUI effects with trophic level. Each dot represents the estimated effect (+/- standard error) of an individual guild in the full SEM.

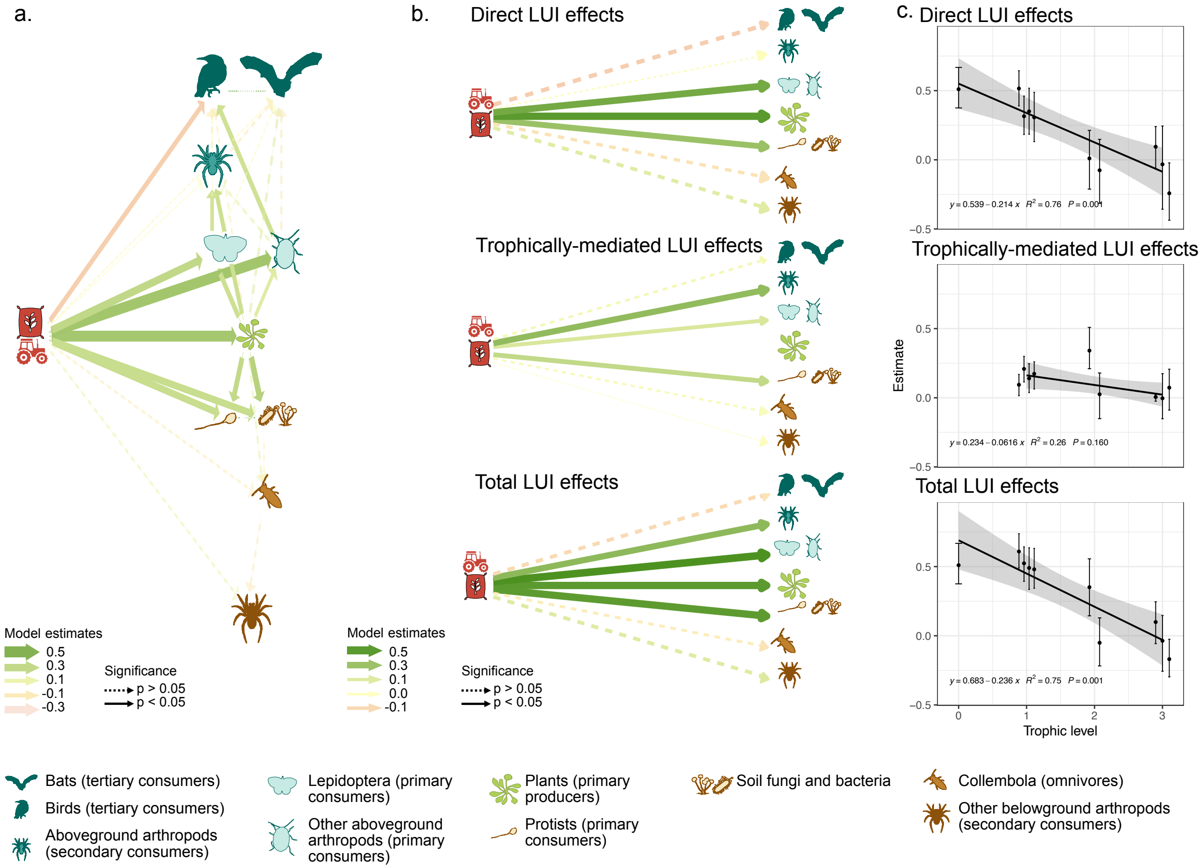

Falster, D.S., Duursma, R.A., Ishihara, M.I., Barneche, D.R., FitzJohn, R.G., Vårhammar, A., Aiba, M., Ando, M., Anten, N., Aspinwall, M.J., Baltzer, J.L., Baraloto, C., Battaglia, M., Battles, J.J., Bond-Lamberty, B., van Breugel, M., Camac, J., Claveau, Y., Coll, L., Dannoura, M., Delagrange, S., Domec, J.-C., Fatemi, F., Feng, W., Gargaglione, V., Goto, Y., Hagihara, A., Hall, J.S., Hamilton, S., Harja, D., Hiura, T., Holdaway, R., Hutley, L.S., Ichie, T., Jokela, E.J., Kantola, A., Kelly, J.W.G., Kenzo, T., King, D., Kloeppel, B.D., Kohyama, T., Komiyama, A., Laclau, J.-P., Lusk, C.H., Maguire, D.A., le Maire, G., Mäkelä, A., Markesteijn, L., Marshall, J., McCulloh, K., Miyata, I., Mokany, K., Mori, S., Myster, R.W., Nagano, M., Naidu, S.L., Nouvellon, Y., O'Grady, A.P., O'Hara, K.L., Ohtsuka, T., Osada, N., Osunkoya, O.O., Peri, P.L., Petritan, A.M., Poorter, L., Portsmuth, A., Potvin, C., Ransijn, J., Reid, D., Ribeiro, S.C., Roberts, S.D., Rodríguez, R., Saldaña-Acosta, A., Santa-Regina, I., Sasa, K., Selaya, N.G., Sillett, S.C., Sterck, F., Takagi, K., Tange, T., Tanouchi, H., Tissue, D., Umehara, T., Utsugi, H., Vadeboncoeur, M.A., Valladares, F., Vanninen, P., Wang, J.R., Wenk, E., Williams, R., de Aquino Ximenes, F., Yamaba, A., Yamada, T.,

Yamakura, T., Yanai, R.D., York, R.A., 2015. BAAD: a Biomass And Allometry Database for woody plants: *Ecological Archives* E096-128. Ecology 96, 1445–1445. https://doi.org/10.1890/14-1889.1

Fierer, N., Leff, J.W., Adams, B.J., Nielsen, U.N., Bates, S.T., Lauber, C.L., Owens, S., Gilbert, J.A., Wall, D.H., Caporaso, J.G., 2012. Cross-biome metagenomic analyses of soil microbial communities and their functional attributes. Proc. Natl. Acad. Sci. 109, 21390–21395. https://doi.org/10.1073/pnas.1215210110

Fiore-Donno, A.M., Bonkowski, M., 2019a. Cercozoa and Endomyxa (Rhizaria, protists), Illumina Sequences, all EPs, grassland and forest, 2011. v3. Biodiversity Exploratories Information System. Dataset. [WWW Document]. URL https://www.bexis.uni-jena.de/ddm/data/Showdata/24426

Fiore-Donno, A.M., Bonkowski, M., 2019b. Cercozoa and Endomyxa (Rhizaria, protists), Illumina Sequences, all EPs, grassland and forest, 2017. v3. Biodiversity Exploratories Information System. Dataset. [WWW Document]. URL https://www.bexis.uni-jena.de/ddm/data/Showdata/24466

Fiore-Donno, A.M., Richter-Heitmann, M., Bonkowski, T., 2020a. Functional traits of two important protistan lineages (Cercozoa and Endomyxa) reveal contrasting distributions of plant parasites and phagotrophs in temperate grassland and forest soils 14.

Fiore-Donno, A.M., Richter-Heitmann, T., Bonkowski, M., 2020b. Contrasting Responses of Protistan Plant Parasites and Phagotrophs to Ecosystems, Land Management and Soil Properties. Front. Microbiol. 11.

Fischer, M., Grassein, F., 2015. Litter decomposition (2013, all grassland EPs). v3. Biodiversity Exploratories Information System. Dataset. [WWW Document]. URL https://www.bexis.uni-jena.de/ddm/data/Showdata/18926

Fischer, M., Schäfer, D., Boch, S., Prati, D., 2017. Vegetation Records for 150 Grassland EPs in 2016, Header Data without Species Identities (incl. biomass). v2. Biodiversity Exploratories Information System. Dataset. [WWW Document]. URL https://www.bexis.uni-jena.de/ddm/data/Showdata/21187

Fonseca, C.R., Overton, J.McC., Collins, B., Westoby, M., 2000. Shifts in Trait-Combinations along Rainfall and Phosphorus Gradients. J. Ecol. 88, 964–977.

Frank, K., Blüthgen, N., 2017. Decomposition (dung removal in g on 300 EPs, season 2014/2015), Taxon: dung beetles (Invertebrates, Scarabaeoidea) - Dungwebs. v3. Biodiversity Exploratories Information System. Dataset. [WWW Document]. URL https://www.bexis.uni-jena.de/ddm/data/Showdata/21206

Frank, K., Hülsmann, M., Assmann, T., Schmitt, T., Blüthgen, N., 2017. Land use affects dung beetle communities and their ecosystem service in forests and grasslands. Agric. Ecosyst. Environ. 243, 114–122. https://doi.org/10.1016/j.agee.2017.04.010

Frawley, B.J., Best, L.B., 1992. Effects of mowing on breeding bird abundance and species composition in alfalfa fields. Biol. Conserv. 60, 222. https://doi.org/10.1016/0006-3207(92)91269-X

Freschet, G.T., Cornelissen, J.H.C., Van Logtestijn, R.S.P., Aerts, R., 2010. Evidence of the 'plant economics spectrum' in a subarctic flora. J. Ecol. 98, 362–373. https://doi.org/10.1111/j.1365-2745.2009.01615.x

Frostegård, Å., Bååth, E., Tunlio, A., 1993. Shifts in the structure of soil microbial communities in limed forests as revealed by phospholipid fatty acid analysis. Soil Biol. Biochem. 25, 723–730. https://doi.org/10.1016/0038-0717(93)90113-P

Frostegård, Å, Tunlid, A., Bååth, E., 1993. Phospholipid Fatty Acid Composition, Biomass, and Activity of Microbial Communities from Two Soil Types Experimentally Exposed to Different Heavy Metals. Appl. Environ. Microbiol. https://doi.org/10.1128/aem.59.11.3605-3617.1993

Frostegård, Å., Tunlid, A., Bååth, E., 1991. Microbial biomass measured as total lipid phosphate in soils of different organic content. J. Microbiol. Methods 14, 151–163. https://doi.org/10.1016/0167-7012(91)90018-L

Giarrizzo, E., Burrascano, S., Chiti, T., de Bello, F., Lepš, J., Zavattero, L., Blasi, C., 2017. Re-visiting historical semi-natural grasslands in the Apennines to assess patterns of changes in species composition and functional traits. Appl. Veg. Sci. 20, 247–258. https://doi.org/10.1111/avsc.12288

Goldmann, K., Buscot, F., Wubet, T., 2018. Abundant fungi on all EPs (from Soil Sampling Campain 2011) - merged grassland and forest data: taxonomic look-up table. Version 3. Biodiversity Exploratories Information System. (Dataset) [WWW Document]. URL https://www.bexis.uni-jena.de/ddm/data/Showdata/24306

Gonzalez-Akre, E., McShea, W., Bourg, N., Anderson-Teixera, K., 2015. Leaf traits data (SLA) for 56 woody species at the Smithsonian Conservation Biology Institute-ForestGEO Forest Dynamic Plot. [Data set]. Version 1.0. Front Royal, Virginia. USA.

Gos, P., Loucougaray, G., Colace, M.-P., Arnoldi, C., Gaucherand, S., Dumazel, D., Girard, L., Delorme, S., Lavorel, S., 2016. Relative contribution of soil, management and traits to co-variations of multiple ecosystem properties in grasslands. Oecologia 180, 1001–1013. https://doi.org/10.1007/s00442-016-3551-3

Gossner, M.M., Simons, N.K., Achtziger, R., Blick, T., Dorow, W.H.O., Dziock, F., Köhler, F., Rabitsch, W., Weisser, W.W., 2015. A summary of eight traits of Coleoptera, Hemiptera, Orthoptera and Araneae, occurring in grasslands in Germany. Sci. Data 2, 150013. https://doi.org/10.1038/sdata.2015.13

Green, W., 2002. USDA PLANTS Compilation, version 1 [WWW Document]. URL http://bricol.net/downloads/data/PLANTSdatabase/

Grime, J.P., 1979. Plant Strategies and Vegetation Processes. John Wiley & Sons Ltd, Chichester.

Grogan, P., 1998. Co2 Flux Measurement Using Soda Lime: Correction for Water Formed During Co2 Adsorption. Ecology 79, 1467–1468. https://doi.org/10.1890/0012-9658(1998)079[1467:CFMUSL]2.0.CO;2

Guy, A.L., Mischkolz, J.M., Lamb, E.G., 2013. Limited effects of simulated acidic deposition on seedling survivorship and root morphology of endemic plant taxa of the Athabasca Sand Dunes in well-watered greenhouse trials. Botany 91, 176–181. https://doi.org/10.1139/cjb-2012-0162

Han, W., Chen, Y., Zhao, F.-J., Tang, L., Jiang, R., Zhang, F., 2012. Floral, climatic and soil pH controls on leaf ash content in China's terrestrial plants: Ecological controls on leaf ash. Glob. Ecol. Biogeogr. 21, 376–382. https://doi.org/10.1111/j.1466-8238.2011.00677.x

Hänsel, F., Forteva, S., Wöllauer, S., Nauss, T., 2019. Öffentlich verfügbare Klimadaten der Exploratorien / Open Climate Data of the Exploratories Project. v4. Biodiversity Exploratories Information System. Dataset. [WWW Document]. URL https://www.bexis.uni-jena.de/tcd/PublicClimateData/Index

Hanson, H.I., Palmu, E., Birkhofer, K., Smith, H.G., Hedlund, K., 2016. Agricultural Land Use Determines the Trait Composition of Ground Beetle Communities. PLOS ONE 11, e0146329. https://doi.org/10.1371/journal.pone.0146329

Herz, K., Dietz, S., Haider, S., Jandt, U., Scheel, D., Bruelheide, H., 2017a. Drivers of intraspecific trait variation of grass and forb species in German meadows and pastures. J. Veg. Sci. 28, 705–716. https://doi.org/10.1111/jvs.12534

Herz, K., Dietz, S., Haider, S., Jandt, U., Scheel, D., Bruelheide, H., 2017b. Predicting individual plant performance in grasslands. Ecol. Evol. 7, 8958–8965. https://doi.org/10.1002/ece3.3393

Hodgson, J.G., Montserrat-Martí, G., Charles, M., Jones, G., Wilson, P., Shipley, B., Sharafi, M., Cerabolini, B.E.L., Cornelissen, J.H.C., Band, S.R., Bogard, A., Castro-Díez, P., Guerrero-Campo, J., Palmer, C., Pérez-Rontomé, M.C., Carter, G., Hynd, A., Romo-Díez, A., de Torres Espuny, L., Royo Pla, F., 2011. Is leaf dry matter content a better predictor of soil fertility than specific leaf area? Ann. Bot. 108, 1337–1345. https://doi.org/10.1093/aob/mcr225

Hoffmann, H., Schloter, M., Wilke, B.-M., 2007. Microscale-scale measurement of potential nitrification rates of soil aggregates. Biol. Fertil. Soils 44, 411–413. https://doi.org/10.1007/s00374-007-0227-5

Iversen, C.M., McCormack, M.L., Powell, A.S., Blackwood, C.B., Freschet, G.T., Kattge, J., Roumet, C., Stover, D.B., Soudzilovskaia, N.A., Valverde-Barrantes, O.J., Bodegom, P.M., Violle, C., 2017. A global Fine-Root Ecology Database to address below-ground challenges in plant ecology. New Phytol. 215, 15–26. https://doi.org/10.1111/nph.14486

Jung, K., Tschapka, M., 2016a. Bat activity in all Exploratories, summer 2009, using acoustic monitoring. v2. Biodiversity Exploratories Information System. Dataset. [WWW Document]. URL https://www.bexis.uni-jena.de/ddm/data/Showdata/19849

Jung, K., Tschapka, M., 2016b. Bat activity in all Exploratories, summer 2010, using acoustic monitoring. v2. Biodiversity Exploratories Information System. Dataset. [WWW Document]. URL https://www.bexis.uni-jena.de/ddm/data/Showdata/19850

Kandeler, E., Berner, D., Marhan, S., Boeddinghaus, R.S., 2017a. Soil enzyme activities of all grassland EPs, soil sampling campaign (SSC) 2011, SCALEMIC. v4. Biodiversity Exploratories Information System. Dataset [WWW Document]. URL https://www.bexis.uni-jena.de/ddm/data/Showdata/20246

Kandeler, E., Marhan, S., Berner, D., Boeddinghaus, R.S., 2017b. Microbial soil properties of all grassland EPs, soil sampling campaign (SSC) 2011, SCALEMIC. v3. Biodiversity Exploratories Information System. Dataset. [WWW Document]. URL https://www.bexis.uni-jena.de/ddm/data/Showdata/20250

Kandeler, E., Marhan, S., Berner, D., Boeddinghaus, R.S., 2017c. Microbial soil properties of all grassland EPs, soil sampling campaign (SSC) 2014, SCALEMIC. v3. Biodiversity Exploratories Information System. Dataset. [WWW Document]. URL https://www.bexis.uni-jena.de/ddm/data/Showdata/20251

Kattge, J., Díaz, S., Lavorel, S., Prentice, I.C., Leadley, P., Bönisch, G., Garnier, E., Westoby, M., Reich, P.B., Wright, I.J., Cornelissen, J.H.C., Violle, C., Harrison, S.P., Van BODEGOM, P.M., Reichstein, M., Enquist, B.J., Soudzilovskaia, N.A., Ackerly, D.D., Anand, M., Atkin, O., Bahn, M., Baker, T.R., Baldocchi, D., Bekker, R., Blanco, C.C., Blonder, B., Bond, W.J., Bradstock, R., Bunker, D.E., Casanoves, F., Cavender-Bares, J., Chambers, J.Q., Chapin Iii, F.S., Chave, J., Coomes, D., Cornwell, W.K., Craine, J.M., Dobrin, B.H., Duarte, L., Durka, W., Elser, J., Esser, G., Estiarte, M., Fagan, W.F., Fang, J., Fernández-Méndez, F., Fidelis, A., Finegan, B., Flores, O., Ford, H., Frank, D., Freschet, G.T., Fyllas, N.M., Gallagher, R.V., Green, W.A., Gutierrez, A.G., Hickler, T., Higgins, S.I., Hodgson, J.G., Jalili, A., Jansen, S., Joly, C.A., Kerkhoff, A.J.,

Kirkup, D., Kitajima, K., Kleyer, M., Klotz, S., Knops, J.M.H., Kramer, K., Kühn, I., Kurokawa, H., Laughlin, D., Lee, T.D., Leishman, M., Lens, F., Lenz, T., Lewis, S.L., Lloyd, J., Llusià, J., Louault, F., Ma, S., Mahecha, M.D., Manning, P., Massad, T., Medlyn, B.E., Messier, J., Moles, A.T., Müller, S.C., Nadrowski, K., Naeem, S., Niinemets, Ü., Nöllert, S., Nüske, A., Ogaya, R., Oleksyn, J., Onipchenko, V.G., Onoda, Y., Ordoñez, J., Overbeck, G., Ozinga, W.A., Patiño, S., Paula, S., Pausas, J.G., Peñuelas, J., Phillips, O.L., Pillar, V., Poorter, H., Poorter, L., Poschlod, P., Prinzing, A., Proulx, R., Rammig, A., Reinsch, S., Reu, B., Sack, L., Salgado-Negret, B., Sardans, J., Shiodera, S., Shipley, B., Siefert, A., Sosinski, E., Soussana, J.-F., Swaine, E., Swenson, N., Thompson, K., Thornton, P., Waldram, M., Weiher, E., White, M., White, S., Wright, S.J., Yguel, B., Zaehle, S., Zanne, A.E., Wirth, C., 2011. TRY – a global database of plant traits. Glob. Change Biol. 17, 2905–2935. https://doi.org/10.1111/j.1365-2486.2011.02451.x

Kattge, J., Knorr, W., Raddatz, T., Wirth, C., 2009. Quantifying photosynthetic capacity and its relationship to leaf nitrogen content for global-scale terrestrial biosphere models. Glob. Change Biol. 15, 976–991. https://doi.org/10.1111/j.1365-2486.2008.01744.x

Kazakou, E., Vile, D., Shipley, B., Gallet, C., Garnier, E., 2006. Co-Variations in Litter Decomposition, Leaf Traits and Plant Growth in Species from a Mediterranean Old-Field Succession. Funct. Ecol. 20, 21–30.

Keil, D., Niklaus, P.A., von Riedmatten, L.R., Boeddinghaus, R.S., Dormann, C.F., Scherer-Lorenzen, M., Kandeler, E., Marhan, S., 2015. Effects of warming and drought on potential N2O emissions and denitrifying bacteria abundance in grasslands with different land-use. FEMS Microbiol. Ecol. 91, fiv066. https://doi.org/10.1093/femsec/fiv066

Keith, H., Wong, S.C., 2006. Measurement of soil CO2 efflux using soda lime absorption: both quantitative and reliable. Soil Biol. Biochem. 38, 1121–1131. https://doi.org/10.1016/j.soilbio.2005.09.012

Kerkhoff, A.J., Fagan, W.F., Elser, J.J., Enquist, B.J., 2006. Phylogenetic and Growth Form Variation in the Scaling of Nitrogen and Phosphorus in the Seed Plants. Am. Nat. 168, E103–E122. https://doi.org/10.1086/507879

Kichenin, E., Wardle, D.A., Peltzer, D.A., Morse, C.W., Freschet, G.T., 2013. Contrasting effects of plant inter- and intraspecific variation on community-level trait measures along an environmental gradient. Funct. Ecol. 27, 1254–1261. https://doi.org/10.1111/1365-2435.12116

Kleyer, M., Bekker, R. m., Knevel, I. c., Bakker, J. p., Thompson, K., Sonnenschein, M., Poschlod, P., Van Groenendael, J. m., Klimeš, L., Klimešová, J., Klotz, S., Rusch, G. m., Hermy, M., Adriaens, D., Boedeltje, G., Bossuyt, B., Dannemann, A., Endels, P., Götzenberger, L., Hodgson, J. g., Jackel, A.-K., Kühn, I., Kunzmann, D., Ozinga, W. a., Römermann, C., Stadler, M., Schlegelmilch, J., Steendam, H. j., Tackenberg, O., Wilmann, B., Cornelissen, J. h. c., Eriksson, O., Garnier, E., Peco, B., 2008. The LEDA Traitbase: a database of life-history traits of the Northwest European flora. J. Ecol. 96, 1266–1274. https://doi.org/10.1111/j.1365-2745.2008.01430.x

Koele, N., Dickie, I.A., Oleksyn, J., Richardson, S.J., Reich, P.B., 2012. No globally consistent effect of ectomycorrhizal status on foliar traits. New Phytol. 196, 845–852. https://doi.org/10.1111/j.1469-8137.2012.04297.x

Konstantinidis, K.T., Tiedje, J.M., 2004. Trends between gene content and genome size in prokaryotic species with larger genomes. Proc. Natl. Acad. Sci. 101, 3160–3165. https://doi.org/10.1073/pnas.0308653100

Kovacevic, V., Schulz, S., Schloter, M., 2019. Nitrifiers Abundances at Grassland EPs. v4. Biodiversity Exploratories Information System. Dataset. [WWW Document]. URL https://www.bexis.uni-jena.de/ddm/data/Showdata/21547

Kühn, I., Durka, W., Klotz, S., 2004. BiolFlor — a new plant-trait database as a tool for plant invasion ecology. Divers. Distrib. 10, 363–365. https://doi.org/10.1111/j.1366-9516.2004.00106.x
La Pierre, K.J., Smith, M.D., 2015. Functional trait expression of grassland species shift with short- and long-term nutrient additions. Plant Ecol. 216, 307–318. https://doi.org/10.1007/s11258-014-0438-4
Lachaise, T., Bergmann, J., Rillig, M.C., van Kleunen, M., 2021. Below- and aboveground traits explain local abundance, and regional, continental and global occurrence frequencies of grassland plants. Oikos 130, 110–120. https://doi.org/10.1111/oik.07874
Laughlin, D.C., Fulé, P.Z., Huffman, D.W., Crouse, J., Laliberté, E., 2011. Climatic constraints on trait-based forest assembly. J. Ecol. 99, 1489–1499.
Laughlin, D.C., Leppert, J.J., Moore, M.M., Sieg, C.H., 2010. A multi-trait test of the leaf-height-seed plant strategy scheme with 133 species from a pine forest flora. Funct. Ecol. 24 493-501 493–501. https://doi.org/10.1111/j.1365-2435.2009.01672.x
Lavorel, S., Garnier, E., 2002. Predicting changes in community composition and ecosystem functioning from plant traits: revisiting the Holy Grail. Funct. Ecol. 16, 545–556. https://doi.org/10.1046/j.1365-2435.2002.00664.x
Le Provost, G., Manning, P., Schöning, I., Schrumpf, M., Weisser, W.W., Lorenzen, K., Vogt, J., Thiele, J., Westphal, C., Dewenter, I.S., 2021a. Aggregated environmental and land-use covariates of the 150 grassland EPs used in "Contrasting responses of above- and belowground diversity to multiple components of land-use intensity". v5. Biodiversity Exploratories Information System. Dataset. [WWW Document]. URL https://www.bexis.uni-jena.de/ddm/data/Showdata/31018
Le Provost, G., Thiele, J., Westphal, C., Penone, C., Allan, E., Neyret, M., van der Plas, F., Ayasse, M., Bardgett, R.D., Birkhofer, K., Boch, S., Bonkowski, M., Buscot, F., Feldhaar, H., Gaulton, R., Goldmann, K., Gossner, M.M., Klaus, V.H., Kleinebecker, T., Krauss, J., Renner, S., Scherreiks, P., Sikorski, J., Baulechner, D., Blüthgen, N., Bolliger, R., Börschig, C., Busch, V., Chisté, M., Fiore-Donno, A.M., Fischer, M., Arndt, H., Hoelzel, N., John, K., Jung, K., Lange, M., Marzini, C., Overmann, J., Paŝalić, E., Perović, D.J., Prati, D., Schäfer, D., Schöning, I., Schrumpf, M., Sonnemann, I., Steffan-Dewenter, I., Tschapka, M., Türke, M., Vogt, J., Wehner, K., Weiner, C., Weisser, W., Wells, K., Werner, M., Wolters, V., Wubet, T., Wurst, S., Zaitsev, A.S., Manning, P., 2021b. Contrasting responses of above- and belowground diversity to multiple components of land-use intensity. Nat. Commun. 12, 3918. https://doi.org/10.1038/s41467-021-23931-1
Leff, J.W., Jones, S.E., Prober, S.M., Barberán, A., Borer, E.T., Firn, J.L., Harpole, W.S., Hobbie, S.E., Hofmockel, K.S., Knops, J.M.H., McCulley, R.L., La Pierre, K., Risch, A.C., Seabloom, E.W., Schütz, M., Steenbock, C., Stevens, C.J., Fierer, N., 2015. Consistent responses of soil microbial communities to elevated nutrient inputs in grasslands across the globe. Proc. Natl. Acad. Sci. 112, 10967–10972. https://doi.org/10.1073/pnas.1508382112
Lekberg, Y., Arnillas, C.A., Borer, E.T., Bullington, L.S., Fierer, N., Kennedy, P.G., Leff, J.W., Luis, A.D., Seabloom, E.W., Henning, J.A., 2021. Nitrogen and phosphorus fertilization consistently favor pathogenic over mutualistic fungi in grassland soils. Nat. Commun. 12, 3484. https://doi.org/10.1038/s41467-021-23605-y
Lepiforum e.V. [WWW Document], n.d. URL https://lepiforum.org/ (accessed 12.7.21).
Lhotsky, B., Csecserits, A., Kovács, B., Botta-Dukát, Z., 2016. New plant trait records of the Hungarian flora. Acta Bot. Hung. 58, 397–400. https://doi.org/10.1556/abot.58.2016.3-4.8

Li, Y., Shipley, B., 2018. Community divergence and convergence along experimental gradients of stress and disturbance. Ecology 99, 775–781. https://doi.org/10.1002/ecy.2162

Liebergesell, M., Reu, B., Stahl, U., Freiberg, M., Welk, E., Kattge, J., Cornelissen, J.H.C., Peñuelas, J., Wirth, C., 2016. Functional Resilience against Climate-Driven Extinctions – Comparing the Functional Diversity of European and North American Tree Floras. PLOS ONE 11, e0148607. https://doi.org/10.1371/journal.pone.0148607

Lin, Y.-S., Medlyn, B.E., Duursma, R.A., Prentice, I.C., Wang, H., Baig, S., Eamus, D., de Dios, V.R., Mitchell, P., Ellsworth, D.S., de Beeck, M.O., Wallin, G., Uddling, J., Tarvainen, L., Linderson, M.-L., Cernusak, L.A., Nippert, J.B., Ocheltree, T.W., Tissue, D.T., Martin-StPaul, N.K., Rogers, A., Warren, J.M., De Angelis, P., Hikosaka, K., Han, Q., Onoda, Y., Gimeno, T.E., Barton, C.V.M., Bennie, J., Bonal, D., Bosc, A., Löw, M., Macinins-Ng, C., Rey, A., Rowland, L., Setterfield, S.A., Tausz-Posch, S., Zaragoza-Castells, J., Broadmeadow, M.S.J., Drake, J.E., Freeman, M., Ghannoum, O., Hutley, L.B., Kelly, J.W., Kikuzawa, K., Kolari, P., Koyama, K., Limousin, J.-M., Meir, P., Lola da Costa, A.C., Mikkelsen, T.N., Salinas, N., Sun, W., Wingate, L., 2015. Optimal stomatal behaviour around the world. Nat. Clim. Change 5, 459–464. https://doi.org/10.1038/nclimate2550

Liu, X., Zhang, L., Huang, M., Zhou, S., 2021. Plant diversity promotes soil fungal pathogen richness under fertilization in an alpine meadow. J. Plant Ecol. 14, 323–336. https://doi.org/10.1093/jpe/rtaa099

Louault, F., Pillar, V. d., Aufrère, J., Garnier, E., Soussana, J.-F., 2005. Plant traits and functional types in response to reduced disturbance in a semi-natural grassland. J. Veg. Sci. 16, 151–160. https://doi.org/10.1111/j.1654-1103.2005.tb02350.x

Lüftenegger, G., Foissner, W., Adam, H., 1985. r-and K-selection in soil ciliates: a field and experimental approach. Oecologia 66, 574–579. https://doi.org/10.1007/BF00379352

Lukeš, P., Stenberg, P., Rautiainen, M., Mõttus, M., Vanhatalo, K.M., 2013. Optical properties of leaves and needles for boreal tree species in Europe. Remote Sens. Lett. 4, 667–676. https://doi.org/10.1080/2150704X.2013.782112

MacDonald, M.A., 2006. The indirect effects of increased nutrient inputs on birds in the UK: a review 220.

Madin, J.S., Nielsen, D.A., Brbic, M., Corkrey, R., Danko, D., Edwards, K., Engqvist, M.K.M., Fierer, N., Geoghegan, J.L., Gillings, M., Kyrpides, N.C., Litchman, E., Mason, C.E., Moore, L., Nielsen, S.L., Paulsen, I.T., Price, N.D., Reddy, T.B.K., Richards, M.A., Rocha, E.P.C., Schmidt, T.M., Shaaban, H., Shukla, M., Supek, F., Tetu, S.G., Vieira-Silva, S., Wattam, A.R., Westfall, D.A., Westoby, M., 2020. A synthesis of bacterial and archaeal phenotypic trait data. Sci. Data 7, 170. https://doi.org/10.1038/s41597-020-0497-4

Maire, V., Wright, I.J., Prentice, I.C., Batjes, N.H., Bhaskar, R., Van Bodegom, P.M., Cornwell, W.K., Ellsworth, D., Niinemets, Ü., Ordoñez, A., Reich, P.B., Santiago, L.S., 2016. Data from: Global effects of soil and climate on leaf photosynthetic traits and rates. https://doi.org/10.5061/DRYAD.J42M7

Maire, V., Wright, I.J., Prentice, I.C., Batjes, N.H., Bhaskar, R., van Bodegom, P.M., Cornwell, W.K., Ellsworth, D., Niinemets, Ü., Ordonez, A., Reich, P.B., Santiago, L.S., 2015. Global effects of soil and climate on leaf photosynthetic traits and rates. Glob. Ecol. Biogeogr. 24, 706–717. https://doi.org/10.1111/geb.12296

Mangels, J., Blüthgen, N., 2017. Life history traits of moths. v2. Biodiversity Exploratories Information System. Dataset. [WWW Document]. URL https://www.bexis.uni-jena.de/ddm/data/Showdata/21228

Mangels, J., Fiedler, K., Schneider, F.D., Blüthgen, N., 2017. Diversity and trait composition of moths respond to land-use intensification in grasslands: generalists replace specialists. Biodivers. Conserv. 26, 3385–3405. https://doi.org/10.1007/s10531-017-1411-z

Martin, A.R., Hale, C.E., Cerabolini, B.E.L., Cornelissen, J.H.C., Craine, J., Gough, W.A., Kattge, J., Tirona, C.K.F., 2018. Inter- and intraspecific variation in leaf economic traits in wheat and maize. AoB PLANTS 10, ply006. https://doi.org/10.1093/aobpla/ply006

Marx, M.-C., Wood, M., Jarvis, S.C., 2001. A microplate fluorimetric assay for the study of enzyme diversity in soils. Soil Biol. Biochem. 33, 1633–1640. https://doi.org/10.1016/S0038-0717(01)00079-7

McMahon, B.J., Doyle, S., Gray, A., Kelly, S.B.A., Redpath, S.M., 2020. European bird declines: Do we need to rethink approaches to the management of abundant generalist predators? J. Appl. Ecol. 57, 1885–1890. https://doi.org/10.1111/1365-2664.13695

Medlyn, B.E., Badeck, F.W., De Pury, D.G.G., Barton, C.V.M., Broadmeadow, M., Ceulemans, R., De Angelis, P., Forstreuter, M., Jach, M.E., Kellomäki, S., Laitat, E., Marek, M., Philippot, S., Rey, A., Strassemeyer, J., Laitinen, K., Liozon, R., Portier, B., Roberntz, P., Wang, K., Jarvis, P.G., 1999. Effects of elevated [CO2] on photosynthesis in European forest species: a meta-analysis of model parameters. Plant Cell Environ. 22, 1475–1495. https://doi.org/10.1046/j.1365-3040.1999.00523.x

Meir, P., Kruijt, B., Broadmeadow, M., Barbosa, E., Kull, O., Carswell, F., Nobre, A., Jarvis, P.G., 2002. Acclimation of photosynthetic capacity to irradiance in tree canopies in relation to leaf nitrogen concentration and leaf mass per unit area. Plant Cell Environ. 25, 343–357. https://doi.org/10.1046/j.0016-8025.2001.00811.x

Mencuccini, M., 2003. The ecological significance of long-distance water transport: short-term regulation, long-term acclimation and the hydraulic costs of stature across plant life forms. Plant Cell Environ. 26, 163–182. https://doi.org/10.1046/j.1365-3040.2003.00991.x

Middleton-Welling, J., Dapporto, L., García-Barros, E., Wiemers, M., Nowicki, P., Plazio, E., Bonelli, S., Zaccagno, M., Šašić, M., Liparova, J., Schweiger, O., Harpke, A., Musche, M., Settele, J., Schmucki, R., Shreeve, T., 2020. A new comprehensive trait database of European and Maghreb butterflies, Papilionoidea. Sci. Data 7, 351. https://doi.org/10.1038/s41597-020-00697-7

Moles, A.T., Falster, D.S., Leishman, M.R., Westoby, M., 2004. Small-seeded species produce more seeds per square metre of canopy per year, but not per individual per lifetime. J. Ecol. 92, 384–396. https://doi.org/10.1111/j.0022-0477.2004.00880.x

Moretti, M., Legg, C., 2009. Combining plant and animal traits to assess community functional responses to disturbance. Ecography 32, 299–309. https://doi.org/10.1111/j.1600-0587.2008.05524.x

Neff, F., Resch, M.C., Marty, A., Rolley, J.D., Schütz, M., Risch, A.C., Gossner, M.M., 2020. Long-term restoration success of insect herbivore communities in semi-natural grasslands: a functional approach. Ecol. Appl. 30. https://doi.org/doi.org/10.1002/eap.2133

Neyret, M., Lachaise, T., van Kleunen, M., Bergmann, J., Manning, P., 2021. Community-weighted mean for plant below-ground traits - 300 forest and grassland EPs - 2008-2018. v3. Biodiversity Exploratories Information System. Dataset. [WWW Document]. URL https://www.bexis.uni-jena.de/ddm/data/Showdata/27608

Nickel, H., Remane, R., 2002. Check list of the planthoppers and leafhoppers of Germany, with notes on food plants, diet width, life cycles, geographic range and conservation status (Hemiptera, Fulgoromorpha and Cicadomorpha). Beitr. Zur Zikadenkunde 5.

NRCS: The PLANTS Database, 2009. . National Plant Data Center: Baton Rouge, LA 70874-74490 USA.

Obermann, J., Sikorski, J., 2019. 16S rRNA gene (V3 region) RNA-based analysis of microbial soil communities at sequence variant level in 148 grassland EP plots, using QIIME2-based bioinformatics, 2011. v2. Biodiversity Exploratories Information System. Dataset [WWW Document]. URL https://www.bexis.uni-jena.de/ddm/data/Showdata/24866

Ollivier, J., Schacht, D., Kindler, R., Groeneweg, J., Engel, M., Wilke, B.-M., Kleineidam, K., Schloter, M., 2013. Effects of repeated application of sulfadiazine-contaminated pig manure on the abundance and diversity of ammonia and nitrite oxidizers in the root-rhizosphere complex of pasture plants under field conditions. Front. Microbiol. 4, 22. https://doi.org/10.3389/fmicb.2013.00022

Onoda, Y., Wright, I.J., Evans, J.R., Hikosaka, K., Kitajima, K., Niinemets, Ü., Poorter, H., Tosens, T., Westoby, M., 2017. Physiological and structural tradeoffs underlying the leaf economics spectrum. New Phytol. 214, 1447–1463. https://doi.org/10.1111/nph.14496

Ordoñez, J.C., van Bodegom, P.M., Witte, J.M., Bartholomeus, R.P., van Hal, J.R., Aerts, R., 2010. Plant Strategies in Relation to Resource Supply in Mesic to Wet Environments: Does Theory Mirror Nature? Am. Nat. 175, 225–239. https://doi.org/10.1086/649582

Pahl, A.T., Kollmann, J., Mayer, A., Haider, S., 2013. No evidence for local adaptation in an invasive alien plant: field and greenhouse experiments tracing a colonization sequence. Ann. Bot. 112, 1921–1930. https://doi.org/10.1093/aob/mct246

Paine, C.E.T., Amissah, L., Auge, H., Baraloto, C., Baruffol, M., Bourland, N., Bruelheide, H., Daïnou, K., de Gouvenain, R.C., Doucet, J.-L., Doust, S., Fine, P.V.A., Fortunel, C., Haase, J., Holl, K.D., Jactel, H., Li, X., Kitajima, K., Koricheva, J., Martínez-Garza, C., Messier, C., Paquette, A., Philipson, C., Piotto, D., Poorter, L., Posada, J.M., Potvin, C., Rainio, K., Russo, S.E., Ruiz-Jaen, M., Scherer-Lorenzen, M., Webb, C.O., Wright, S.J., Zahawi, R.A., Hector, A., 2015. Globally, functional traits are weak predictors of juvenile tree growth, and we do not know why. J. Ecol. 103, 978–989. https://doi.org/10.1111/1365-2745.12401

Paula, S., Arianoutsou, M., Kazanis, D., Tavsanoglu, Ç., Lloret, F., Buhk, C., Ojeda, F., Luna, B., Moreno, J.M., Rodrigo, A., Espelta, J.M., Palacio, S., Fernández-Santos, B., Fernandes, P.M., Pausas, J.G., 2009. Fire-related traits for plant species of the Mediterranean Basin. Ecology 90, 1420–1420. https://doi.org/10.1890/08-1309.1

Peco, B., de Pablos, I., Traba, J., Levassor, C., 2005. The effect of grazing abandonment on species composition and functional traits: the case of dehesa grasslands. Basic Appl. Ecol. 6, 175–183. https://doi.org/10.1016/j.baae.2005.01.002

Penone, C., Allan, E., Boch, S., Bonkowski, M., Goldmann, K., Gossner, M.M., Wolters, V., 2021a. Assembled species information from grassland EPs (2008-2020) for multidiversity synthesis - November 2020. v3. Biodiversity Exploratories Information System. Dataset. [WWW Document]. URL https://www.bexis.uni-jena.de/ddm/data/Showdata/27706

Penone, C., Allan, E., Boch, S., Bonkowski, M., Goldmann, K., Gossner, M.M., Wolters, V., Blüthgen, N., Chisté, M., Feldhaar, H., Fischer, M., Jung, K., Krauss, J., Sikorski, J., Teuscher, M., Tschapka, M., Wurst, S., 2021b. Assembled RAW diversity from grassland EPs (2008-2020) for multidiversity synthesis - November 2020. v2. Biodiversity Exploratories Information System. Dataset. [WWW Document]. URL https://www.bexis.uni-jena.de/ddm/data/Showdata/27707

Penone, C., Renner, S., Teuscher, M., Fischer, M., 2022. Bird traits for all species in the Exploratories EPs [WWW Document]. URL https://www.bexis.uni-jena.de/ddm/data/Showdata/31368

Petersen, H., 1980. Population dynamic and metaboliccharacterization of Collembola species in a beechforest ecosystem., in: Soil Biology Asrelated to Land Use Practices. Presented at the roceedings of the VIIinternational soil zoology colloquium of the ISSS, D indal, D.L. (Ed), Syracuse, pp. 806-833Pe.

Pianka, E.R., 1970. On r- and K-Selection. Am. Nat. 104, 592–597.

Pigot, A.L., Sheard, C., Miller, E.T., Bregman, T.P., Freeman, B.G., Roll, U., Seddon, N., Trisos, C.H., Weeks, B.C., Tobias, J.A., 2020. Macroevolutionary convergence connects morphological form to ecological function in birds. Nat. Ecol. Evol. 4, 230–239. https://doi.org/10.1038/s41559-019-1070-4

Prati, D., Fischer, M., Minker, J., Schmitt, B., 2013. Measurement of biomass (2013, all grassland EPs). v2. Biodiversity Exploratories Information System. Dataset. [WWW Document]. URL https://www.bexis.uni-jena.de/ddm/data/Showdata/16786

Prentice, I.C., Meng, T., Wang, H., Harrison, S.P., Ni, J., Wang, G., 2011. Evidence of a universal scaling relationship for leaf CO2 drawdown along an aridity gradient. New Phytol. 190, 169–180. https://doi.org/10.1111/j.1469-8137.2010.03579.x

Preston, K.A., Cornwell, W.K., Denoyer, J.L., 2006. Wood density and vessel traits as distinct correlates of ecological strategy in 51 California coast range angiosperms. New Phytol. 170, 807–818. https://doi.org/10.1111/j.1469-8137.2006.01712.x

Price, C.A., Enquist, B.J., 2007. Scaling Mass and Morphology in Leaves: An Extension of the WBE Model. Ecology 88, 1132–1141.

Quested, H.M., Cornelissen, J.H.C., Press, M.C., Callaghan, T.V., Aerts, R., Trosien, F., Riemann, P., Gwynn-Jones, D., Kondratchuk, A., Jonasson, S.E., 2003. Decomposition of Sub-Arctic Plants with Differing Nitrogen Economies: A Functional Role for Hemiparasites. Ecology 84, 3209–3221.

Renner, S.C., Hoesel, W. van, 2017. Ecological and Functional Traits in 99 Bird Species over a Large-Scale Gradient in Germany. Data 2, 12. https://doi.org/10.3390/data2020012

Rolo, V., López-Díaz, M.L., Moreno, G., 2012. Shrubs affect soil nutrients availability with contrasting consequences for pasture understory and tree overstory production and nutrient status in Mediterranean grazed open woodlands. Nutr. Cycl. Agroecosystems 93, 89–102. https://doi.org/10.1007/s10705-012-9502-4

Ruess, L., Chamberlain, P.M., 2010. The fat that matters: Soil food web analysis using fatty acids and their carbon stable isotope signature. Soil Biol. Biochem. 42, 1898–1910. https://doi.org/10.1016/j.soilbio.2010.07.020

Sandel, B., Corbin, J.D., Krupa, M., 2011. Using plant functional traits to guide restoration: A case study in California coastal grassland. Ecosphere 2, art23. https://doi.org/10.1890/ES10-00175.1

Saulich, A., Musolin, D., 2021. Seasonal Development of Plant Bugs (Heteroptera, Miridae): Subfamily Mirinae, Tribe Stenodemini. Entomol. Rev. 101, 147–161. https://doi.org/10.1134/S0013873821020019

Schäfer, D., Fischer, M., Klaus, V., Busch, V., 2016a. Vegetation Records for 150 Grassland EPs in 2014, Header Data without Species Identities (incl. biomass). v2. Biodiversity Exploratories Information System. Dataset. [WWW Document]. URL https://www.bexis.uni-jena.de/ddm/data/Showdata/19807
Schäfer, D., Fischer, M., Klaus, V., Busch, V., 2016b. Vegetation Records for 150 Grassland EPs in 2015, Header Data without Species Identities (incl. biomass). v2. Biodiversity Exploratories Information System. Dataset. [WWW Document]. URL https://www.bexis.uni-jena.de/ddm/data/Showdata/19809
Schäfer, D., Prati, D., Fischer, M., 2018. Vegetation Records for 150 Grassland EPs in 2017, Header Data without Species Identities (incl. biomass). v2. Biodiversity Exploratories Information System. Dataset [WWW Document]. URL https://www.bexis.uni-jena.de/ddm/data/Showdata/23486
Schall, O., Petermann, R., 2014. National Report on Bat Conservation in the Federal Republic of Germany 2010-2013. Federal Ministry for the Environment, Nature Conservation, Building and Nuclear Safety (BMUB).
Schenk, N., Penone, C., Allan, E., Hamer, U., Oelmann, Y., Westphal, C., Schöning, I., Stempfhuber, B., Schrumpf, M., Rillig, M.C., Chisté, M., Weisser, W., Freitag, M., Schloter, M., Marhan, S., Blüthgen, N., Busch, V., Gossner, M.M., Fischer, M., 2021. Assembled ecosystem measures from grassland EPs (2008-2018) for multifunctionality synthesis - June 2020. Biodiversity Exploratories Information System. Dataset. [WWW Document]. URL https://www.bexis.uni-jena.de/ddm/data/Showdata/27087
Scherer-Lorenzen, M., Schulze, E.-D., Don, A., Schumacher, J., Weller, E., 2007. Exploring the functional significance of forest diversity: A new long-term experiment with temperate tree species (BIOTREE). Perspect. Plant Ecol. Evol. Syst. 9, 53–70. https://doi.org/10.1016/j.ppees.2007.08.002
Schinner, F., Öhlinger, R., Kandeler, E., Margesin, R., 1996. Methods in Soil Biology, 1st edition. ed. Springer, Berlin Heidelberg.
Schmitt, B., Prati, D., Fischer, M., 2011. Measurement of biomass (2010, all grassland EPs). v2. Biodiversity Exploratories Information System. Dataset [WWW Document]. URL https://www.bexis.uni-jena.de/ddm/data/Showdata/12706
Schmitt, B., Prati, D., Fischer, M., Minker, J., 2012a. Measurement of biomass (2011, all grassland EPs). v3. Biodiversity Exploratories Information System. Dataset. [WWW Document]. URL https://www.bexis.uni-jena.de/ddm/data/Showdata/14346
Schmitt, B., Prati, D., Fischer, M., Minker, J., 2012b. Measurement of biomass (2012, all grassland EPs). v2. Biodiversity Exploratories Information System. Dataset. [WWW Document]. URL https://www.bexis.uni-jena.de/ddm/data/Showdata/14986
Schmitt, M., Mehltreter, K., Sundue, M., Testo, W., Watanabe, T., Jansen, S., 2017. The evolution of aluminum accumulation in ferns and lycophytes. Am. J. Bot. 104, 573–583. https://doi.org/10.3732/ajb.1600381
Schöning, I., Heublein, J., Klötzing, T., Schrumpf, M., Trumbore, S., 2013. MinSoil 2011 - Soil Texture. v3. Biodiversity Exploratories Information System. Dataset. [WWW Document]. URL https://www.bexis.uni-jena.de/ddm/data/Showdata/14686
Schöning, I., Solly, E., Klötzing, T., Schrumpf, M., Trumbore, S., 2015a. Mineral soil pH values of all experimental plots (EP) of the Biodiversity Exploratories project from 2011, Soil (core project). v8. Biodiversity Exploratories Information System. Dataset. [WWW Document]. URL https://www.bexis.uni-jena.de/ddm/data/Showdata/14447
Schöning, I., Ying Gan, H., Heublein, J., Klötzing, T., Schrumpf, M., Trumbore, S., 2015b. MinSoil 2014 - Soil pH. v3. Biodiversity Exploratories Information System. Dataset. [WWW Document]. URL https://www.bexis.uni-jena.de/ddm/data/Showdata/19067

Schweingruber, F.H., Landolt, W., 2005. The Xylem Database. Updated. Swiss Federal Research Institute WSL.
Sheremetev, S.N., 2005. Herbs on the soil moisture gradient (water relations and the structural-functional organization) (in Russian). KMV, Moscow.
Sibly, R.M., Witt, C.C., Wright, N.A., Venditti, C., Jetz, W., Brown, J.H., 2012. Energetics, lifestyle, and reproduction in birds. Proc. Natl. Acad. Sci. 109, 10937–10941. https://doi.org/10.1073/pnas.1206512109
Siefert, A., Fridley, J.D., Ritchie, M.E., 2014. Community Functional Responses to Soil and Climate at Multiple Spatial Scales: When Does Intraspecific Variation Matter? PLOS ONE 9, e111189. https://doi.org/10.1371/journal.pone.0111189
Sikorski, J., Overmann, J., Marzini, C., 2019. 16S rRNA gene (V3 region) RNA-based analysis of microbial soil communities at sequence variant level in 150 grassland EP plots, using QIIME2-based bioinformatics, 2014. v3. Biodiversity Exploratories Information System. Dataset. [WWW Document]. URL https://www.bexis.uni-jena.de/ddm/data/Showdata/25066
Simons, N.K., Weisser, W.W., Gossner, M.M., 2016. Multi-taxa approach shows consistent shifts in arthropod functional traits along grassland land-use intensity gradient. Ecology 15-0616.1. https://doi.org/10.1890/15-0616.1
Smith, M.S., Tiedje, J.M., 1979. Phases of denitrification following oxygen depletion in soil. Soil Biol. Biochem. 11, 261–267. https://doi.org/10.1016/0038-0717(79)90071-3
Smith, N.G., Dukes, J.S., 2017. LCE: leaf carbon exchange data set for tropical, temperate, and boreal species of North and Central America. Ecology 98, 2978–2978. https://doi.org/10.1002/ecy.1992
Smith, S.W., Woodin, S.J., Pakeman, R.J., Johnson, D., Wal, R., 2014. Root traits predict decomposition across a landscape-scale grazing experiment. New Phytol. 203, 851–862. https://doi.org/10.1111/nph.12845
Socher, S.A., Prati, D., Boch, S., Müller, J., Klaus, V.H., Hölzel, N., Fischer, M., 2012. Direct and productivity-mediated indirect effects of fertilization, mowing and grazing on grassland species richness. J. Ecol. 100, 1391–1399. https://doi.org/10.1111/j.1365-2745.2012.02020.x
Solly, E., Schöning, I., 2013. MinSoil 2011 Root decomposition. v2. Biodiversity Exploratories Information System. Dataset. [WWW Document]. URL https://www.bexis.uni-jena.de/ddm/data/Showdata/16666
Spasojevic, M.J., Suding, K.N., 2012. Inferring community assembly mechanisms from functional diversity patterns: the importance of multiple assembly processes. J. Ecol. 100, 652–661. https://doi.org/10.1111/j.1365-2745.2011.01945.x
Spasojevic, M.J., Turner, B.L., Myers, J.A., 2016. When does intraspecific trait variation contribute to functional beta-diversity? J. Ecol. 104, 487–496. https://doi.org/10.1111/1365-2745.12518
Staab, M., Simons, N.K., Gossner, M.M., Weisser, W.W., Blüthgen, N., 2022. Body size and life-history traits of arthropod species. v7. Biodiversity Exploratories Information System. Dataset. [WWW Document]. URL https://www.bexis.uni-jena.de/ddm/data/Showdata/31122
Stempfhuber, B., Schloter, M., 2017. Abundance of nitrogen fixing microbes and ammonia oxidizers in grassland. v2. Biodiversity Exploratories Information System. Dataset. [WWW Document]. URL https://www.bexis.uni-jena.de/ddm/data/Showdata/13986
Stempfhuber, B., Schloter, M., 2016. Nitrogen Pools and Potential Nitrification taken at Soil Sampling Campaign 2014. v3. Biodiversity Exploratories Information System. Dataset. [WWW Document]. URL https://www.bexis.uni-jena.de/ddm/data/Showdata/19847
Takkis, K., 2014. Changes in plant species richness and population performance in response to habitat loss and fragmentation (Thesis).

Teuscher, M., Fischer, M., 2021. Bird survey and trait data on all grassland and forest EPs 2018. v3. Biodiversity Exploratories Information System. Dataset. [WWW Document]. URL https://www.bexis.uni-jena.de/ddm/data/Showdata/25306

Thuiller, W., n.d. Traits of European Alpine Flora - Wilfried Thuiller - OriginAlps Project.

Tobias, J.A., Sheard, C., Pigot, A.L., Devenish, A.J.M., Yang, J., Sayol, F., Neate-Clegg, M.H.C., Alioravainen, N., Weeks, T.L., Barber, R.A., Walkden, P.A., MacGregor, H.E.A., Jones, S.E.I., Vincent, C., Phillips, A.G., Marples, N.M., Montaño-Centellas, F.A., Leandro-Silva, V., Claramunt, S., Darski, B., Freeman, B.G., Bregman, T.P., Cooney, C.R., Hughes, E.C., Capp, E.J.R., Varley, Z.K., Friedman, N.R., Korntheuer, H., Corrales-Vargas, A., Trisos, C.H., Weeks, B.C., Hanz, D.M., Töpfer, T., Bravo, G.A., Remeš, V., Nowak, L., Carneiro, L.S., Moncada R., A.J., Matysioková, B., Baldassarre, D.T., Martínez-Salinas, A., Wolfe, J.D., Chapman, P.M., Daly, B.G., Sorensen, M.C., Neu, A., Ford, M.A., Mayhew, R.J., Fabio Silveira, L., Kelly, D.J., Annorbah, N.N.D., Pollock, H.S., Grabowska-Zhang, A.M., McEntee, J.P., Carlos T. Gonzalez, J., Meneses, C.G., Muñoz, M.C., Powell, L.L., Jamie, G.A., Matthews, T.J., Johnson, O., Brito, G.R.R., Zyskowski, K., Crates, R., Harvey, M.G., Jurado Zevallos, M., Hosner, P.A., Bradfer-Lawrence, T., Maley, J.M., Stiles, F.G., Lima, H.S., Provost, K.L., Chibesa, M., Mashao, M., Howard, J.T., Mlamba, E., Chua, M.A.H., Li, B., Gómez, M.I., García, N.C., Päckert, M., Fuchs, J., Ali, J.R., Derryberry, E.P., Carlson, M.L., Urriza, R.C., Brzeski, K.E., Prawiradilaga, D.M., Rayner, M.J., Miller, E.T., Bowie, R.C.K., Lafontaine, R.-M., Scofield, R.P., Lou, Y., Somarathna, L., Lepage, D., Illif, M., Neuschulz, E.L., Templin, M., Dehling, D.M., Cooper, J.C., Pauwels, O.S.G., Analuddin, K., Fjeldså, J., Seddon, N., Sweet, P.R., DeClerck, F.A.J., Naka, L.N., Brawn, J.D., Aleixo, A., Böhning-Gaese, K., Rahbek, C., Fritz, S.A., Thomas, G.H., Schleuning, M., 2022. AVONET: morphological, ecological and geographical data for all birds. Ecol. Lett. 25, 581–597. https://doi.org/10.1111/ele.13898

Tribouillois, H., Fort, F., Cruz, P., Charles, R., Flores, O., Garnier, E., Justes, E., 2015. A Functional Characterisation of a Wide Range of Cover Crop Species: Growth and Nitrogen Acquisition Rates, Leaf Traits and Ecological Strategies. PLOS ONE 10, e0122156. https://doi.org/10.1371/journal.pone.0122156

Tschapka, M., Renner, S., Jung, K., 2017a. Bird survey data 2008, all 300 EPs. v2. Biodiversity Exploratories Information System. Dataset. [WWW Document]. URL https://www.bexis.uni-jena.de/ddm/data/Showdata/21446

Tschapka, M., Renner, S., Jung, K., 2017b. Bird survey data 2009, all 300 EPs. v2. Biodiversity Exploratories Information System. Dataset [WWW Document]. URL https://www.bexis.uni-jena.de/ddm/data/Showdata/21447

Tschapka, M., Renner, S., Jung, K., 2017c. Bird survey data 2010, all 300 EPs. v2. Biodiversity Exploratories Information System. Dataset. [WWW Document]. URL https://www.bexis.uni-jena.de/ddm/data/Showdata/21448

Tschapka, M., Renner, S., Jung, K., 2017d. Bird survey data 2011, all 300 EPs. v2. Biodiversity Exploratories Information System. Dataset. [WWW Document]. URL https://www.bexis.uni-jena.de/ddm/data/Showdata/21449

VERGUTZ, L., MANZONI, S., PORPORATO, A., NOVAIS, R.F., JACKSON, R.B., 2012. A Global Database of Carbon and Nutrient Concentrations of Green and Senesced Leaves 0.22148 MB. https://doi.org/10.3334/ORNLDAAC/1106

Vile, D., 2005. Significations fonctionnelle et écologique des traits des espèces végétales exemple dans une succession post-culturale méditerranéenne et généralisations.

WALKER, A.P., 2014. A Global Data Set of Leaf Photosynthetic Rates, Leaf N and P, and Specific Leaf Area 1.103161 MB. https://doi.org/10.3334/ORNLDAAC/1224
Wang, H., Harrison, S.P., Prentice, I.C., Yang, Y., Bai, F., Furstenau Togashi, H., Wang, M., Zhou, S., Ni, J., 2017. The China Plant Trait Database. https://doi.org/10.1594/PANGAEA.871819
Weigelt, A., Mommer, L., Andraczek, K., Iversen, C.M., Bergmann, J., Bruelheide, H., Fan, Y., Freschet, G.T., Guerrero-Ramírez, N.R., Kattge, J., Kuyper, T.W., Laughlin, D.C., Meier, I.C., van der Plas, F., Poorter, H., Roumet, C., van Ruijven, J., Sabatini, F.M., Semchenko, M., Sweeney, C.J., Valverde-Barrantes, O.J., York, L.M., McCormack, M.L., 2021. An integrated framework of plant form and function: The belowground perspective. New Phytol. nph.17590. https://doi.org/10.1111/nph.17590
Weisser, W., Gossner, M., Pasalic, E., Lange, M., Türke, M., Gallenberger, I., Simons, N., Staab, M., 2019. Sweep net samples from grasslands since 2008: Araneae, Coleoptera, Hemiptera, Orthoptera. v4. Biodiversity Exploratories Information System. Dataset. [WWW Document]. URL https://www.bexis.uni-jena.de/ddm/data/Showdata/21969
Werner, G.D.A., Cornelissen, J.H.C., Cornwell, W.K., Soudzilovskaia, N.A., Kattge, J., West, S.A., Kiers, E.T., 2018. Symbiont switching and alternative resource acquisition strategies drive mutualism breakdown. Proc. Natl. Acad. Sci. 115, 5229–5234. https://doi.org/10.1073/pnas.1721629115
Westoby, M., Gillings, M.R., Madin, J.S., Nielsen, D.A., Paulsen, I.T., Tetu, S.G., 2021. Trait dimensions in bacteria and archaea compared to vascular plants. Ecol. Lett. ele.13742. https://doi.org/10.1111/ele.13742
Wilkinson, G.S., South, J.M., 2002. Life history, ecology and longevity in bats. Aging Cell 1, 124–131. https://doi.org/10.1046/j.1474-9728.2002.00020.x
Wilson, K.B., Baldocchi, D.D., Hanson, P.J., 2000. Spatial and seasonal variability of photosynthetic parameters and their relationship to leaf nitrogen in a deciduous forest. Tree Physiol. 20, 565–578. https://doi.org/10.1093/treephys/20.9.565
Wirth, C., Lichstein, J.W., 2009. The Imprint of Species Turnover on Old-Growth Forest Carbon Balances - Insights From a Trait-Based Model of Forest Dynamics, in: Wirth, C., Gleixner, G., Heimann, M. (Eds.), Old-Growth Forests: Function, Fate and Value, Ecological Studies. Springer, Berlin, Heidelberg, pp. 81–113. https://doi.org/10.1007/978-3-540-92706-8_5
Wolfgang Wagner, n.d. Europäische Schmetterlinge und ihre Ökologie [WWW Document]. URL http://www.pyrgus.de/ (accessed 2.7.22).
Wright, I.J., Reich, P.B., Westoby, M., Ackerly, D.D., Baruch, Z., Bongers, F., Cavender-Bares, J., Chapin, T., Cornelissen, J.H., Diemer, M., 2004. The worldwide leaf economics spectrum. Nature 428, 821–827.
Yguel, B., Bailey, R., Tosh, N.D., Vialatte, A., Vasseur, C., Vitrac, X., Jean, F., Prinzing, A., 2011. Phytophagy on phylogenetically isolated trees: why hosts should escape their relatives. Ecol. Lett. 14, 1117–1124. https://doi.org/10.1111/j.1461-0248.2011.01680.x
Zanne, A.E., Lopez-Gonzalez, G., Coomes, D.A., Ilic, J., Jansen, S., Lewis, S.L., Miller, R.B., Swenson, N.G., Wiemann, M.C., Chave, J., 2009. Global wood density database [WWW Document]. URL http://hdl.handle.net/10255/dryad.10235.
